# Tissue-like Viscoelasticity Drives Muscle Stem Cell Expansion Through Mechanical Memory Formation

**DOI:** 10.64898/2026.11.06.764681

**Authors:** Haiying Hu, Tae Yoon Kwon, Zichen Gao, Ping Zhou, Christopher M Madl

## Abstract

Mechanical cues are potent regulators of stem cell fate, with even transient exposure to substrates with varied elastic stiffness driving persistent changes in cell fate. Such elastic mechanical memory contributes to the loss of regenerative potential in skeletal muscle stem cells (MuSCs) when cultured on rigid substrates, hampering the translation of myogenic cell replacement therapies. Beyond elastic mechanical properties, cells also respond to time-dependent, viscoelastic mechanical properties, such as stress relaxation. However, whether tissue-like viscoelasticity can impart a mechanical memory that directs stem cell fate remains an open question. Accordingly, dual-network hydrogel biomaterials with decoupled control of stiffness, stress relaxation, and adhesive ligand presentation were engineered to recapitulate the viscoelastic mechanical properties of native muscle. Muscle-like viscoelastic substrates promote in vitro MuSC activation and accelerate stem cell expansion, matching in vivo activation timescales. Mechanistically, physiological viscoelasticity promotes focal adhesion stabilization and enhances Rac-Akt signaling, establishing a viscoelastic mechanical memory that supports progenitor expansion and persistent maintenance of a stem-like phenotype. These findings identify matrix viscoelasticity as a potent and persistent regulator of MuSC fate and provide a framework for engineering biomaterials that preserve MuSC stemness and expansion ex vivo.

## Introduction

Adult stem cells are responsible for maintaining tissue function throughout life and regenerating tissues following injury.^[1]^ The maintenance, expansion, and differentiation of stem cells during tissue repair is orchestrated by a complex interplay of biochemical and mechanical factors from the extracellular microenvironment.^[2,3]^ Previous work has demonstrated that the elastic mechanical properties, or stiffness, of the microenvironment are potent regulators of stem cell fate.^[4–7]^ Stem cells, such as mesenchymal stromal cells and skeletal muscle stem cells (MuSCs) “remember” transient exposure to microenvironments of varied stiffness, forming mechanical memories that bias subsequent fate determination.^[8–10]^

The formation of stem cell mechanical memory is an essential consideration in the development of stem cell-based therapies. Many proposed stem cell therapies require ex vivo expansion and/or in vitro gene editing of isolated stem cells, exposing the cells to engineered environments with dramatically different properties compared to the native stem cell niche.^[11]^ In the case of MuSCs, earlier attempts to develop cell therapies for heritable muscle wasting disorders like Duchenne muscular dystrophy (DMD) failed due to the inability of transplanted cells to durably engraft as functional stem cells.^[12–14]^ Later work demonstrated that MuSCs are acutely sensitive to microenvironmental stiffness, rapidly losing engraftment potential when cultured on rigid tissue culture plastic.^[15]^ Culture on hydrogel substrates with muscle like stiffness partially restored engraftment potential.^[7]^ We recently demonstrated that MuSCs acquire a mechanical memory during activation ex vivo that impairs the expansion of functional progenitors, helping explain why culture on substrates with muscle-like stiffness improves engraftment.^[10]^

However, matching microenvironmental stiffness only partially rescues expanded MuSC function, with the expanded cells unable to achieve the engraftment potential of freshly isolated cells.^[7]^ We postulate that this decreased function is due to limitations of existing hydrogel systems that exhibit predominantly elastic material properties. Muscle tissue is highly viscoelastic, as its macromolecular networks reorganize and dissipate applied stresses over time.^[16–18]^ Thus, in addition to stiffness, the time-dependent viscoelastic mechanical properties of muscle tissue may provide important cues that regulate MuSC behavior. Stem cells, including MuSCs, respond to viscoelastic mechanical cues,^[16,19–24]^ suggesting tissue-like viscoelasticity is a crucial material design consideration for stem cell expansion platforms. However, whether adult stem cells form memories of microenvironmental viscoelasticity remains unknown. For MuSCs, we anticipated that matching both the stiffness and viscoelastic stress relaxation timescale of healthy muscle tissue would drive the expansion of functional progenitor cells, opening the door to cell-based therapies for diseases like DMD when gene editing and exon skipping approaches fall short.^[25–28]^

Existing hydrogel systems for MuSC culture fail to simultaneously match both the elastic stiffness and viscoelastic stress relaxation time of muscle.^[7,10,23,24]^ We thus developed a synthetic hydrogel platform that could stably present cultured cells with muscle-like stiffness and stress relaxation time, while enabling consistent presentation of cell adhesive cues, during extended culture in vitro. We developed dual network hydrogels that combined the dynamic nature of hydrazone crosslinks with the stability of strain-promoted azide-alkyne cycloaddition (SPAAC) crosslinks to achieve both physiological viscoelasticity and long-term mechanical stability in a single platform. These dual-network hydrogels revealed that viscoelastic substrates engage a focal adhesion-mediated Rac-PAK-Akt signaling axis, uncovering a previously unrecognized mechanism of viscoelastic mechanosensing in MuSCs. Muscle-like viscoelasticity promotes earlier MuSC activation and proliferation by modulating the balance of Rac1/Cdc42:RhoA signaling. Furthermore, we identified a persistent viscoelastic mechanical memory mediated by Rac-PAK-Akt signaling, which contributes to the preservation of MuSC stemness during prolonged culture. Collectively, these findings demonstrate that matrix stiffness and stress relaxation jointly regulate MuSC activation, mechanosensing, proliferation and fate determination, providing new insights into the mechanobiology of muscle regeneration and establishing matrix viscoelasticity as a key design parameter for maintaining stem cell potency during ex vivo expansion.

### Hydrogels with physiologic viscoelasticity enhance MuSC expansion

Native mouse skeletal muscle exhibits a storage modulus of approximately 4 kPa (elastic modulus ∼ 12 kPa) ^[7]^ and a stress relaxation half-time (τ_1/2_) of ∼10^2^ s (**Fig. 1A**). To recapitulate these mechanical properties, we developed a double-network hydrogel platform with independent tuning of stiffness and stress relaxation time using multi-arm poly(ethylene glycol) (PEG) macromers functionalized with aldehyde and hydrazine moieties.^[29,30]^ These two macromers assemble into a dynamic, reversible hydrazone-crosslinked network, enabling stress relaxation. A second, elastic, SPAAC-based PEG network was incorporated to stabilize the dynamic viscoelastic network and reinforce hydrogel structure ^[31]^ (**Fig. 1B,C**). The chemical structure and reaction kinetics of the SPAAC network were important design considerations. Incorporation of a hydrophobic, bulky SPAAC crosslinker can reduce hydrogel swelling and thereby limit network erosion and degradation of the dynamic hydrazone network. In addition, the kinetics of the SPAAC reaction can influence the connectivity of the reinforcing network and, consequently, the extent to which the hydrazone network can dissipate stress. We therefore selected dibenzocyclooctyne (DBCO) based SPAAC chemistry for the initial hydrogel formulations. For stiffer, fibrotic-like hydrogels with a storage modulus of ∼13.3 kPa, higher polymer concentrations were required, which accelerated network formation and increased the formation of defects within the SPAAC network. To mitigate this effect and achieve sufficient reinforcement of the stiffer hydrogels, we incorporated cyclooctye (CO) based SPAAC chemistry, which exhibits slower reaction kinetics than DBCO-based crosslinking. This slower reaction promotes a more controlled formation of the reinforcing network while preserving the dynamic stress relaxation behavior imparted by the hydrazone network. To assess relaxation times faster than muscle tissue, we employed boronic ester crosslinks formed from multi-arm PEG precursors functionalized with nitrodopamine and arylboronic acids to enable relaxation on the order of 10^-1^ s ^[32–34]^ (**Figs. 1B, S1A,B**). To prevent rapid gel erosion, a similar strategy using SPAAC-based reinforcement was applied to improve hydrogel stability.^[34]^ Purely SPAAC-based networks with irreversible crosslinks were used as predominantly elastic controls.

**Fig. 1.**
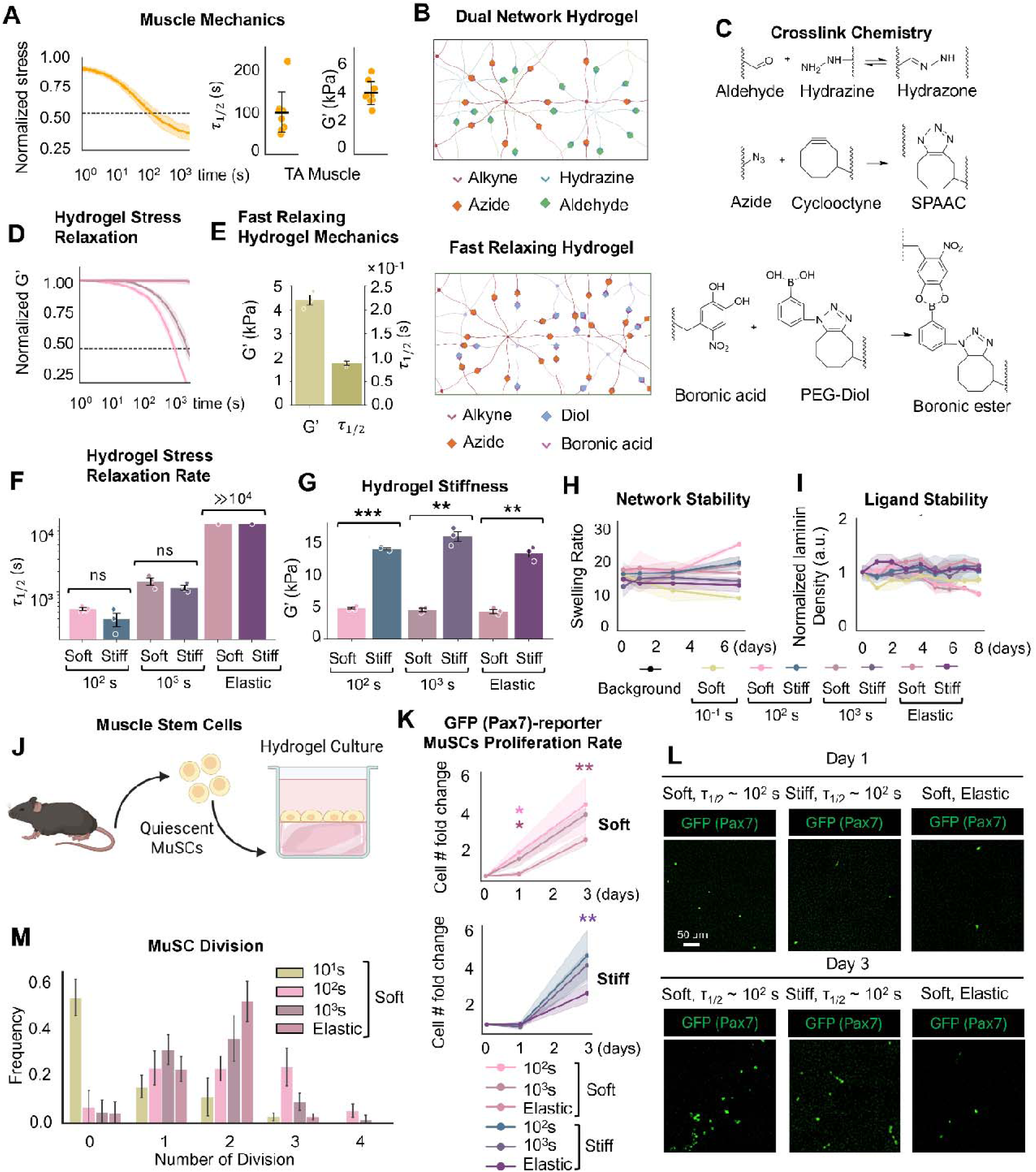
A tunable viscoelastic hydrogel platform recapitulates muscle mechanics and accelerates MuSC expansion in vitro. (**A**) Mechanical properties of native mouse tibialis anterior (TA) muscle, showing storage modulus (*G*′) of ∼4 kPa and a physiological stress relaxation half-time (τ_1/2_) of ∼10² s. (**B**) Schematics of the PEG hydrogel network architecture used to generate hydrogels with distinct stress relaxation dynamics, including the dual-network hydrazone-SPAAC hydrogel and the fast-relaxing hydrogel incorporating boronate ester crosslinks. (**C**) Chemical schemes illustrating the hydrazone, SPAAC, and boronate ester crosslinking chemistries used to control hydrogel stress relaxation and network stability. (**D**) Stress relaxation behavior of PEG hydrogels with different relaxation dynamics. (**E**) Fast-relaxing PEG hydrogels exhibit a storage modulus of ∼4 kPa and a stress relaxation half-time of ∼10^-1^ s. (**F**) Stress relaxation half-times (τ_1/2_) of soft and stiff hydrogels with target relaxation times of ∼10^2^, 10^3^, and ≫10^4^ s (elastic). (**G**) Storage modulus (*G*′) of soft and stiff hydrogels across different stress relaxation conditions. PEG hydrogels maintain stable (**H**) swelling and (**I**) laminin presentation across mechanical conditions. (**J**) Schematics illustrating the culture protocols for freshly isolated mouse MuSCs on gels. (**K**) Quantification of cell number fold change shows more Pax7^+^ progenitors on soft, stress-relaxing hydrogels by day 1. (**L**) Representative images of Pax7-EGFP reporter MuSCs cultured on soft and stiff hydrogels with physiological stress relaxation (τ_1/2_∼10^2^ s) and soft, elastic hydrogels at days 1 and 3 after isolation. (**M**) Frequency distribution of MuSC division number from day 2 to day 3 under different stress relaxation conditions. Data are presented as mean ± s.d. from *n* ≥ 3 independent replicates for hydrogel characterization and from *n* ≥ 300 cells analyzed per hydrogel condition for single cell measurements. Statistical significance was determined using pairwise Welch’s *t*-tests with multiple-comparison correction. *P* < *0.05 (*), P < 0.01 (\***\***), P **<** 0.001 (***)*. In (**J**), the color of the significance stars indicates the corresponding mechanical condition compared with the elastic control.

The stress relaxation times of the hydrazone-crosslinked hydrogels were tuned by varying network connectivity, with networks formed from macromers of higher functionality exhibiting slower stress relaxation.^[22,29]^ By varying the architecture and concentrations of the macromers, we prepared hydrogels spanning physiologically relevant stiffnesses (storage modulus, *G’* ∼ 4 kPa, soft, representative of healthy muscle, and *G’* ∼ 13 kPa, stiff, representative of fibrotic muscle ^[35,36]^) and stress relaxation timescales (τ_1/2_ ∼10^-1^, 10^2^, 10^3^, and ≫10^4^ s) (**Figs. 1D-G, S1C-G**), as characterized by oscillatory rheology. This hydrogel platform enables independent control of stiffness and viscous dissipation, allowing physiologically relevant biophysical features of the MuSC niche to be recapitulated *ex vivo*. The resulting hydrogels remain mechanically stable, exhibit negligible swelling during culture, and maintain consistent laminin presentation over time to support cell attachment and spreading (**Figs. 1H,I, S1H**).

Following muscle injury in vivo, MuSCs activate and begin dividing within 24 hours, expanding 4- to 12-fold within the first 3 days.^[37,38]^ We recently reported that MuSCs cultured in vitro form a mechanical memory that biased subsequent cell fate decisions based on the substrate stiffness experienced during the first 3 days post-isolation as the cells exited quiescence and activated.^[10]^ Accordingly, we asked whether ECM viscoelasticity similarly primed MuSC fate commitment during activation, examining MuSC responses to substrates with varying stiffness and stress relaxation behavior at days 1 and 3 post-isolation.

Using reporter MuSCs expressing EGFP in response to expression of the stemness marker Pax7,^[39]^ the number of Pax7-expressing progenitors at early time points was quantified following culture on hydrogels with defined mechanical properties. By day 1, a greater number of MuSCs cultured on soft, viscoelastic hydrogels were observed, compared to cells cultured on soft, elastic controls (**Figs. 1J,K, S2A**). No significant differences in cell number were observed between viscoelastic and elastic hydrogels of fibrotic-like stiffness. Notably, cell division on soft, physiologically relaxing hydrogels was already detectable by day 1 (**Fig. S2B-E, Supporting Movies S1-3**), consistent with the onset of cell division observed in vivo.^[37]^ Thus, physiological viscoelasticity supports native-like activation kinetics not observed in purely elastic hydrogels.^[7]^ Furthermore, analysis of subsequent cell division kinetics revealed that MuSCs cultured on hydrogels with physiologically relevant stress relaxation rates proliferated more rapidly than those cultured on rapidly relaxing or predominantly elastic matrices (**Fig. 1L**). In vivo, activated MuSCs exhibit an average cell-cycle duration of approximately 8 hrs.^[40]^ Consistent with this timescale, physiologically relaxing hydrogels supported a higher frequency of MuSCs undergoing three divisions within 24 h, indicating that muscle-like viscoelasticity more closely recapitulates the cell-cycle dynamics of activated MuSCs in vivo. Collectively, these findings demonstrate that physiological substrate viscoelasticity promotes MuSC proliferation at early time points post-isolation.

### Substrate viscoelasticity promotes MuSC activation

To validate that the accelerated MuSC expansion observed on physiologically relaxing substrates reflects earlier activation, freshly isolated MuSCs were seeded on the laminin-functionalized hydrogels. After 1 day of culture, cells were fixed and analyzed by immunofluorescence staining for activation-associated markers, including 5-ethynyl-2′-deoxyuridine (EdU) incorporation,^[10]^ Ki67 expression,^[41]^ p38 mitogen-activated protein kinase (MAPK) activity,^[10,42]^ expression of the Polycomb repressive complex 2 (PRC2) catalytic subunit EZH2,^[43]^ and the subcellular localization of the forkhead transcription factors FOXO3a and FOXO1.^[44–46]^

At day 1 post-isolation, MuSC proliferation and activation exhibited a biphasic dependence on matrix viscoelasticity. MuSCs cultured on soft hydrogels with physiologically relevant stress relaxation rates displayed the greatest degree of activation, as evidenced by increased fractions of EdU□(**Figs. 2A, S3**) and Ki67 cells (**Figs. 2B**, **S3**), together with elevated levels of phosphorylated p38 MAPK (**Fig. 2C**) and EZH2 expression (**Figs. 2D**, **S3**). Nuclear-to-cytoplasmic translocation of FOXO proteins is a hallmark of MuSC activation and exit from quiescence.^[44,45]^ Notably, FOXO3a localization was strongly regulated by matrix viscoelasticity, with faster stress relaxing hydrogels promoting greater cytoplasmic accumulation of FOXO3a (**Figs. 2E**, **S4A**). Matrix viscoelasticity also modestly enhanced the cytoplasmic translocation of FOXO1 (**Fig. S4B-D**). These findings indicate that physiological viscoelasticity promotes MuSC exit from quiescence and entry into the cell cycle, thereby accelerating activation and proliferation.

**Fig. 2.**
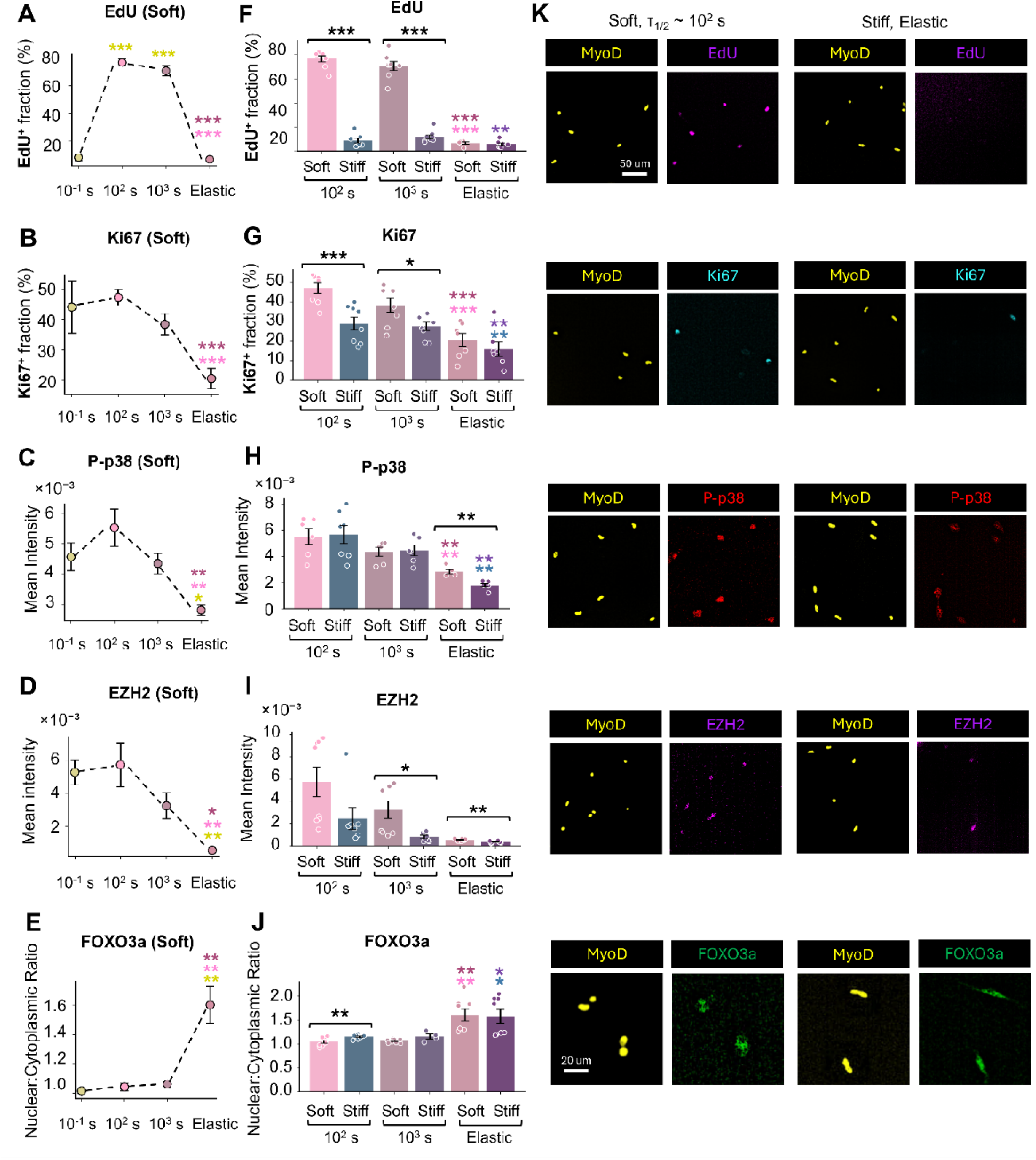
Physiological viscoelasticity promotes MuSC activation. Quantification of (**A** and **F**) EdU incorporation, (**B** and **G**) Ki67^+^ fraction, (**C** and **H**) phosphorylated p38 (p-p38) intensity, (**D** and **I**) EZH2 expression, and (**E** and **J**) the nuclear-to-cytoplasmic ratio of FOXO3a in MuSCs cultured on hydrogels with varying stress relaxation rates (τ_1/2_ ∼10^-1^, 10^2^, 10^3^, and ≫10^4^ s) under soft conditions (**A**, **B**, **C**, **D**, **E**), or varying stiffnesses (soft and stiff; τ_1/2_ ∼10^2^, 10^3^, and ≫10^4^ s) (**F**, **G**, **H**, **I**, **J**). (**K**) Representative immunofluorescence images of MuSCs stained for MyoD together with EdU, Ki67, p-p38, EZH2, or FOXO3a. Each data point represents the fraction of positive cells (**A**, **B**, **F**, **G**), mean fluorescence intensity (**C**, **D**, **H**, **I**), or mean nuclear-to-cytoplasmic translocation ratio (**E**, **J**) calculated from N ≥ 300 cells per hydrogel condition. Data are presented as mean ± s.d. from *n* ≥ 4 independent biological replicates. Statistical significance was determined using pairwise Welch’s *t*-tests with multiple-comparison correction. *P* < 0.05 (\**), P < 0.01 (**), and P < 0.001 (\*\*\**). In (**A**-**H**), the color of the significance stars indicates the corresponding mechanical condition compared with the elastic control.

Substrate stiffness further synergized with stress relaxation to influence activation dynamics. Increased EdU incorporation (**Fig. 2F**) and Ki67 expression (**Fig. 2G**) were observed specifically in stress relaxing hydrogels with muscle-like stiffness, whereas MuSC activation remained limited on purely elastic substrates. In contrast, substrate stiffness more broadly influenced activation-associated signaling, with MuSCs cultured on soft hydrogels exhibiting elevated EZH2 expression relative to stiff hydrogels (**Fig. 2I**). Substrate stiffness did not significantly affect FOXO3 translocation, whereas physiologically relaxing matrices promoted greater FOXO3a nuclear-to-cytoplasmic translocation across stiffness conditions. Collectively, these results indicate that stiffness and viscous dissipation cooperatively regulate MuSC activation, with physiologically relevant stress relaxation enabling robust cell-cycle entry on compliant matrices.

By day 3 post-isolation, differences in proliferation among the mechanical conditions became less pronounced. Although MuSCs cultured on substrates with muscle-like viscoelasticity continued to exhibit slightly higher fractions of EdU^+^ (**Fig. S5A**) and Ki67^+^ cells (**Fig. S5B**), more than 70% of MuSCs were positive for proliferation markers across all mechanical conditions, indicating that the majority of cells had activated by this time point. The levels of phosphorylated p38 MAPK and EZH2 remained comparable to those observed at day 1, while differences between mechanical conditions were still evident, with physiologically relevant viscoelasticity associated with higher levels of both markers (**Fig. S5C,D**). Physiological viscoelasticity continued to promote FOXO3a nuclear-to-cytoplasmic translocation at day 3, although this effect was less pronounced than at day 1. MuSCs cultured on elastic substrates exhibited reduced FOXO3a nuclear-to-cytoplasmic translocation compared with day 1, whereas cells cultured on viscoelastic substrates maintained FOXO3a cytoplasmic localization at levels comparable to those observed at day 1 (**Fig. S6A-C**). In contrast, FOXO1 nuclear-to-cytoplasmic translocation was generally reduced at day 3 compared with day 1. However, physiologically relaxing substrates continued to promote greater cytoplasmic localization of FOXO1, consistent with the pattern observed for FOXO3a (**Fig. S6D-F**). These findings suggest that while the differences in proliferation diminish as MuSCs progress through activation, substrate viscoelasticity continues to influence activation-related signaling.

### Matrix viscoelasticity regulates Rac1/Cdc42-to-Rho balance and Akt activity

To determine the mechanosensitive signaling pathways regulated by substrate viscoelasticity, we first examined the subcellular localization of Yes-associated protein 1 (YAP), as MuSCs are known to respond to substrate stiffness through YAP/TAZ-mediated mechanotransduction.^[36]^ Furthermore, YAP/TAZ transcriptional coactivators have been proposed to function as intracellular mechanical rheostats that encode mechanical memory in stem cells.^[8,47]^ However, we observed no significant differences in YAP localization across hydrogels with different stress relaxation rates. Instead, increased nuclear YAP localization was primarily associated with stiffer hydrogels (**Fig. S7**). These findings are insufficient to explain the complex MuSC behaviors elicited by viscoelastic matrices, suggesting that YAP signaling predominantly reflects sensing of stiffness rather than viscous dissipation. Therefore, MuSCs likely sense matrix stiffness and viscous dissipation through distinct mechanotransduction mechanisms, indicating that the effects of viscoelasticity on cell fate are not solely mediated by canonical YAP/TAZ signaling pathways and may instead involve alternative regulators of cytoskeletal remodeling and force transmission.

We next examined the activity of Rho GTPases in MuSCs using in situ binding assays,^[48]^ as this family of GTPases has been implicated in maintenance of quiescence and MuSC activation in mouse models.^[48–50]^ At day 1, hydrogels with faster stress relaxation rates increased the activities of both Rac/Cdc42 and Rho (Fig. 3, A to F). However, the balance between Rac/Cdc42 and Rho activity exhibits a biphasic trend, with hydrogels with physiological stress relaxation resulting in the highest Rac/Cdc42:Rho activity ratio (**Figs. 3G**, **S8**). With respect to stiffness, soft viscoelastic hydrogels enhanced both Rac/Cdc42 and Rho activities compared with stiff viscoelastic hydrogels (**Figs. 3D,F**, **S8**), but the Rac/Cdc42:Rho activity ratio was not significantly altered between stiffness conditions (**Figs. 3H**, **S8**). At day 3, hydrogels exhibiting muscle-like stiffness (*G’* ∼ 4 kPa) and physiological stress relaxation behavior (τ_1/2_ ∼ 10^2^ s) continued to exhibit the highest Rac/Cdc42:Rho activity ratio. However, the effects of viscoelasticity on Rac and Rho signaling became less pronounced at this later activation stage (**Fig. S9**), suggesting that viscoelastic regulation of Rac-Rho signaling is most prominent during the early phase of MuSC activation.

**Fig. 3.**
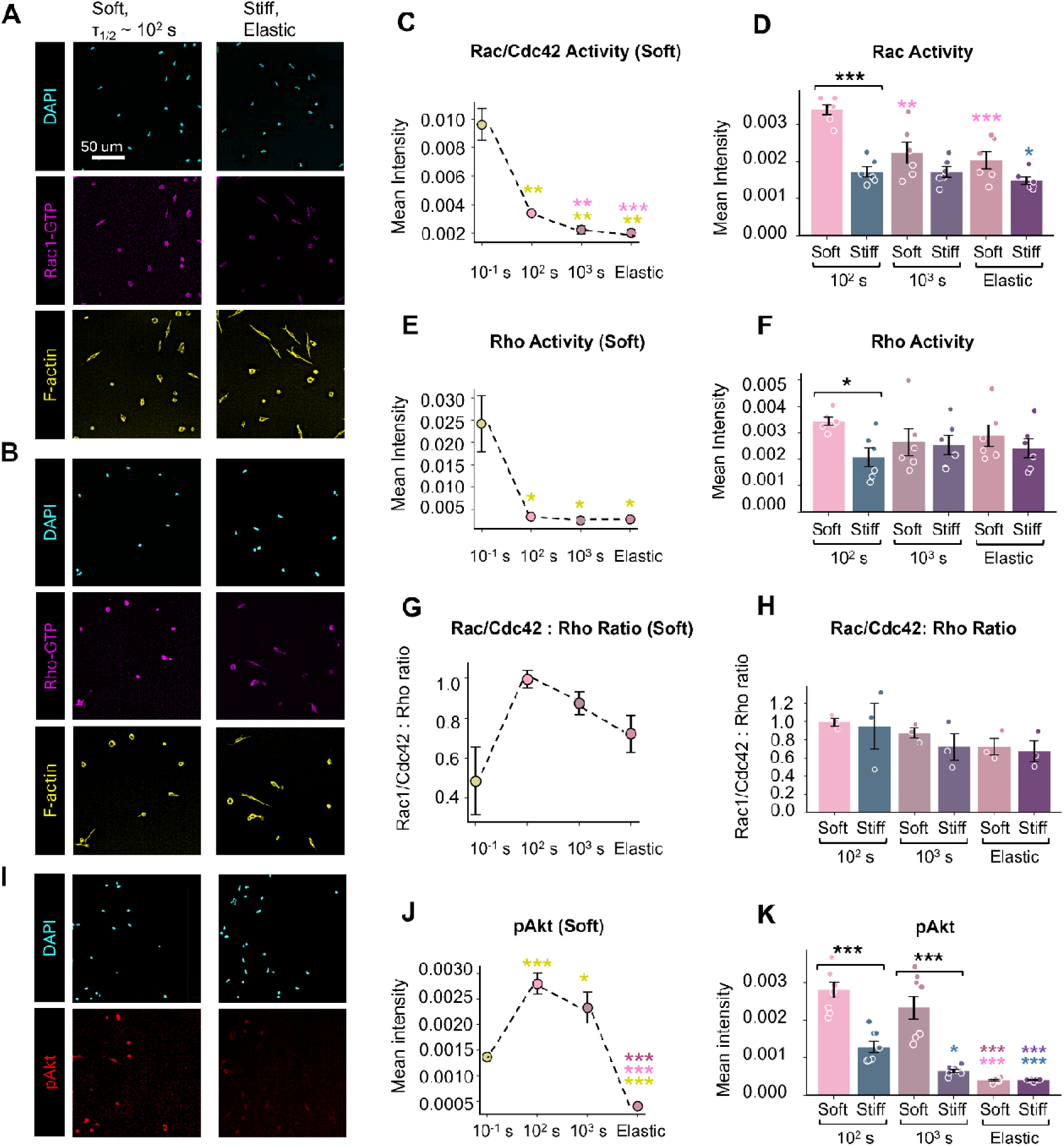
Matrix viscoelasticity regulates Rac/Cdc42-Rho signaling balance and Akt activation. Representative immunofluorescence images of (**A**) active Rac/Cdc42 and (**B**) active Rho in MuSCs cultured on soft hydrogels with physiological stress relaxation (τ_1/2_ ∼ 10^2^ s) and stiff elastic hydrogels. Quantification of (**C** and **D**) Rac/Cdc42 activity, (**E** and **F**) Rho activity, and (**G** and **H**) the Rac/Cdc42:Rho activity ratio in MuSCs cultured on hydrogels with varying stress relaxation rates under soft conditions (**C**, **E**, **G**) or across soft and stiff hydrogels (**D**, **F**, **H**). (**I**) Representative immunofluorescence images of pAkt in MuSCs cultured on soft hydrogels with physiological stress relaxation (τ_1/2_ ∼ 10^2^ s) and stiff elastic hydrogels. Quantification of pAkt staining in MuSCs cultured on hydrogels with varying stress relaxation rates (**J**) under soft conditions or (**K**) across soft and stiff hydrogels. Each data point represents mean fluorescence intensity (**C**-**F**, **J**, **K**) calculated from N ≥ 200 cells per hydrogel condition. Data are presented as mean ± s.d. from *n* ≥ 3 independent biological replicates. Statistical significance was determined using linear mixed-effects models (LMMs), with mechanical condition as a fixed effect and independent experiments as a random effect. Pairwise comparisons were performed with multiple-comparison correction. *P < 0.05 (*), P < 0.01 (**), and P < 0.001 (***)*. In (**C**-**H**) and (**J**-**K**), the color of the significance stars indicates the corresponding mechanical condition compared with the elastic control.

To investigate how the altered Rac/Cdc42:Rho balance may influence cell fate, we subsequently investigated the activity of protein kinase B (Akt). Because Akt acts downstream of Rac signaling and partially antagonizes Rho-mediated pathways, it can serve as an important integrator of microenvironmental cues that regulate MuSC activation and survival.^[51,52]^ Measuring Akt activity by immunostaining for phosphorylated (active) Akt (pAkt) revealed a biphasic relationship for Akt activity as a function of stress relaxation time, following a similar trend to that observed for the Rac/Cdc42:Rho activity ratio (**Fig. 3I,J**). Two-way ANOVA demonstrated that substrate stiffness and stress relaxation behavior synergistically regulate Akt signaling activity, with optimal activation occurring on substrates with physiological viscoelasticity (**Fig. 3I-K**). Hydrogels that recapitulate both physiological stiffness and stress relaxation behavior also exhibited enhanced Akt signaling at day 3 post-isolation (**Fig. S10**). These findings indicate that physiological viscoelasticity promotes Akt activation, providing a potential mechanism by which viscoelastic mechanosensing regulates MuSC activation and expansion.^[52–54]^

### Focal adhesion signaling mediates viscoelastic mechanosensing regulating MuSCs activation

To determine the mechanism by which viscoelasticity regulates MuSC activation, we first examined cell morphology and motility during culture on the hydrogel substrates. Live-cell imaging revealed that cells maintained a more static morphology on soft hydrogels with physiological stress relaxation (τ_1/2_ ∼ 10^2^ s). This phenotype was characterized by a smaller cell spread area (**Fig. 4A**), reduced variability in cell shape, as indicated by a lower standard deviation of the aspect ratio (**Fig. 4B, and D**), and a slower average migration speed (**Fig. 4C**). Together, these observations suggest that cell-matrix adhesion dynamics are regulated by viscous dissipation, leading us to investigate focal adhesion signaling as a potential mechanism underlying viscoelastic mechanosensing.

**Fig. 4.**
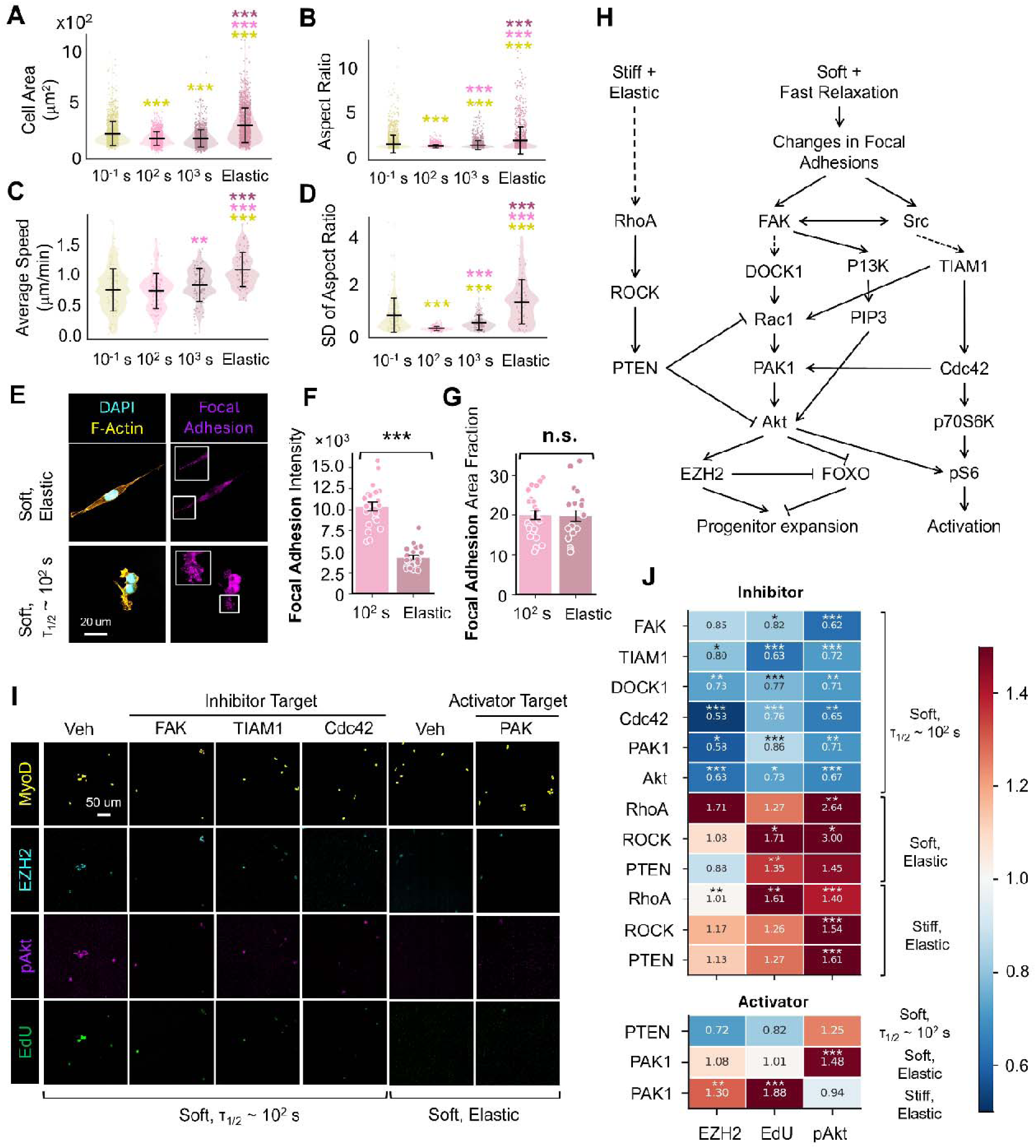
Focal adhesion-mediated mechanotransduction links substrate viscoelasticity to MuSC activation via Rac-Akt signaling. Quantification of **(A)** cell spread area, **(B)** aspect ratio, (**C**) average migration speed, and (**D**) the standard deviation of aspect ratio in MuSCs cultured on soft hydrogels with varying stress relaxation rates from days 2 to 3 post-isolation. (**E**) Representative immunofluorescence images of F-actin and focal adhesions in MuSCs cultured on soft elastic hydrogels and soft hydrogels with physiological stress relaxation (τ_1/2_ ∼ 10^2^ s). Quantification of (**F**) focal adhesion intensity and (**G**) focal adhesion area fraction. (**H**) Proposed model illustrating how MuSCs transduce physiological viscoelasticity. (**I**) Representative immunofluorescence images of MuSCs stained for MyoD, EZH2, pAkt, and EdU following small molecule inhibition of key signaling components. (**J**) Heat maps summarizing changes in EZH2 and pAkt expression, cell proliferation (EdU^+^), following inhibition or activation of mechanotransduction pathways under different mechanical conditions. Data in (**A**-**D**, **F**, and **G**) are presented as mean ± s.d. from *n* ≥ 20 individual cells and were compared using two-sided Welch’s *t*-tests. Data in (**J**) are presented as mean ± s.d. from *n* ≥ 4 independent biological replicates and were analyzed using linear mixed-effects models (LMMs), with treatment condition included as a fixed effect and independent muscle batches included as a random effect. Detailed results are presented in the Supporting Information (fig. S11 to S14). *P < 0.05 (*), P < 0.01 (**), and P < 0.001 (***)*. In (**A**-**D**), the color of the significance stars indicates the corresponding mechanical condition compared with the elastic control. In (**J**), significance stars indicate comparisons between each pharmacological perturbation and the corresponding vehicle control.

Accordingly, we next examined focal adhesion assembly at the cell-matrix interface by immunostaining for paxillin and quantitatively analyzing focal adhesion properties using the Focal Adhesion Analysis Server.^[55]^ MuSCs cultured on hydrogels with physiological stress relaxation (τ_1/2_ ∼ 10^2^ s) exhibited more intense focal adhesions compared with elastic substrates (**Fig. 4F**), while no significant differences in focal adhesion area fraction were observed across mechanical conditions (**Fig. 4G**). These findings suggest that viscous dissipation regulates the maturation and stability of focal adhesions rather than their overall abundance.

Based on these observations, we propose that MuSCs sense matrix viscoelasticity through focal adhesion-mediated mechanotransduction, which subsequently regulates downstream Rac/Cdc42 and Akt signaling pathways (**Fig. 4H**). Specifically, soft substrates with physiological stress relaxation behavior promote the formation of more stable focal adhesions, leading to the activation of focal adhesion kinase (FAK) and protein-tyrosine kinase Src signaling. FAK activation then initiates a signaling cascade involving the Rho-guanine nucleotide exchange factor (GEF) DOCK1, Rac1, and p21-activated kinase (PAK), which enhances Akt activation and, in turn, promotes progenitor expansion through increased EZH2 expression and cytoplasmic translocation of FOXO transcription factors.^[45,56–58]^ In parallel, FAK and Src promote Cdc42 activation via the GEF TIAM1, reinforcing the pro-activation and expansion signal.^[59]^ In contrast, stiff and elastic matrices favor RhoA-ROCK signaling, resulting in increased expression of phosphatase and tensin homolog (PTEN) and suppression of Akt signaling, thereby limiting progenitor expansion.^[45,60]^

To test this mechanistic model, we performed pharmacological perturbation studies targeting key components of the proposed signaling network. Freshly isolated MuSCs were seeded onto soft hydrogels with physiological stress relaxation (τ_1/2_ ∼ 10^2^ s) and separately treated for 24 h with inhibitors of FAK (Defactinib), Akt (Akt inhibitor VIII), TIAM1 (NSC23766), DOCK1 (TBOPP), Cdc42 (ML-141) and PAK1 (NVS-PAK1-1) to disrupt signaling pathways proposed to be activated by physiological viscous dissipation. To test whether signaling associated with predominantly elastic matrices could override viscous sensing, MuSCs cultured on viscoelastic hydrogels were also treated with a PTEN activator (Tyroserleutide hydrochloride). Alternatively, MuSCs cultured on elastic substrates were separately treated for 24 h with inhibitors of RhoA (Rhosin), ROCK (Y-27632), and PTEN (bpV(HOpic)), as well as a PAK1 activator (Fingolimod), to determine whether suppressing RhoA-ROCK signaling and activating the Rac-PAK-Akt axis could recapitulate the effects of physiological viscous dissipation. Following treatment, cells were fixed and analyzed for EdU incorporation, pAkt, and EZH2 expression.

Our findings indicate that pharmacological inhibition of FAK, the Rac-PAK-Akt axis, and TIAM1/Cdc42 signaling markedly reduced Akt activity and EZH2 expression, as well as MuSC proliferative fraction (**Figs. 4J, S11, S12**), suggesting that these pathways are critical mediators of early MuSC activation on viscoelastic matrices. Conversely, suppression of RhoA-ROCK signaling on elastic substrates increased pAkt and EZH2 expression and modestly enhanced the EdU fraction (**Fig. 4J**), indicating that inhibiting signaling associated with elastic matrices can partially promote progenitor activation. Interestingly, activation of PTEN on soft, stress relaxing hydrogels had no significant effect on MuSC behavior, suggesting that transient (24 h) pharmacological perturbation of this single factor is insufficient to override ECM-derived mechanical cues that regulate MuSC fate decisions. Similarly, activation of PAK1 on elastic hydrogels increased EZH2 expression and cell proliferation on stiff substrates and enhanced Akt activity on soft substrates but was insufficient to simultaneously rescue all three readouts (**Fig. 4J**). Together, these results suggest that focal adhesion-mediated mechanotransduction regulates MuSC fate through coordinated Rho, Rac and Cdc42 GTPase and Akt signaling, and that no single downstream signaling component is sufficient to fully recapitulate the effects of physiological viscoelasticity.

### Viscoelasticity regulates MuSC fate determination

Prior work has demonstrated that substrate stiffness drives persistent changes in MuSC function after 7 days in culture, with excessively stiff substrates limiting progenitor expansion, increasing commitment to differentiation, and impairing in vivo engraftment.^[7,10]^ To assess whether substrate viscoelasticity similarly affects the longer-term fate determination in expanded cells, freshly isolated MuSCs were seeded on the laminin-functionalized hydrogels. After 7 days of culture, cells were fixed and analyzed by immunofluorescence staining for the myogenic markers Pax7, MyoD, myogenin, and skeletal muscle actin (Acta1). The fraction of cells retaining proliferative potential at day 7 was quantified using EdU incorporation.^[10]^ Hydrogels exhibiting both muscle-like stiffness (*G’* ∼ 4 kPa) and stress relaxation behavior (τ_1/2_ ∼ 10^2^ s) yielded the highest fractions of Pax7^+^ and EdU^+^ cells and the lowest fraction of Acta1^+^ cells across all mechanical conditions tested (**Fig. 5**). These results indicate that physiologically relevant matrix mechanics best preserve MuSC stemness and proliferative capacity while suppressing differentiation at 7 days post-isolation.

**Fig. 5.**
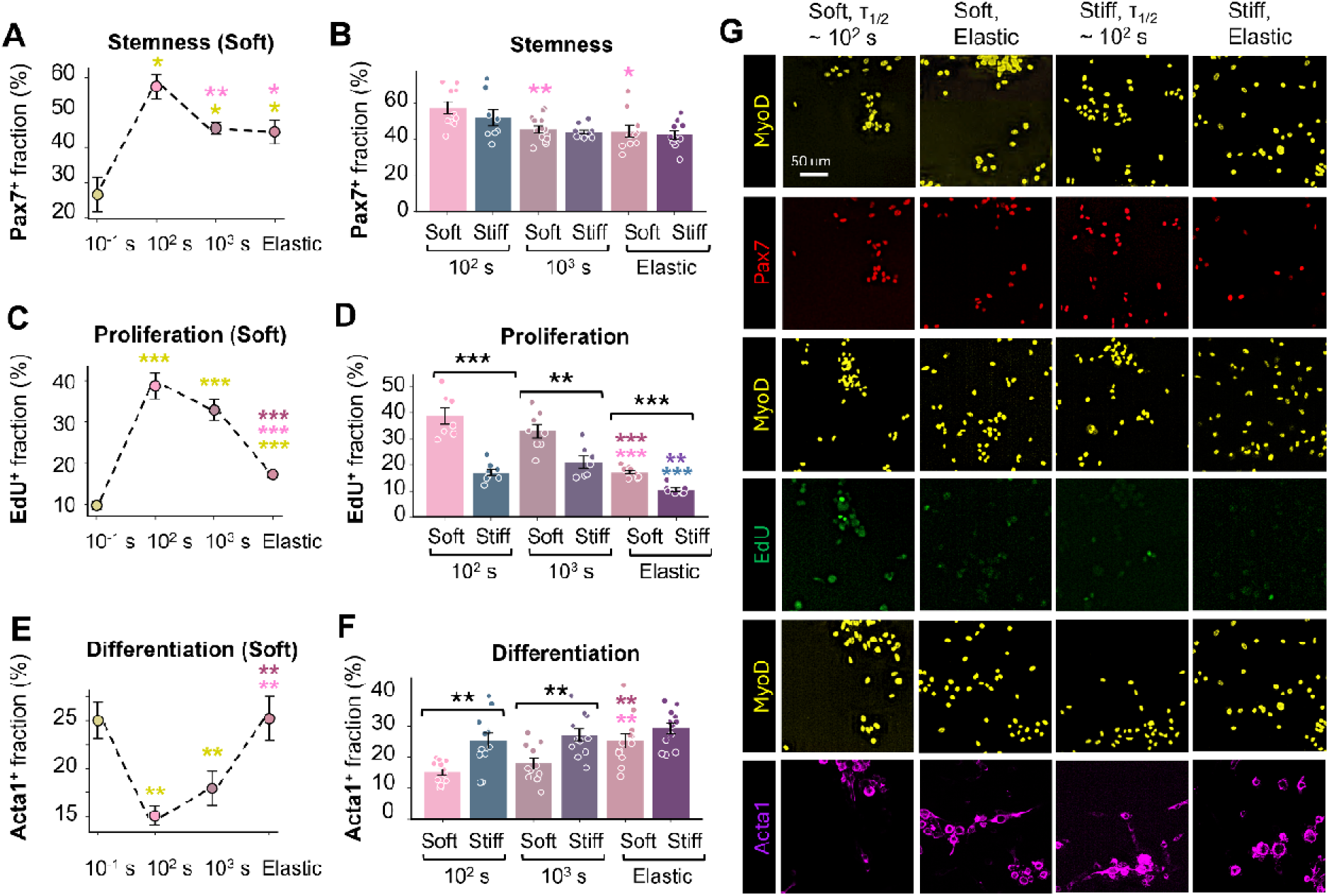
Matrix stiffness and stress relaxation synergistically regulate MuSC fate. (**A**-**F**) Quantification of Pax7^+^, EdU^+^, and Acta1^+^ MuSC populations cultured on 2D hydrogels with varying stress relaxation rates (τ_1/2_ ∼10^-1^, 10^2^, 10^3^, and ≫10^4^ s) under soft conditions (**A**, **C**, **E**) or varying stiffnesses (soft and stiff; τ_1/2_ ∼10^2^, 10^3^, and ≫10^4^ s) (**B**, **D**, **F**). (**G**) Representative immunofluorescence images of MuSCs stained for MyoD, Pax7, EdU, and Acta1 to assess stemness, proliferation, and differentiation after 7 days of culture. Each data point represents the fraction of positive cells (**A**-**F**) calculated from N ≥ 300 cells per hydrogel condition. Data are presented as mean ± s.d. from *n* ≥ 3 independent biological replicates. Statistical significance was determined using pairwise Welch’s *t*-tests with multiple-comparison correction. *P < 0.05 (*), P < 0.01 (**), and P < 0.001 (***)*. In (**A**-**F**), the color of the significance stars indicates th corresponding mechanical condition compared with the elastic control.

In contrast, hydrogels with the same stiffness but exhibiting either faster or slower stress relaxation, as well as predominantly elastic hydrogels, resulted in reduced proportions of proliferative stem-like cells (**Fig. 5C**) and increased differentiation (**Fig. 5E**). Furthermore, MuSCs cultured on fibrotic-like stiffness substrates (*G’* ∼ 13 kPa) displayed diminished proliferative capacity and increased differentiation, as evidenced by reduced EdU incorporation (**Fig. 5B**) and increased fractions of Acta1^+^ (**Fig. 5F**) and myogenin^+^ cells (**Fig. S15B,C**). These results are consistent with our findings that muscle-like viscoelasticity helps maintain high Rac/Cdc42:Rho activity ratio and enhanced Akt signaling, both of which contribute to the maintenance of MuSC stemness. Collectively, these findings demonstrate that stiffness and viscoelasticity jointly shape MuSC fate decisions.

### Focal adhesion-mediated Rac-PAK-Akt signaling establishes a viscoelastic memory in MuSCs

We previously demonstrated that the substrate stiffness MuSCs experienced during activation induced a mechanical memory that influenced subsequent cell fate decisions, even after th stiffness of the substrate was changed.^[10]^ Given that substrate viscoelasticity resulted in differences in cell fate after 7 days of culture, we finally asked whether the viscoelasticity of the substrate during activation similarly induces a mechanical memory, biasing later cell fate decisions. We thus investigated whether disrupting mechanotransduction during the first three days of culture on viscoelastic substrates could block viscoelastic mechanical memory formation and thereby impair myogenic progenitor expansion. Freshly isolated MuSCs were seeded onto soft, stress relaxing hydrogels and separately treated with inhibitors of FAK, Akt, DOCK1, TIAM1, and Cdc42 for the first 3 days of culture, after which the inhibitors were washed out, and cells were cultured for an additional 4 days (**Fig. 6A**). Transient inhibition of FAK or Akt on stress-relaxing hydrogels suppressed MuSC proliferation and increased differentiation, shifting these phenotypes toward levels observed in the soft, elastic vehicle control at day 7 (**Fig. 6B-D**). Similarly, inhibition of DOCK1, TIAM1, or Cdc42 reduced proliferative capacity and modestly decreased the Pax7□population (**Fig. S16A,B,E-G**).

**Fig. 6.**
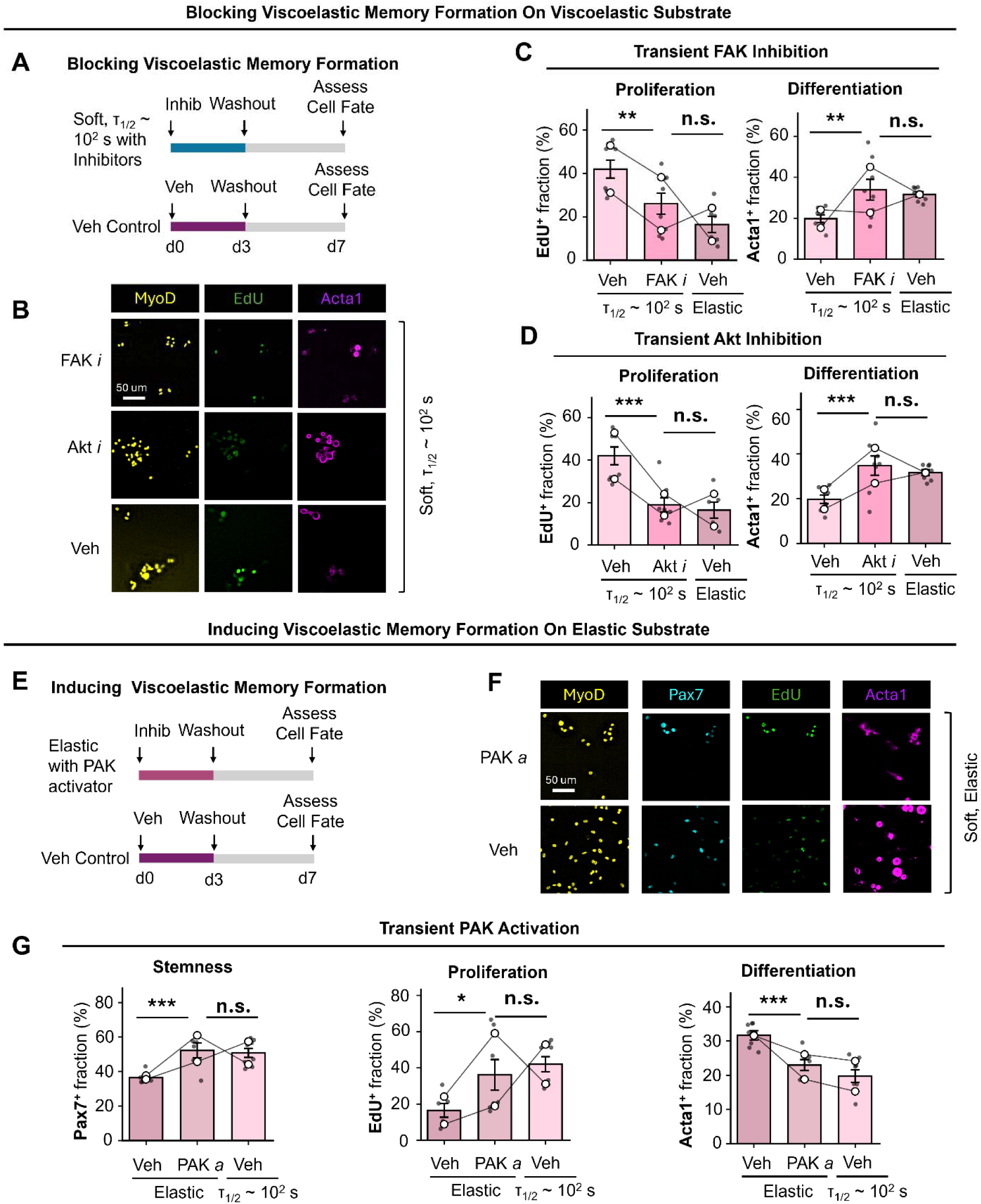
Focal adhesion-mediated FAK-Rac-PAK-Akt signaling establishes viscoelastic memory and regulates subsequent MuSC fate. (**A**) Experimental schematic for transient inhibition of signaling during the first 3 days of culture on soft hydrogels with physiological stress relaxation (τ_1/2_ ∼10^2^ s), followed by inhibitor washout and assessment of cell fate at day 7. (**B**) Representative immunofluorescence images of MuSCs stained for MyoD, EdU, and Acta1 following transient FAK or Akt inhibition. (**C, D**) Quantification of EdU^+^ and Acta1^+^ MuSC fractions following transient FAK (**C**) or Akt (**D**) inhibition. (**E**) Experimental schematic for transient PAK1 activation during the first 3 days of culture on predominantly elastic substrates, followed by washout and assessment of cell fate at day 7. (**F**) Representative immunofluorescence images of MuSCs stained for MyoD, Pax7, EdU, and Acta1 following transient PAK1 activation. (**G**) Quantification of Pax7^+^, EdU^+^, and Acta1^+^ MuSC fractions following transient PAK1 activation on soft elastic hydrogels. Each data point represents the fraction of positive cells calculated from ≥100 cells per hydrogel condition. Data are presented as mean ± s.d. from n ≥ 4 independent biological replicates. Open circles represent the mean values of independent experiments, with lines connecting paired experimental replicates. Statistical significance was determined using linear mixed-effects models (LMMs), with treatment condition included as a fixed effect and independent muscle batches included as a random effect. Pairwise comparisons were adjusted for multiple testing. *P < 0.05 (*), P < 0.01 (**), and P < 0.001 (***)*.

We next asked whether activating viscoelasticity-associated signaling during the MuSC activation window could induce a memory-like phenotype on predominantly elastic substrates. MuSCs cultured on elastic substrates were transiently treated with a PAK1 activator during the first 3 days of culture, followed by washout and continued culture until day 7 (**Fig. 6E**). Transient activation of PAK1 on predominantly elastic substrates increased the proliferative (EdU□) and stem-like (Pax7□) fractions while reducing commitment toward differentiation (Acta1□), restoring these phenotypes to levels observed under physiological viscoelasticity (**Figs. 6F,G, S16H,I**). Together, these findings suggest that FAK-Rac-PAK-Akt signaling is responsible for formation of a viscoelastic memory on soft, stress relaxing matrices, characterized by the persistent maintenance of a stem-like myogenic progenitor phenotype. Furthermore, enhancing PAK-Akt signaling during the first three days of culture is sufficient to recapitulate this viscoelastic memory on elastic substrates, thereby promoting MuSC self-renewal and limiting differentiation.

## Discussion

Chemically defined hydrogels were designed to recapitulate the physiological and pathophysiological viscoelastic mechanical properties of skeletal muscle. Healthy mouse tibialis anterior (TA) muscle exhibits a storage modulus of approximately 4 kPa and a stress relaxation half-time of ∼10^2^ s (**Fig. 1A**).^[7,17]^ These mechanical properties are altered during aging and disease. Aging is associated with thickening of the basal lamina and increased stiffness of the muscle sheath,^[61]^ whereas fibrosis is characterized by excessive collagen deposition and matrix remodeling,^[62]^ leading to increased tissue stiffness and altered viscoelastic properties.^[62,63]^ We therefore engineered a series of PEG-based hydrogels with independently tunable stiffness and stress relaxation dynamics recapitulating the range of mechanical properties encountered in healthy and pathological muscle in vitro (G’∼4-13 kPa and τ_1/2_∼10^2^-10^3^ s) to determine how physiological viscoelasticity impacts MuSC fate.

With this hydrogel system, we demonstrated that substrate viscoelasticity is a critical regulator of MuSC activation and subsequent fate determination (**Fig. 7**). Using hydrogels with independently tunable stiffness and stress relaxation behavior, we show that physiologically relevant matrix mechanics promote proliferation, preserve stemness, and suppress differentiation during extended culture. We further identify a biphasic relationship between MuSC activation and stress relaxation, whereby both excessively rapid and excessively slow relaxation impair early proliferation, highlighting the importance of matching the dynamic mechanical properties of native muscle.

**Fig. 7.**
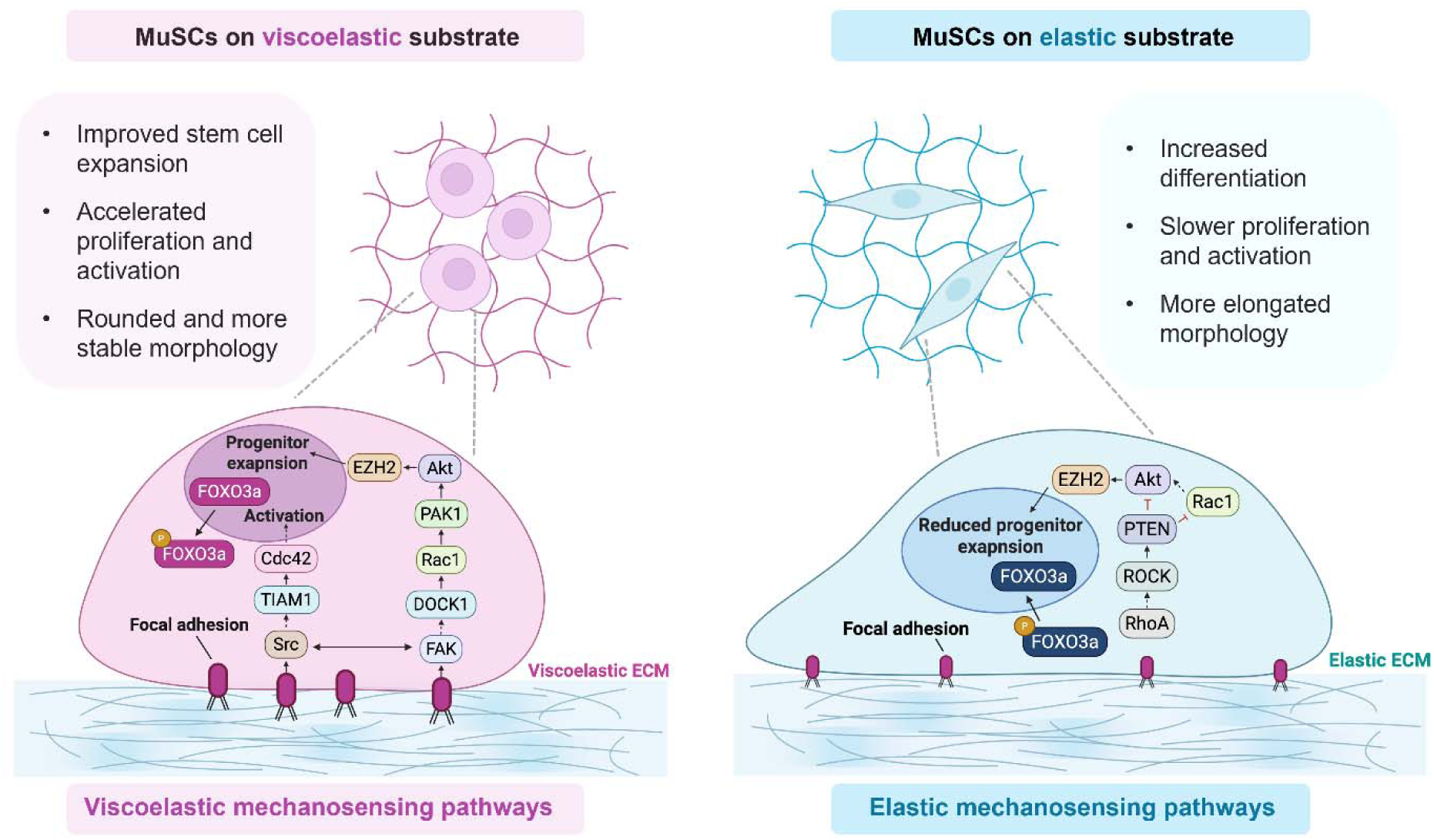
Proposed model of viscoelastic mechanosensing and mechanical regulation of MuSC fate. Schematic illustrating the distinct cellular responses and mechanotransduction pathway engaged by MuSCs on viscoelastic versus predominantly elastic substrates. MuSCs cultured on physiological viscoelastic substrates exhibit enhanced progenitor expansion, accelerated proliferation and activation, and a rounder, more stable cell morphology. Physiological viscoelasticity promotes focal adhesion signaling and activates Cdc42 and Rac1, enhancing Akt signaling, FOXO3a cytoplasmic localization, and EZH2-associated regulation of progenitor expansion and activation. In contrast, MuSCs cultured on predominantly elastic substrates exhibit slower proliferation and activation, reduced progenitor expansion, increased differentiation, and a more elongated morphology. Elastic substrates are associated with reduced focal adhesion stability and a shift toward RhoA-ROCK signaling, together with PTEN-associated suppression of Akt signaling and increased nuclear localization of FOXO3a.

Mechanistically, our results suggest that stiffness and viscous dissipation are sensed through distinct but complementary pathways. While the localization of known mechanosensitive transcriptional coactivators YAP/TAZ was altered by substrate stiffness, YAP/TAZ nuclear-to-cytoplasmic localization was not significantly affected by substrate stress relaxation dynamics. We therefore investigated the mechanisms underlying MuSC sensing of viscous dissipation. Physiologically relaxing matrices promoted focal adhesion stabilization, increased Rac/Cdc42:Rho activity ratios, enhanced Rac-PAK-Akt signaling, and increased FOXO3a nuclear-to-cytoplasmic translocation during early activation. Together, these findings identify focal adhesion-mediated mechanotransduction as a key link between substrate viscous dissipation and MuSC fate. The distinct pathways for stiffness and viscous sensing in MuSC contrast with previous studies of mesenchymal stromal cells and neural progenitor cells, in which integrin clustering was proposed as a common mechanism linking stiffness and viscous mechanosensing.^[64,65]^ Our findings instead suggest that stiffness and stress relaxation cooperate to regulate MuSC fate through separate mechanotransduction mechanisms that converge on downstream signaling programs governing activation, proliferation, and fate.

Our results extend the concept of stem cell mechanical memory, previously established for matrix stiffness,^[4,8,10,66,67]^ to viscous sensing. Transient inhibition of FAK-Rac-PAK-Akt signaling during the first three days of culture was sufficient to impair later MuSC self-renewal, whereas transient activation of PAK signaling on elastic substrates partially recapitulated the effects of physiological viscoelasticity. These findings suggest that the activation window represents a critical period during which MuSCs encode mechanical information that subsequently influences long-term fate decisions. Furthermore, the limited ability of small molecule perturbations to fully override mechanically-induced changes in cell fate indicates that viscoelasticity regulates a complex and interconnected signaling network, highlighting engineered microenvironments as a more effective strategy for maintaining stem cell function than targeting single biochemical signaling pathways.

Looking forward, these findings provide design principles for engineering biomaterials that preserve MuSC potency and suggest that restoring physiological viscoelasticity may represent a promising strategy to improve the success of cell-based regenerative therapies. More broadly, engineering mechanical memory through the physical microenvironment may provide a new strategy for programming stem cell state and long-term regenerative function, with matrix viscoelasticity serving as a critical design parameter.

## Experimental

### General Considerations

Reagents used for the synthesis of hydrogel precursors were purchased from Fisher Scientific, Sigma-Aldrich, Honeywell, TCI America, or BLDpharm unless otherwise noted. Water-sensitive reactions were conducted using commercially available anhydrous solvents. PEG-aldehyde (MW: 20 kDa, 4arm) was purchased from creative PEGWorks and PEG precursors were purchased from JenKem Technology. Aliphatic aldehyde-functionalized PEG (MW: 20 kDa, 8arm) was synthesized and characterized as previously described.^[22]^ Detailed synthetic procedures for DBCO-C6-NHS ester (1-{[6-(11,12-didehydrodibenzo[b,f]azocin-5(6H)-yl)-6-oxohexanoyl]oxy}-2,5-pyrrolidinedione) are provided in the Supplementary Materials. Hydrogel rheological characterization and muscle stiffness measurements were performed on a Rheometrics ARES-RFS3 strain-controlled rheometer. Muscle tissue relaxation was measured in compression using a TA Instruments Discovery HR-30 Hybrid rheometer. Fluorescence microscopy images were taken with a Zeiss LSM 900 laser scanning confocal microscope equipped with a broadband (385–730 nm) widefield epifluorescence light source and a Photometrics Prime BSI camera. ^1^H and ^13^C NMR spectra were recorded on a Bruker NEO 400 MHz or 600 MHz or Varian Inova 500 MHz spectrometers. Chemical shifts were reported in parts per million (ppm) and were referenced to the residual solvent peak (CDCl3, δ^1^H = 7.26 ppm, δ^13^C = 77.16 ppm, and DMSO-d6, δ^1^H = 2.50 ppm, δ^13^C = 39.52 ppm).

### Synthesis and characterization of CO-NHS ester

CO-O-COOH ((2-cyclooctyn-1-yloxy)acetic acid) was synthesized as described in the Supplementary Materials, following a known synthetic route.^[68]^ To prepare CO-NHS ester (cyclooctyne-O-NHS ester,) *N*-hydroxysuccinimide (NHS, 0.476 g, 4.13 mmol, 1.5 equiv.) and *N*-(3-dimethylaminopropyl)-*N* ′ -ethylcarbodiimide hydrochloride (EDC· HCl, 0.791 g, 4.13 mmol, 1.5 equiv.) were dissolved in anhydrous acetonitrile (10 mL) in a 50 mL round-bottom flask equipped with a magnetic stir bar. CO-O-COOH (0.500 g, 2.74 mmol, 1.0 equiv.) was then added, and the reaction vessel was purged with argon. The reaction was allowed to proceed under argon at room temperature for 24 h. Following completion, the solvent was removed under reduced pressure, and the residue was dissolved in dichloromethane (50 mL). The organic phase was washed sequentially with water (3 × 50 mL), and brine (50 mL), dried over anhydrous magnesium sulfate, and filtered. The filtrate was concentrated in vacuo to yield CO-NHS as a viscous yellow oil (0.704 g, yield: 91.8%). ^1^H NMR characterization is provided in **fig. S17**.

### Synthesis and characterization of (3-(Azidomethyl)phenyl)boronic acid

(3-(Azidomethyl)phenyl)boronic acid was synthesized according to a published protocol ^[33]^. Sodium azide (1.72 g, 5 equiv) and 3-bromomethylphenyl boronic acid (0.810 g, 1 equiv) were dissolved in anhydrous DMF (15 mL) and stirred at room temperature for 24 hours after which time the reaction was extracted with DCM (50 mL ×3). The combined organic layers were washed with brine (50 mL ×2), and dried over anhydrous magnesium sulfate (MgSO_4_), and concentrated in vacuo to give the product as a cream-colored solid (0.476 g, mass yield: 71.3%). Successful functionalization was confirmed by ^1^H NMR spectroscopy (**fig. S20**).

### Synthesis and Characterization of PEG-DBCO and PEG-CO

4-arm PEG-amine (MW = 10 kDa) and either DBCO-C6-NHS ester or CO-NHS ester (1.5 equiv. per amine) were fully dissolved in DMF in a two-neck round bottom flask. *N,N*-diisopropylethylamine (6 equiv. per amine) was added to the mixture, and the reaction was allowed to proceed at room temperature overnight. The resulting PEGs were precipitated into ice-cold diethyl ether, collected by centrifugation, and dried under argon. The pellet was re-dissolved in water, dialyzed against cold Milli-Q water for 2 days (MWCO 2kDa, 4 × 4.5 L, 4 °C), flash-frozen in liquid nitrogen, and lyophilized to yield a light-yellow solid. ^1^H NMR characterization is provided in **fig. S21 to 22**.

### Synthesis and characterization of PEG-nitrodopamine

Nitrodopamine was synthesized as described in the Supplementary Materials, following published protocols ^[32,34]^. PEG-*p*-nitrophenylcarbonate (PEG-NPC) was prepared according to a published protocol with minor modifications ^[34]^. Briefly, 4-arm PEG-hydroxyl (MW = 10 kDa) was dissolved in anhydrous DCM in a round-bottom flask. Then, *p*-nitrophenyl chloroformate (8 equiv. per hydroxyl) and TEA (9 equiv. per hydroxyl) were added, and the reaction was allowed to proceed for 48 h at room temperature under argon. The resulting PEG-NPC was precipitated into ice-cold diethyl ether, collected by centrifugation, and dried under argon.

To synthesize PEG-nitrodopamine, PEG-NPC was dissolved in anhydrous DMF and reacted with nitrodopamine (3 equiv. per NPC group) and TEA (6 equiv. per NPC group) in a round-bottom flask under argon. The reaction was allowed to proceed for 48 h at room temperature. The resulting PEG-ND was precipitated into ice-cold diethyl ether, collected by centrifugation, and dried under argon. The product was then dissolved in Milli-Q water, dialyzed against cold Milli-Q water (MWCO 2 kDa, 4 × 4.5 L, 4 °C) for 2 days, flash-frozen in liquid nitrogen, and lyophilized to yield a yellow solid. Successful functionalization was confirmed by ^1^H NMR spectroscopy (**fig. S23**).

### Synthesis and characterization of PEG-hydrazine

PEGs were functionalized with hydrazine moieties through amide coupling of the terminal amines, following established methods ^[22,69]^. Briefly, multi-arm PEG-amine HCl salts (MW: 10 kDa, 4arm; MW: 20 kDa, 8arm) were fully dissolved in anhydrous DMF at room temperature. In a separate round-bottom flask, tri-Boc-hydrazinoacetic acid (4 equiv. per amine) and HATU (4 equiv. per amine; Advanced Chemtech) were dissolved in DMF. Diisopropylethylamine (DIEA, 10 equiv. per amine) was added, and the solution was stirred for 10 min to activate the carboxylic acid. The activated mixture was then added dropwise to the PEG solution, and the reaction was allowed to proceed overnight at room temperature. The resulting PEG-tri-Boc-hydrazine was precipitated by dropwise addition into ice-cold diethyl ether and collected by centrifugation. A portion (10 mg) was reserved for ^1^H NMR analysis.

The remaining product was dissolved in 6 mL of a 1:1 (v:v) mixture of DCM:trifluoroacetic acid (TFA) plus triisopropylsilane (TIPS, 100 µL) and MilliQ-grade water (100 µL) to remove the Boc protecting group. The deprotection reaction was carried out at room temperature for 4 h. The resulting PEG-hydrazine was precipitated into ice-cold diethyl ether, collected by centrifugation, and dried under argon. A portion (10 mg) was reserved to confirm deprotection by ^1^H NMR spectroscopy. The remaining product was dissolved in Milli-Q water, dialyzed against cold Milli-Q water for 2 days (MWCO 2kDa, 4 × 4.5 L, 4 °C), flash-frozen in liquid nitrogen, and lyophilized to yield a white solid. ^1^H NMR characterization is shown in **fig. S24 to 25**.

### Synthesis and characterization of azide-functionalized PEG

To prepare 20 kDa 8-arm PEG-azide (PEG-N_3_), 20 kDa 8-arm PEG-OH (10 g, 4 mmol -OH, 1 eq.) was dissolved in anhydrous DCM (40 mL), followed by the addition of triethylamine (0.84 mL, 6.0 mmol, 1.5 eq.). The mixture was cooled on an ice bath and methanesulfonylchloride (0.46 mL, 6 mmol, 1.5 eq.) was added dropwise over 30 min. The mixture was stirred overnight at room temperature, concentrated to a volume of a few milliliters, precipitated in cold diethyl ether and then centrifuged to collect crude intermediate. The crude intermediate was dissolved in DMF (20 mL) without further purification and sodium azide (0.40 g, 6.0 mmol) was added slowly to the solution. The reaction mixture was stirred at 80 °C overnight. After cooling to room temperature, the mixture was precipitated in cold diethyl ether and centrifuged to collect the crude product. The crude product was redissolved in DCM (20 mL) and filtered. The organic phase was washed with H_2_O (40 mL) and brine (20 mL), dried over anhydrous MgSO_4_, and concentrated to a few milliliters. The product was precipitated in cold diethyl ether, isolated via centrifugation, and dried under vacuum to obtain product as a white powder (86% yield). ^1^H NMR and FT-IR characterization is provided in **fig. S26 to 27**.

### Hydrogel formation

To prepare the dual network hydrogels, PEG precursors were dissolved in PBS at room temperature and maintained on ice until used. The precursor concentrations were chosen to maintain a ratio of hydrazine to aldehyde reactive groups > 1, and 1.5:1 ratio of azide to alkyne (DBCO or CO). Hydrogel compositions are detailed in **tables S1 and S2**. To prepare the fast-relaxing hydrogels, (3-(azidomethyl)phenyl)boronic acid was dissolved in ethanol to a concentration of 20 mg/ml. PEG precursors were dissolved in PBS at room temperature and maintained on ice until used. Hydrogel compositions are detailed in **table S3**. Elastic hydrogels were prepared with a fixed PEG-N concentration and varying PEG-CO concentrations to tune hydrogel stiffness. Detailed hydrogel compositions are provided in **table S4**.

### Hydrogel rheological characterization

Oscillatory rheology was performed using hydrogel precursor solutions prepared as described above and a 25 mm, 0.02-rad cone-and-plate geometry. For hydrogels targeting a storage modulus of *G*′ ∼ 4 kPa, precursor solutions were gelled in situ on a Peltier stage maintained at 22 °C. Time sweep measurements were performed for 1 h at 22 °C using 5% strain and a frequency of 1 Hz. Following gelation, temperature ramps from 22 °C to 37 °C were conducted at 5% strain.

For hydrogels targeting *G*΄ ∼ 13 kPa and the fast-relaxing hydrogel formulation, precursor solutions were gelled in situ at 37 °C. To prevent premature gelation during sample loading, the Peltier stage was first cooled to 4 ° C and subsequently heated to 37 °C. Time sweep measurements were performed for 5 h at 37 °C using 5% strain and a frequency of 1 Hz. No temperature ramp was applied to these formulations.

For all hydrogel formulations, frequency sweeps were performed at 37 °C over a frequency range of 0.1–10 Hz at a constant strain of 3% (**fig. S1G**). Strain sweeps were conducted at 37 °C and 1 Hz, with strain varying from 0.1% to 10% (**fig. S1H**). Stress relaxation measurements were performed at 37 °C by applying a constant strain of 5%. Stress relaxation half-times (τ_1/2_) were calculated from the measured stress relaxation curves (**Fig. 1, D and F, fig. S1C**). Storage moduli (G′) were determined from the frequency sweeps and are reported as the average value measured at approximately 1 Hz (**Fig. 1, F and G**).

### Muscle tissue rheological characterization

Tissue stiffness was measured by oscillatory rheology with freshly collected TA muscles from young (∼2 mo) mice using an 8 mm parallel plate geometry. Time sweep measurements were performed for 5 min at room temperature using 1% strain and a frequency of 1 Hz, and frequency sweeps were performed at room temperature over a frequency range of 0.1–10 Hz at a constant strain of 1%. Storage moduli (G′) were determined from the frequency sweeps and are reported as the average value measured at approximately 1 Hz (**Fig. 1A**).

Tissue stress relaxation was measured by compression testing with freshly collected TA muscles of young (∼2 mo) mice using a 25 mm parallel plate geometry at room temperature. Following determination of the sample thickness, tissues were compressed at a constant strain rate of 1% strain s ¹ to a final compressive strain of 10%. The applied strain was then held constant while the decay in normal stress was recorded over time. Stress relaxation half-times (τ_1/2_) were calculated from the resulting relaxation curves. Representative stress relaxation profiles are shown in **Fig. 1A**.

### Hydrogel stability

Hydrogel stability was assessed by measuring the mass swelling ratio over time. Briefly, 50 μL hydrogel samples were cast in pre-weighed vials and allowed to gel overnight at 37 °C in a humidified incubator. Gels were then incubated in 1 mL of MuSC culture medium (FluoroBrite DMEM supplemented with 15% fetal bovine serum (FBS), 1× L-glutamine, 1× sodium pyruvate, 1× non-essential amino acids, 1× penicillin/streptomycin, and 2.5 ng mL ¹ FGF-2) for 0, 1, 3, or 7 days. At each time point, the medium was removed and the hydrated gels were weighed to determine their swollen mass. Samples were subsequently frozen at −80 °C, lyophilized overnight, and weighed again to determine their dry mass. The mass swelling ratio was calculated as the ratio of the mass of water in the swollen hydrogel to the dry mass of the polymer network.

### Laminin functionalization and quantification

Laminin functionalization was performed according to previous protocols with some modifications.^[7,70]^ After buffer exchanging into PBS by dialysis, laminin-111 was reacted with a 50-fold molar excess of N□-PEG-NHS (MW ∼ 3.5 kDa) to generate azide-functionalized laminin-111 or with a 50-fold molar excess of Ald-Ph-PEG8-NHS ester (BroadPharm, BP-40630) to generate benzaldehyde-functionalized laminin-111. Azide-functionalized laminin-111 was used for ligand incorporation into purely elastic and fast-relaxing hydrogels, whereas both azide- and benzaldehyde-functionalized laminin-111 were incorporated into dual-network hydrogels. Laminin density was quantified using Cy5-labeled laminin and monitored daily for up to 8 days, following a previously established protocol ^[10]^.

### Muscle stem cell isolation

All animal experiments were approved by the Institutional Animal Care and Use Committee of the University of Pennsylvania and performed in compliance with institutional guidelines. Young (∼2 mo) C57BL/6 mice (The Jackson Laboratory) were used for immunofluorescence and cell division analyses. Pax7-EGFP reporter mice,^[39]^ generously provided by Prof. Foteini Mourkioti (University of Pennsylvania), were used for live-cell imaging to quantify changes in progenitor cell numbers. MuSCs were isolated from the hindlimb muscles of mice as previously described with minor modifications.^[7,10,70]^ Briefly, dissected muscles were enzymatically dissociated with collagenase and dispase using a gentleMACS Octo Dissociator (Miltenyi Biotec) and filtered through a 60 μm cell strainer. Cell suspensions were incubated with PE-Cy7-conjugated antibodies against CD45, CD11b, CD31, and Sca1 (BD Biosciences) to exclude hematopoietic, endothelial, and stromal cell populations, and stained with integrin-α7-PE (Ablab) and CD34-Alexa Fluor 647 (BD Biosciences) to identify MuSCs. MuSCs were isolated by fluorescence-activated cell sorting (FACS) as integrin-α7□CD34□cells. Antibodies used for FACS sorting are summarized in **table S5**.

### Hydrogel preparation for cell culture

PEG hydrogels for MuSC culture were formed directly in glass bottom 96-well plates (Cellvis). Glass surfaces were chemically-functionalized via silane chemistry to enable covalent hydrogel attachment. For dual-network hydrogels, 3-aminopropyltriethoxysilane (1 equiv.) and *N*-succinimidyl 4-formylbenzoate ^[71,72]^ (1.25 equiv.) were dissolved in anhydrous DMF, followed by the addition of *N,N*-diisopropylethylamine (6 equiv.). The reaction was allowed to proceed overnight at room temperature with agitation. The resulting solution was diluted with ethanol, acetic acid, and (3-azidopropyl)trimethoxysilane to final concentrations of 90% ethanol, 10% acetic acid, and 1% silane. For purely elastic and fast-relaxing hydrogels, glass surfaces were functionalized with (3-azidopropyl)trimethoxysilane as previously described.^[10,70]^

Sterile PEG precursor solutions were prepared on ice and mixed according to the formulations described above. Hydrogel precursor solutions (15 μL per well) were immediately transferred to silanized wells and centrifuged at 2,000 × g for 15 min to ensure uniform coating of the glass surface. Samples were then centrifuged at 2,000 × g at 40 °C for 1 h to facilitate strain-promoted azide-alkyne cycloaddition (SPAAC) crosslinking, followed by incubation at 37 °C for 5 h in a humidified incubator to complete gelation.

Hydrogels were subsequently functionalized with laminin-111 overnight at room temperature with gentle agitation. Purely elastic and fast-relaxing hydrogels were incubated with azide-functionalized laminin-111, whereas dual-network hydrogels were incubated with a mixture of azide-functionalized and benzaldehyde-functionalized laminin-111. Prior to cell seeding, laminin solutions were removed and hydrogels were washed three times with sterile PBS. The plates were then equilibrated in cell culture medium at 37°C for 30 min before cell seeding.

### Muscle stem cell culture and analysis

Cells were cultured as previously described with minor modifications.^[10]^ Freshly isolated MuSCs were seeded directly onto hydrogels at a density of 3,000 cells per well for the day 1 and day 3 time points, and 500 cells per well for the day 7 time point. Cells were cultured in MuSC culture medium, with medium changes performed on days 3 and 6. For mechanotransduction perturbation studies, cells were treated with the indicated inhibitors or activators at the time of seeding. Reagents and their concentrations used in each experiment are summarized in **table S6**.

### Immunofluorescence analysis of cultured MuSCs

Cells were fixed, permeabilized, blocked, stained, and washed according to established protocols.^[10,70]^ Antibodies used for general immunofluorescence staining are listed in **tables S7 to 8**. EdU incorporation assays and RhoA and Rac1 GTPase activity staining were performed as previously described.^[10]^ Samples were imaged with Zeiss LSM 900 laser scanning confocal microscope equipped with a broadband (385–730 nm) widefield epifluorescence light source. For whole-gel analysis, 6 × 6 tiled images were acquired and processed using our previously reported GPU-accelerated image processing pipeline (CRISP) for image deconvolution and stitching.^[10,70,73]^ Quantitative image analysis was performed using CellProfiler.^[74]^

### Live cell imaging and analysis

Live imaging of muscle stem cells was performed using the Zeiss LSM900 microscope under incubation conditions of 37°C and 5% CO□. For live cell imaging shown in Fig. 4A to B, cells were seeded on hydrogels immediately after isolation, and imaging was started two days after seeding. Phase contrast images were acquired every 30 minutes for a total duration of one day using a 20× objective lens. For live-cell imaging shown in fig. S2B to E, freshly isolated MuSCs were seeded on hydrogels immediately after isolation, and imaging was initiated 20 h after seeding. Phase-contrast images were acquired at 33-min intervals for 2 days using a 20× objective. To analyze the cell images, we used a manually trained segmentation model based on Cellpose ^[75]^ to generate binary cell masks at each time point. Cell tracking was then performed using the Hungarian algorithm.^[76]^

### Focal adhesion staining and quantification

For paxillin staining, cells were cultured on hydrogel substrates for 36 h, the culture medium was aspirated, and the samples were fixed with 4% paraformaldehyde for 40 min at 37°C. Samples were then permeabilized with 0.1% Triton X-100 for 3 min and blocked with blocking solution (5% w/v BSA and 5% v/v goat serum in DPBS) for 3 h at room temperature. Paxillin primary antibody (BD Biosciences, 610051; 1:100) was incubated overnight at 4°C in antibody dilution solution (2.5% w/v BSA and 2.5% v/v goat/donkey serum in DPBS), followed by overnight incubation with secondary antibody (1:400) in the same solution. Three DPBS washes were performed after each antibody incubation. Confocal imaging was performed using a Zeiss LSM900 microscope with a 63× objective. Focal adhesion size and intensity were quantified with an online Focal Adhesion Analysis Server.^[55]^

### Statistical analysis

Statistical analyses were performed in Python (version 3.12.4) using Jupyter Notebook, with data processing and statistical testing conducted using pandas, SciPy, statsmodels, and NumPy. Sample sizes (n) are indicated in the corresponding figure legends and represent independent biological replicates unless otherwise stated. Pairwise comparisons were performed using two-sided Welch’s t-tests. For datasets pooled across independent MuSC isolations, linear mixed-effects models (LMMs) were used with biological replicate (isolation) included as a random effect and experimental condition included as a fixed effect. P values were adjusted for multiple comparisons where appropriate. n.s. = not significant (p > 0.05), *p < 0.05, **p < 0.01, ***p < 0.001.

## Acknowledgements

The authors thank Prof. Foteini Mourkioti for providing the Pax7-EGFP reporter mice, Chung Yin Matthew Cheung for assistance with MuSC isolation, and Prof. Claudia Loebel for access to mechanical testing equipment. The authors gratefully acknowledge the use of facilities and instrumentation supported by U.S. National Science Foundation (NSF) through the University of Pennsylvania Materials Research Science and Engineering Center (MRSEC) (DMR-2309043). T.Y.K. was supported by a Postdoctoral Fellowship Program (Nurturing Next-generation Researchers) granted by the National Research Foundation of Korea (RS-2024-00407821). Funding for this work was provided by the U.S. National Institutes of Health (NIH, R00 AG071738 and R35 GM154913 to C.M.M.) and a seed grant from the Center for Precision Engineering for Health (CPE4H, to C.M.M.).

## Author Contributions

H.H. and C.M.M designed research; H.H., T.Y.K., and Z.G. performed research; H.H., T.Y.K. and P.Z. contributed new reagents/analytic tools; H.H. and T.Y.K. analyzed data; and H.H. and C.M.M. wrote the paper.

## Data, code, and materials availability

All source data will be made publicly available upon final publication.

## References

[1] H. M. Blau, G. Q. Daley, N. Engl. J. Med. 2019, 380, 1748.

[2] E. Fuchs, H. M. Blau, Cell Stem Cell 2020, 27, 532.

[3] C. M. Madl, S. C. Heilshorn, Annu Rev Biomed Eng 2018, 20, 21.

[4] A. J. Engler, S. Sen, H. L. Sweeney, D. E. Discher, Cell 2006, 126, 677.

[5] N. Huebsch, P. R. Arany, A. S. Mao, D. Shvartsman, O. A. Ali, S. A. Bencherif, J. Rivera-Feliciano, D. J. Mooney, Nat Mater 2010, 9, 518.

[6] K. Saha, A. J. Keung, E. F. Irwin, Y. Li, L. Little, D. V. Schaffer, K. E. Healy, Biophys J 2008, 95, 4426.

[7] P. M. Gilbert, K. L. Havenstrite, K. E. G. Magnusson, A. Sacco, N. A. Leonardi, P. Kraft, N. K. Nguyen, S. Thrun, M. P. Lutolf, H. M. Blau, Science 2010, 329, 1078.

[8] C. Yang, M. W. Tibbitt, L. Basta, K. S. Anseth, Nat. Mater. 2014, 13, 645.

[9] C. X. Li, N. P. Talele, S. Boo, A. Koehler, E. Knee-Walden, J. L. Balestrini, P. Speight, A. Kapus, B. Hinz, Nat Mater 2017, 16, 379.

[10] C. M. Madl, Y. X. Wang, C. A. Holbrook, S. Su, X. Shi, F. J. Byfield, G. Wicki, I. A. Flaig, H. M. Blau, Proc. Natl. Acad. Sci. 2024, 121, e2406787121.

[11] C. M. Madl, S. C. Heilshorn, H. M. Blau, Nature 2018, 557, 335.

[12] E. Gussoni, G. K. Pavlath, A. M. Lanctot, K. R. Sharma, R. G. Miller, L. Steinman, H. M. Blau, Nature 1992, 356, 435.

[13] J. P. Tremblay, F. Malouin, R. Roy, J. Huard, J. P. Bouchard, A. Satoh, C. L. Richards, Cell Transplant. 1993, 2, 99.

[14] Mendell Jerry R., Kissel John T., Amato Anthony A., King Wendy, Signore Linda, Prior Thomas W., Sahenk Zarife, Benson Sandra, McAndrew Patricia E., Rice Robert, Nagaraja Haikady, Stephens Ralph, Lantry Laura, Morris Glen E., Burghes Arthur H.M., N. Engl. J. Med. 1995, 333, 832.

[15] A. Sacco, R. Doyonnas, P. Kraft, S. Vitorovic, H. M. Blau, Nature 2008, 456, 502.

[16] O. Chaudhuri, L. Gu, D. Klumpers, M. Darnell, S. A. Bencherif, J. C. Weaver, N. Huebsch, H. Lee, E. Lippens, G. N. Duda, D. J. Mooney, Nat. Mater. 2016, 15, 326.

[17] O. Chaudhuri, J. Cooper-White, P. A. Janmey, D. J. Mooney, V. B. Shenoy, Nature 2020, 584, 535.

[18] G. A. Hoffmann, J. Y. Wong, M. L. Smith, Biochemistry 2019, 58, 4710.

[19] A. R. Cameron, J. E. Frith, J. J. Cooper-White, Biomaterials 2011, 32, 5979.

[20] E. Qiao, C. A. Fulmore, D. V. Schaffer, S. Kumar, Proc. Natl. Acad. Sci. 2024, 121, e2317711121.

[21] M. S. Huang, B. L. LeSavage, S. Ghorbani, A. E. Gilchrist, J. G. Roth, C. Huerta-López, E. A. Mozipo, R. S. Navarro, S. C. Heilshorn, Nat. Commun. 2025, 16, 5213.

[22] L. E. Brown, D. Bakker, P. Zhou, C. M. Madl, J. Mater. Chem. B 2026, 14, 4740.

[23] T.-L. Chang, A. N. Borelli, A. A. Cutler, B. B. Olwin, K. S. Anseth, Acta Biomater. 2025, 192, 48.

[24] E. B. Roloson, W.-H. Jung, S. L. McNamara, C. L. Van Stone, N. Lafuente-Gómez, D. M. Morgan, R. M. Gibbs, L. L. Rubin, G. N. Duda, D. J. Mooney, Adv. Healthc. Mater. 2026, 15, e02775.

[25] J. J. Dowling, Nat. Rev. Neurol. 2016, 12, 675.

[26] R. J. Butterfield, P. B. Shieh, H. Li, M. Binks, T. G. McDonnell, K. A. Ryan, M. Delnomdedieu, B. A. Belluscio, S. Neelakantan, D. I. Levy, P. F. Schwartz, E. C. Smith, Nat. Med. 2025, 1.

[27] C. Pascual-Morena, S. Patiño-Cardona, I. Martínez-García, J. Fernández-Bravo-Rodrigo, M. Garrido-Miguel, Hum. Genet. 2025, DOI 10.1007/s00439-025-02758-6.

[28] N. Elangkovan, G. Dickson, J. Neuromuscul. Dis. 2021, 8, S303.

[29] D. D. McKinnon, D. W. Domaille, J. N. Cha, K. S. Anseth, Adv. Mater. 2014, 26, 865.

[30] B. L. LeSavage, D. Zhang, C. Huerta-López, A. E. Gilchrist, B. A. Krajina, K. Karlsson, A. R. Smith, K. Karagyozova, K. C. Klett, M. S. Huang, C. Long, G. Kaber, C. M. Madl, P. L. Bollyky, C. Curtis, C. J. Kuo, S. C. Heilshorn, Nat. Mater. 2024, 23, 1138.

[31] M. W. Jaeschke, A. N. Borelli, N. P. Skillin, T. J. White, K. S. Anseth, Adv. Healthc. Mater. 2024, 13, 2400062.

[32] B. Marco-Dufort, R. Iten, M. W. Tibbitt, J. Am. Chem. Soc. 2020, 142, 15371.

[33] A. Yu. Fedorov, A. A. Shchepalov, A. V. Bol"shakov, A. S. Shavyrin, Yu. A. Kurskii, J.-P. Finet, S. V. Zelentsov, Russ. Chem. Bull. 2004, 53, 370.

[34] S. Tang, H. Ma, H.-C. Tu, H.-R. Wang, P.-C. Lin, K. S. Anseth, Adv. Sci. 2018, 5, 1800638.

[35] S. E. Brashear, R. P. Wohlgemuth, G. Gonzalez, L. R. Smith, J. Physiol. 2021, 599, 943.

[36] J. S. Silver, K. A. Günay, A. A. Cutler, T. O. Vogler, T. E. Brown, B. T. Pawlikowski, O. J. Bednarski, K. L. Bannister, C. J. Rogowski, A. G. Mckay, F. W. DelRio, B. B. Olwin, K. S. Anseth, Sci. Adv. 2021, 7, eabe4501.

[37] M. T. Webster, U. Manor, J. Lippincott-Schwartz, C.-M. Fan, Cell Stem Cell 2016, 18, 243.

[38] Y. He, Y. Heng, Z. Qin, X. Wei, Z. Wu, J. Qu, Sci. Adv. 2023, 9, eadi1891.

[39] E. D. Tichy, D. K. Sidibe, C. D. Greer, N. M. Oyster, P. Rompolas, N. A. Rosenthal, H. M. Blau, F. Mourkioti, Skelet. Muscle 2018, 8, 27.

[40] P. Rocheteau, B. Gayraud-Morel, I. Siegl-Cachedenier, M. A. Blasco, S. Tajbakhsh, Cell 2012, 148, 112.

[41] Y. Ono, Y. Urata, S. Goto, S. Nakagawa, P. O. Humbert, T.-S. Li, P. S. Zammit, Cell Rep. 2015, 10, 1135.

[42] J. Segalés, E. Perdiguero, P. Muñoz-Cánoves, Front. Cell Dev. Biol. 2016, 4, DOI 10.3389/fcell.2016.00091.

[43] A. H. Juan, A. Derfoul, X. Feng, J. G. Ryall, S. Dell’Orso, A. Pasut, H. Zare, J. M. Simone, M. A. Rudnicki, V. Sartorelli, Genes Dev. 2011, 25, 789.

[44] L. García-Prat, E. Perdiguero, S. Alonso-Martín, S. Dell’Orso, S. Ravichandran, S. R. Brooks, A. H. Juan, S. Campanario, K. Jiang, X. Hong, L. Ortet, V. Ruiz-Bonilla, M. Flández, V. Moiseeva, E. Rebollo, M. Jardí, H.-W. Sun, A. Musarò, M. Sandri, A. del Sol, V. Sartorelli, P. Muñoz-Cánoves, Nat. Cell Biol. 2020, 22, 1307.

[45] F. Yue, P. Bi, C. Wang, T. Shan, Y. Nie, T. L. Ratliff, T. P. Gavin, S. Kuang, Nat. Commun. 2017, 8, 14328.

[46] S. D. Gopinath, A. E. Webb, A. Brunet, T. A. Rando, Stem Cell Rep. 2014, 2, 414.

[47] T. P. Lele, A. Brock, S. R. Peyton, Ann. Biomed. Eng. 2020, 48, 2103.

[48] A. P. Kann, M. Hung, W. Wang, J. Nguyen, P. M. Gilbert, Z. Wu, R. S. Krauss, Cell Stem Cell 2022, 29, 933.

[49] S. Eliazer, J. M. Muncie, J. Christensen, X. Sun, R. S. D’Urso, V. M. Weaver, A. S. Brack, Cell Stem Cell 2019, 25, 654.

[50] Y. Peng, J. Du, R. Li, S. Günther, N. Wettschureck, S. Offermanns, Y. Wang, A. Schneider, T. Braun, Cell Discov. 2024, 10, 76.

[51] M. D. Boppart, D. J. Burkin, S. J. Kaufman, Biochim. Biophys. Acta 2011, 1812, 439.

[52] N. Moriya, M. Miyazaki, Am. J. Physiol.-Regul. Integr. Comp. Physiol. 2018, 314, R741.

[53] G. Wang, H. Zhu, C. Situ, L. Han, Y. Yu, T. H. Cheung, K. Liu, Z. Wu, EMBO J. 2018, 37, EMBJ201798239.

[54] J. T. Rodgers, K. Y. King, J. O. Brett, M. J. Cromie, G. W. Charville, K. K. Maguire, C. Brunson, N. Mastey, L. Liu, C.-R. Tsai, M. A. Goodell, T. A. Rando, Nature 2014, 510, 393.

[55] M. E. Berginski, S. M. Gomez, F1000Research 2013, 2, 68.

[56] M. Higuchi, K. Onishi, C. Kikuchi, Y. Gotoh, Nat. Cell Biol. 2008, 10, 1356.

[57] E. J. Koubek, L. C. Santy, Cells 2022, 11, 3565.

[58] M. E. Gonzalez, G. D. Naimo, T. Anwar, A. Paolì, S. R. Tekula, S. Kim, N. Medhora, S. A. Leflein, J. Itkin, R. Trievel, K. M. Kidwell, Y.-C. Chen, L. Mauro, E. Yoon, S. Andò, C. G. Kleer, iScience 2022, 25, DOI 10.1016/j.isci.2022.104827.

[59] S. K. Quadri, Microvasc. Res. 2012, 83, 3.

[60] S. Yang, H.-M. Kim, Biomaterials 2012, 33, 2902.

[61] H. E. Boers, M. Haroon, F. Le Grand, A. D. Bakker, J. Klein Nulend, R. T. Jaspers, J. Orthop. Res. 2018, 36, 632.

[62] L. R. Smith, E. R. Barton, Am. J. Physiol. - Cell Physiol. 2014, 306, C889.

[63] W. Zhang, M. Jadidi, S. A. Razian, G. A. Holzapfel, A. Kamenskiy, D. A. Nordsletten, Acta Biomater. 2024, 188, 223.

[64] M. Darnell, A. O’Neil, A. Mao, L. Gu, L. L. Rubin, D. J. Mooney, Proc. Natl. Acad. Sci. 2018, 115, E8368.

[65] M. A. Schwartz, D. W. DeSimone, Curr. Opin. Cell Biol. 2008, 20, 551.

[66] N. Bouhrira, S. Pizarro, V. Senf, K. B. Margulies, 2025.

[67] J. L. Balestrini, S. Chaudhry, V. Sarrazy, A. Koehler, B. Hinz, Integr. Biol. 2012, 4, 410.

[68] M. Burk, S. Rothstein, P. Dubé, Org. Process Res. Dev. 2018, 22, 108.

[69] B. M. Richardson, C. J. Walker, L. J. Macdougall, J. W. Hoye, M. A. Randolph, S. J. Bryant, K. S. Anseth, Biomater. Sci. 2020, 8, 3804.

[70] C. M. Madl, I. A. Flaig, C. A. Holbrook, Y. X. Wang, H. M. Blau, Biomaterials 2021, 275, 120973.

[71] C. A. DeForest, D. A. Tirrell, Nat. Mater. 2015, 14, 523.

[72] P. Zhou, L. Brown, C. M. Madl, Adv. Funct. Mater. 2025, 35, 2422047.

[73] A. R. Palla, M. Ravichandran, Y. X. Wang, L. Alexandrova, A. V. Yang, P. Kraft, C. A. Holbrook, C. M. Schürch, A. T. V. Ho, H. M. Blau, Science 2021, 371, eabc8059.

[74] C. McQuin, A. Goodman, V. Chernyshev, L. Kamentsky, B. A. Cimini, K. W. Karhohs, M. Doan, L. Ding, S. M. Rafelski, D. Thirstrup, W. Wiegraebe, S. Singh, T. Becker, J. C. Caicedo, A. E. Carpenter, PLOS Biol. 2018, 16, e2005970.

[75] C. Stringer, T. Wang, M. Michaelos, M. Pachitariu, Nat. Methods 2021, 18, 100.

[76] H. W. Kuhn, Nav. Res. Logist. Q. 1955, 2, 83.

