## Supporting Information for "Tissue-like Viscoelasticity Drives Muscle Stem Cell Expansion Through Mechanical Memory Formation"

**Supporting Text**

1. DBCO synthesis

DBCO-$C_6$-NHS ester was synthesized according to a previously published protocols with some modifications (*1*, *2*).

1.1 Synthesis of Dibenzosuberenone Oxime

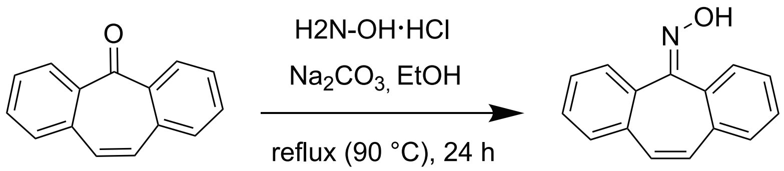

Dibenzosuberenone (15.0 g, 72 mmol, 1.0 equiv.) was dissolved in anhydrous ethanol (90 mL) in a 500 mL round-bottom flask. Hydroxylamine hydrochloride (20.23 g, 291 mmol, 4.0 equiv.) and sodium carbonate (11.57 g, 109 mmol, 1.5 equiv.) were added, and the flask was fitted with a reflux condenser and purged with argon. The reaction mixture was heated to reflux (approximately 90 °C) and stirred for 24 h. After cooling to room temperature, the reaction mixture was slowly poured into ice-cold 1 M HCl (150 mL). Once gas evolution had ceased, the resulting precipitate was collected by vacuum filtration and washed successively with 1 M HCl (3 × 50 mL) and deionized water (3 × 50 mL). The product was dried under vacuum overnight to afford dibenzosuberenone oxime as a white solid (15.38 g, 95.5% yield).

$^1$H NMR (500 MHz, $CDCl_3$): δ 9.42 (br s, 1H), 7.69 (dd, J = 7.0, 1.2 Hz, 1H), 7.61 (m, 1H), 7.47-7.35 (m, 6H), 6.93 (m, 2H).

$^{13}$C NMR (126 MHz, $CDCl_3$): δ 156.37, 135.31, 134.42, 133.65, 130.65, 130.66, 130.51, 130.38, 128.81, 128.70, 127.66, 127.53.

1.2 Synthesis of Dibenzo[c,e]azepine N-Oxide

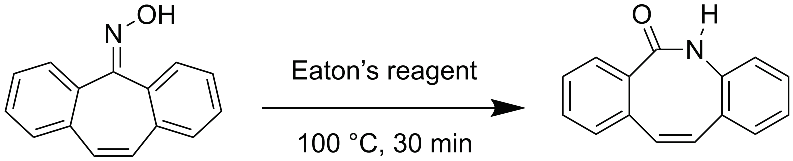

Dibenzosuberenone oxime (12.5 g, 56.5 mmol) was added to a round-bottom flask equipped with a magnetic stir bar. Eaton's reagent (25 mL) was added, and the reaction mixture was purged with argon and heated to 100 °C with stirring. Upon heating, the solid dissolved, and the reaction mixture gradually darkened. After stirring at 100 °C for 30 min, the hot reaction mixture was carefully poured onto crushed ice (65 g) to quench the reaction and stirred with a glass rod until the ice had completely melted. A gray precipitate formed immediately and was collected by vacuum filtration. The solid was washed thoroughly with deionized water (10 × 50 mL) until the filtrate reached a constant pH, then dried under vacuum overnight to afford the nitrone intermediate as a gray powder (10.55 g, quantitative yield).

$^{1}H$ NMR (500 MHz, $CDCl_3$): δ ppm 8.07 (br s, 1H), 7.51 (d, J = 7.8 Hz, 1H), 7.34 (t, J = 7.5 Hz, 1H), 7.29 (t, J = 7.3 Hz, 1H), 7.25-7.15 (m, 4H), 7.09 (d, J = 7.7 Hz, 1H), 6.99 (d, J = 11.5 Hz, 1H), 6.88 (d, J = 11.7 Hz, 1H).

$^{13}C$ NMR (126 MHz, $CDCl_3$): δ 173.78, 135.10, 135.00, 134.33, 134.13, 132.96, 129.73, 129.17, 128.29, 128.08, 128.03, 127.61, 127.12, 126.22.

### 1.3 Synthesis of Dibenzocyclooctene Intermediate

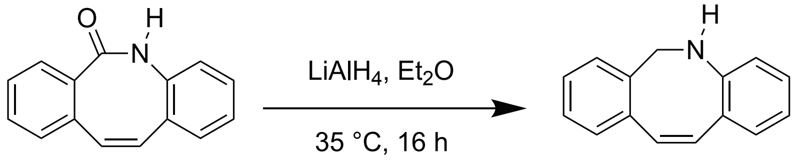

The nitrone intermediate (12.5 g, 56.5 mmol, 1.0 equiv.) was added to a 500 mL round-bottom flask equipped with a magnetic stir bar. Anhydrous diethyl ether (100 mL) was added, and the flask was cooled in an ice bath and purged with argon. Lithium aluminum hydride ($LiAlH_4$, 10.7 g, 282 mmol, 5.0 equiv.) was added portionwise to the stirred suspension. After complete addition, the ice bath was removed, the flask was fitted with a reflux condenser, and the reaction mixture was heated to reflux (35 °C) for 15 h under an argon atmosphere.

The reaction mixture was cooled to room temperature and then returned to an ice bath. Dichloromethane (100 mL) was added, followed by the slow addition of water (10 mL) with vigorous stirring. Aqueous NaOH (15 wt%, 11 mL) was then added, and the resulting aluminum salts were gently broken apart with a spatula. Additional dichloromethane (150 mL) and water (33 mL) were subsequently added, and the mixture was stirred while warming to room temperature. Anhydrous sodium sulfate was added, and the suspension was stirred for an additional 10–15 min before being filtered under vacuum. The filter cake was washed with ethyl acetate (approximately 300 mL total). The combined organic filtrates were concentrated, and the crude product was dissolved in dichloromethane (approximately 50 mL) and filtered to remove residual inorganic solids. Removal of the solvent under reduced pressure afforded the dibenzocyclooctene intermediate as a yellow powder (8.34 g, 71.3% yield).

$^{1}H$ NMR (500 MHz, $CDCl_3$): δ ppm 7.29 (m, 1H), 7.25–7.17 (m, 3H), 7.00 (d, J = 7.6 Hz, 1H), 6.91 (td, J = 7.6 Hz, 1.2 Hz, 1H), 6.63 (t, J = 7.4 Hz, 1H), 6.57 (d, J = 13.2 Hz, 1H), 6.49 (d, J = 8.1 Hz, 1H), 6.39 (d, J = 13.2 Hz, 1H), 4.61 (s, 2H), 4.31 (br s, 1H).

$^{13}C$ NMR (126 MHz, $CDCl_3$): δ 147.07, 139.19, 138.12, 134.67, 132.71, 130.10, 128.82, 127.92, 127.59, 127.34, 127.31, 121.72, 117.90, 117.68, 49.53

1.4 Synthesis of Dibenzocyclooctene Methyl Adipate Alkene Intermediate

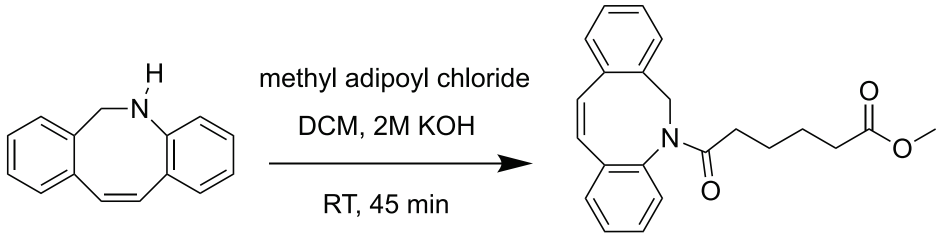

A $C_6$ spacer was incorporated to minimize steric hindrance and improve accessibility of the DBCO moiety during subsequent conjugation reactions (*1*). The dibenzocyclooctene intermediate (5.92 g, 28 mmol, 1.0 equiv.) was dissolved in dichloromethane (52 mL), followed by the addition of 2 M aqueous KOH (46.5 mL). Methyl adipoyl chloride (5.0 g, 28 mmol, 1.0 equiv.), dissolved in dichloromethane (14 mL), was added dropwise to the vigorously stirred biphasic reaction mixture over 30 min. The reaction mixture was then stirred at room temperature for an additional 45 min.

The organic layer was separated, and the aqueous phase was extracted with dichloromethane (3 × 25 mL). The combined organic extracts were dried over anhydrous $Na_2SO_4$, filtered, and concentrated under reduced pressure to afford the crude product as a yellow oil. Purification by flash column chromatography on silica gel using hexanes/ethyl acetate (3:1, v/v) afforded the dibenzocyclooctene methyl ester as a yellow oil (7.78 g, 79.5 % yield).

$^1$H NMR (500 MHz, $CDCl_3$): δ ppm 7.28 (m, 4H), 7.17 (m, 4H), 6.79 (d, J = 13.2 Hz, 1H), 6.60 (d, J = 13.2 Hz, 1H), 5.51 (d, J = 14.9 Hz, 1H), 4.23 (d, J = 14.9 Hz, 1H), 3.63 (s, 3H), 2.19 (t, J = 7.4 Hz, 2H), 2.06 (m, 1H), 1.89 (m, 1H), 1.53 (m, 2H), 1.44 (m, 2H).

$^{13}$C NMR (126 MHz, $CDCl_3$): δ 173.80, 172.16, 141.03, 136.18, 135.72, 134.78, 132.52, 132.01, 131.18, 130.29, 128.33, 128.05, 127.80, 127.25, 127.15, 126.89, 54.50, 51.37, 34.07, 33.66, 24.59, 24.27.

1.5 Synthesis of Dibenzocyclooctene Carboxylic Acid Intermediate

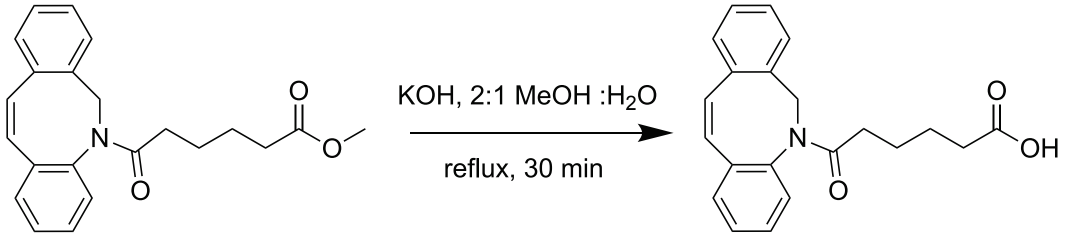

The dibenzocyclooctene methyl ester intermediate (7.75 g, 22.2 mmol, 1.0 equiv.) was dissolved in methanol (30 mL) in a 150 mL round-bottom flask. Separately, potassium hydroxide (KOH, 7.47 g, 133 mmol, 6.0 equiv.) was dissolved in Milli-Q water (15 mL). The aqueous KOH solution was added to the solution of the dibenzocyclooctene methyl ester intermediate with stirring. The flask was fitted with a reflux condenser, and the reaction mixture was heated to reflux for 30 min.

After cooling to room temperature, the reaction mixture was acidified to pH ≈ 2 with 1 M HCl. Dichloromethane (50 mL) was added, and the mixture was transferred to a separatory funnel. The organic layer was collected, and the aqueous phase was extracted with dichloromethane (2 × 25 mL). The combined organic extracts were washed with water (2 × 100 mL), dried over anhydrous $MgSO_4$, filtered, and concentrated under reduced pressure. The resulting residue was further dried under vacuum overnight to afford dibenzocyclooctene carboxylic acid as an off-white solid (5.4 g, 72.5 % yield).

$^1H$ NMR (500 MHz, $CDCl_3$): δ ppm 7.28 (m, 4H), 7.16 (m, 4H), 6.80 (d, J = 12.9 Hz, 1H), 6.60 (d, J = 13.2 Hz, 1H), 5.50 (d, J = 14.9 Hz, 1H), 4.24 (d, J = 14.9 Hz, 1H), 2.21 (t, J = 7.4 Hz, 2H), 2.07 (m, 1H), 1.90 (dt, J = 15.3, 7.4 Hz, 1H), 1.55 (m, 2H), 1.44 (m, 2H).

$^{13}C$ NMR (126 MHz, $CDCl_3$): δ 178.72, 172.49, 140.91, 136.17, 135.76, 134.66, 132.60, 132.00, 131.20, 130.34, 128.39, 128.04, 127.92, 127.29, 127.19, 126.97, 54.58, 34.08, 33.59, 24.45, 24.04.

1.6 Synthesis of DBCO–COOH

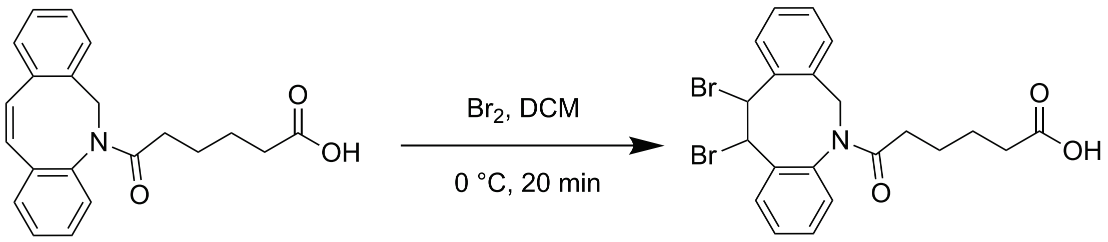

The dibenzocyclooctene carboxylic acid intermediate (4.2 g, 12.5 mmol) was dissolved in anhydrous dichloromethane (50 mL) in a 150 mL round-bottom flask equipped with a magnetic stir bar. The flask was purged with argon and cooled to 0 °C using an ice bath. A solution of bromine (2.6 g, 16.25 mmol, 1.3 equiv.) in anhydrous dichloromethane (5 mL) was added dropwise under stirring. Upon completion of the addition, the reaction mixture was stirred at 0 °C for an additional 30 min.

The reaction was quenched by the addition of aqueous sodium thiosulfate solution (15 wt%, 15 mL) with vigorous stirring to remove excess bromine. The mixture was transferred to a separatory funnel, and the organic phase was collected. The aqueous phase was extracted with dichloromethane (3 × 50 mL), and the combined organic extracts were washed with brine (50 mL), dried over anhydrous $MgSO_4$, filtered, and concentrated under reduced pressure. The resulting residue was dried under vacuum for approximately 1 h to afford the crude dibrominated intermediate as a yellow foam (5.51 g) that was used without further purification.

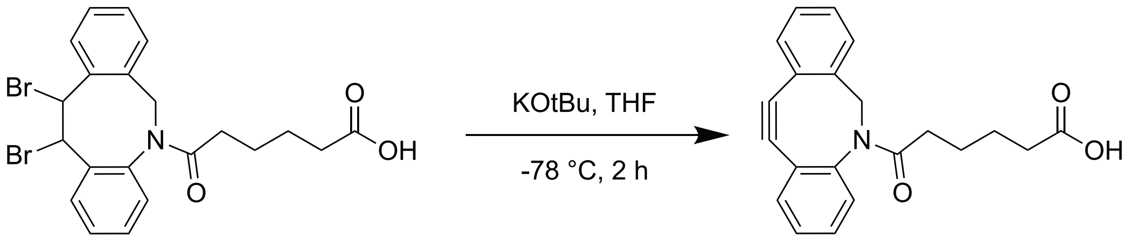

The dibrominated intermediate was dissolved in anhydrous THF (50 mL) in a 500 mL round-bottom flask equipped with a magnetic stir bar. The flask was purged with argon and cooled to −78 °C using a dry ice/isopropanol bath. A solution of potassium tert-butoxide (KOtBu, 1 M in THF, 40 mL) was added dropwise under stirring. The reaction mixture was maintained at −78 °C for 1.5 h, followed by stirring for an additional 30 min after completion of the addition. The mixture was then allowed to warm to room temperature and quenched with 1 M aqueous sodium bisulfate solution until the pH reached approximately 2.

Dichloromethane (100 mL) was added, and the organic phase was separated. The aqueous phase was extracted with dichloromethane, and the combined organic extracts were washed with water (2 × 50 mL) and brine (50 mL), dried over anhydrous $MgSO_4$, filtered, and concentrated under reduced pressure. The resulting residue was dissolved in diethyl ether (40–50 mL) and stored at −20 °C overnight to precipitate residual impurities. The precipitate was removed by filtration, and the filtrate was concentrated under reduced pressure. The crude product was dissolved in a minimal amount of dichloromethane and purified by silica gel chromatography using 5% MeOH in dichloromethane as the eluent. Product-containing fractions were combined and concentrated under reduced pressure to afford dibenzocyclooctyne carboxylic acid (DBCO-$C_6$-COOH) as a slightly orange solid (3.13 g, 75.1% yield).

$^1$H NMR (500 MHz, $CDCl_3$): δ ppm 7.70 (d, J = 7.6 Hz, 1H), 7.44-7.24 (m, 7H), 5.17 (d, J = 13.9 Hz, 1H), 3.67 (d, J = 13.7 Hz, 1H), 2.24 (m, 1H), 2.14 (t, J = 7.2 Hz, 2H), 1.92 (m, 1H), 1.52-1.31 (m, 4H)

$^{13}$C NMR (126 MHz, $CDCl_3$): δ 178.81, 173.16, 151.65, 147.90, 132.27, 128.87, 128.31, 128.26, 128.04, 127.68, 127.08, 125.45, 122.94, 122.53, 115.09, 107.66, 55.29, 34.32, 33.45, 24.53, 23.83

1.7 Synthesis of DBCO–NHS

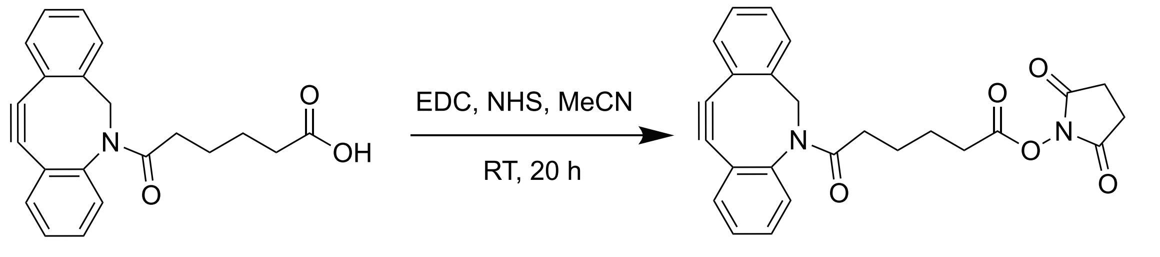

DBCO-$C_6$-COOH (3.0 g, 9.0 mmol) was dissolved in anhydrous acetonitrile (50 mL) and transferred to a 150 mL round-bottom flask equipped with a magnetic stir bar. *N*-hydroxysuccinimide (NHS, 1.55 g, 13.5 mmol, 1.5 equiv.) and EDC·HCl (2.59 g, 13.5 mmol, 1.5 equiv.) were added, and the reaction flask was purged with argon. The reaction mixture was stirred at room temperature for 20 h.

The solvent was removed under reduced pressure at 35 °C. The residue was dissolved in dichloromethane (50 mL), washed with water (3 × 50 mL) and brine (50 mL), dried over anhydrous $MgSO_4$, filtered, and concentrated under reduced pressure. The resulting yellow oil was further dried under high vacuum overnight to afford DBCO-$C_6$-NHS as a yellow solid (2.63 g, 67.6 % yield).

$^1$H NMR (500 MHz, $CDCl_3$): δ 7.71 (d, J = 7.6 Hz, 1H), 7.46–7.26 (m, 7H), 5.17 (d, J = 13.9 Hz, 1H), 3.68 (d, J = 13.9 Hz, 1H), 2.82 (m, 4H), 2.42 (td, J = 7.2 Hz, 3.4 Hz, 2H), 2.26 (m, 1H), 1.93 (m, 1H), 1.60–1.45 (m, 4H).

$^{13}C$ NMR (126 MHz, $CDCl_3$): δ 172.60, 169.04, 168.21, 151.66, 147.95, 132.22, 128.87, 128.36, 128.26, 128.04, 127.69, 127.10, 125.48, 122.91, 122.53, 115.08, 107.67, 55.27, 34.04, 30.44, 25.52, 25.50, 24.38, 23.68.

2. CO-O-NHS ester synthesis

(2-Cyclooctyn-1-yloxy) acetic acid was prepared following a published protocol (*3*) with some modification.

2.1 Synthesis of 8,8-Dibromobicyclo [5.1.0] octane

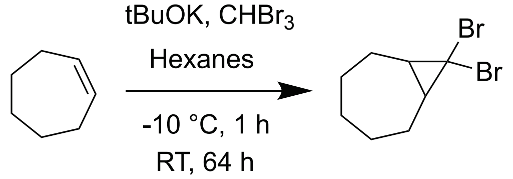

Potassium *tert*-butoxide (14.5 g, 130 mmol, 2.50 equiv.) was added to a 100 mL round-bottom flask equipped with a magnetic stir bar. Hexanes (9.8 mL) were added, followed by cycloheptene (6.10 mL, 52 mmol, 1.00 equiv.). The resulting suspension was stirred and cooled to −10 °C using a 1:1 (v/v) acetone/ice bath. Bromoform (6.8 mL, 78 mmol, 1.50 equiv.) was diluted with hexanes (10 mL) and added to the reaction mixture over 1 h using an addition funnel. Upon completion of the addition, the reaction temperature was gradually increased to room temperature over 3 h. The resulting suspension was then stirred at room temperature for 64 h. Water (39 mL) and 1 M HCl (50 mL) were added to the reaction mixture. The resulting biphasic mixture was transferred to a separatory funnel and diluted with hexanes (15 mL). The phases were separated, and the aqueous phase was extracted with hexanes (2 × 25 mL). The combined organic phases were dried over $Na_2SO_4$, filtered, and concentrated under a stream of dry air to afford 8,8-dibromobicyclo [5.1.0] octane as an oil (10.10 g, 73% mass recovery). The crude product was used directly in the subsequent step without further purification.

$^1H$ NMR (400 MHz, $CDCl_3$): δ 2.34 – 2.16 (m, 2H), 1.94 – 1.75 (m, 3H), 1.75 – 1.65 (m, 2H), 1.51 – 1.28 (m, 2H, overlap with water), 1.25 – 1.07 (m, 3H).

2.2 Synthesis of [(Z)-2-Bromo-2-cycloocten-1-yloxy] acetic Acid, Methyl Ester

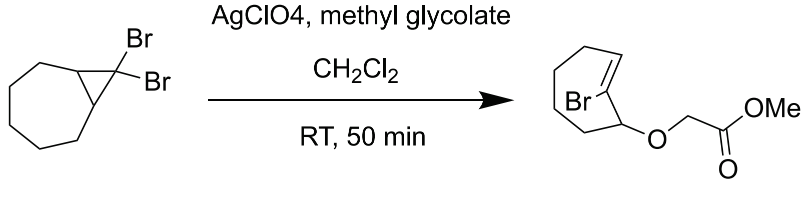

$CH_2Cl_2$ (20 mL) and 8,8-dibromobicyclo [5.1.0] octane (10.1 g, 27.9 mmol, 1.00 equiv.) were added to a 100 mL round-bottom flask equipped with a magnetic stir bar. Methyl glycolate (27 mL, 260 mmol, 9.3 equiv.) was then added. The reaction was performed at room temperature, and the flask was protected from light using aluminum foil. Silver perchlorate (15.7 g, 76 mmol, 2.00 equiv.) was added in a single portion. The resulting mixture was stirred at room temperature for 50 min, during which time a white precipitate formed. Saturated aqueous $NaHCO_3$ solution (65 mL) was added, followed by saturated aqueous NaCl solution (30 mL). The precipitated silver salts were filtered off and washed with methyl *tert*-butyl ether (MTBE, 40 mL). The filtrate was transferred to a separatory funnel and diluted with MTBE (20 mL). The phases were separated, and the organic phase was washed with water (2 × 20 mL), dried over $MgSO_4$, filtered, and concentrated in vacuo. The resulting crude product was obtained as an oil (9.6 g, 92% mass recovery) and used directly in the subsequent step without further purification.

$^1H$ NMR (400 MHz, $CDCl_3$): δ 6.21 (dd, J = 11.7, 4.2 Hz, 1H), 4.23 (d, J = 16.5 Hz, 1H), 4.17 – 4.06 (m, 1H), 3.97 (d, J = 16.4 Hz, 1H), 2.73 (qd, J = 11.9, 5.4 Hz, 1H), 2.34 – 2.19 (m, 1H), 2.12 – 1.85 (m, 4H), 1.70 (m, J = 15.2, 7.3, 3.8, 1.5 Hz, 1H), 1.54 – 1.41 (m, 1H), 1.35 – 1.23 (m, 1H), 0.93 – 0.72 (m, 1H).

2.3 Synthesis of (2-Cyclooctyn-1-yloxy) acetic Acid

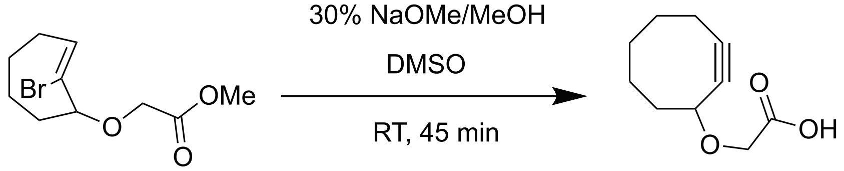

DMSO (10 mL) and [(Z)-2-bromo-2-cycloocten-1-yloxy] acetic acid methyl ester (5.0 g, 18 mmol, 1.00 equiv.) were added to a 100 mL round-bottom flask equipped with a magnetic stir bar. After the compound had completely dissolved, sodium methoxide (30% in methanol, 23 mL, 124 mmol, 6.9 equiv.) was added. The resulting mixture was stirred at room temperature for 45 min. The reaction mixture was then transferred to a separatory funnel and diluted with water (250 mL) and $CH_2Cl_2$ (100 mL). The phases were separated, and the aqueous phase was extracted with $CH_2Cl_2$ (2 × 50 mL). The combined organic phases were discarded. The aqueous phase was acidified to pH < 2 with 1 M HCl (200 mL) and then extracted with MTBE (50 mL). The resulting organic phase was washed with water (2 × 50 mL), dried over $MgSO_4$, filtered, and concentrated in vacuo to afford (2-cyclooctyn-1-yloxy) acetic acid as a tan solid (2.3 g, 70% yield).

$^1H$ NMR (400 MHz, $CDCl_3$): δ 4.39 (ddt, J = 7.1, 5.2, 2.1 Hz, 1H), 4.24 (d, J = 16.8 Hz, 1H), 4.09 (d, J = 16.8 Hz, 1H), 2.31 – 2.12 (m, 3H), 2.04 (m, J = 13.7, 9.0, 6.7, 1.2 Hz, 1H), 1.98 – 1.76 (m, 3H), 1.69 – 1.62 (m, 2H), 1.54 – 1.45 (m, 1H).

3. Nitrodopamine synthesis

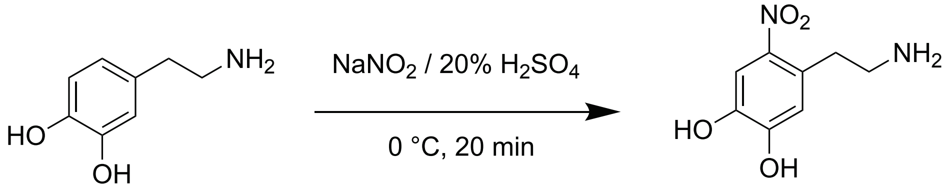

The synthesis of nitrodapamine was performed according to a published procedure (*4*) with some modifications. Dopamine (500 mg, 2.64 mmol) and $NaNO_2$ (630 mg, 9.13 mmol) were added to a 100 mL round-bottom flask, followed by Milli-Q water (15 mL). The mixture was stirred until the compounds were completely dissolved and then cooled in an ice bath. Sulfuric acid (20% v/v, 2.5

mL) was carefully added to the solution under vigorous stirring. After 20 min, the resulting precipitate was collected by vacuum filtration and washed three times with cold water, followed by three washes with cold MeOH, to afford nitrodopamine as a yellow solid (375 mg, 48% yield).

$^1$H NMR (400 MHz, DMSO) δ 7.44 (s, 1H), 6.56 (s, 1H), 3.08 – 2.98 (m, 4H).

### Supporting Figures

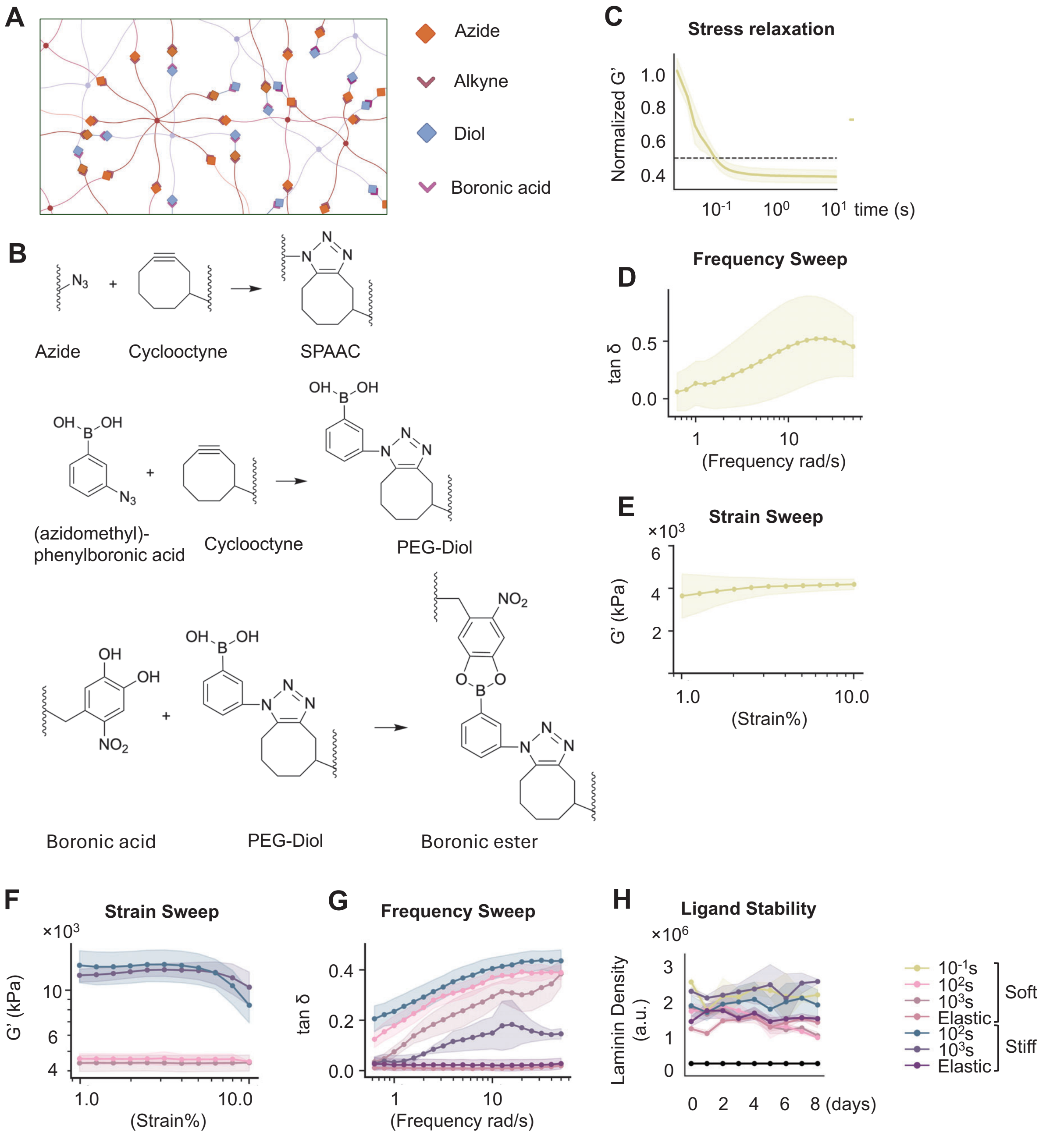

**Figure S1. Characterization of hydrogel chemical and mechanical properties and design of fast stress relaxing boronate ester hydrogels.** (**A**) Schematic and (**B**) chemical reactions illustrating the crosslinking strategy used to generate SPAAC stabilized boronate ester hydrogels. (**C**) Representative stress relaxation profile of the fast-relaxing boronate ester hydrogel with $\tau_{1/2}$ ~ $10^{-1}$ s, shown as normalized G′ over time. The dashed line indicates 50% of the initial normalized G′. (**D**) Frequency sweep measurement showing tan δ of the fast-relaxing boronate ester hydrogel.

(**E**) Strain sweep measurement showing G′ of the fast-relaxing boronate ester hydrogel. (**F**) Quantification of laminin density over 8 days on soft and stiff hydrogels with varying stress relaxation properties. (**G**) Frequency sweep measurements showing the loss tangent (tan $\delta$) of soft and stiff hydrogels with varying stress relaxation properties. (**H**) Strain sweep measurements showing the storage modulus (G′) of soft and stiff hydrogels with varying stress relaxation properties. Data are presented as mean ± s.d. from $n \geq 3$ independent replicates for hydrogel characterization.

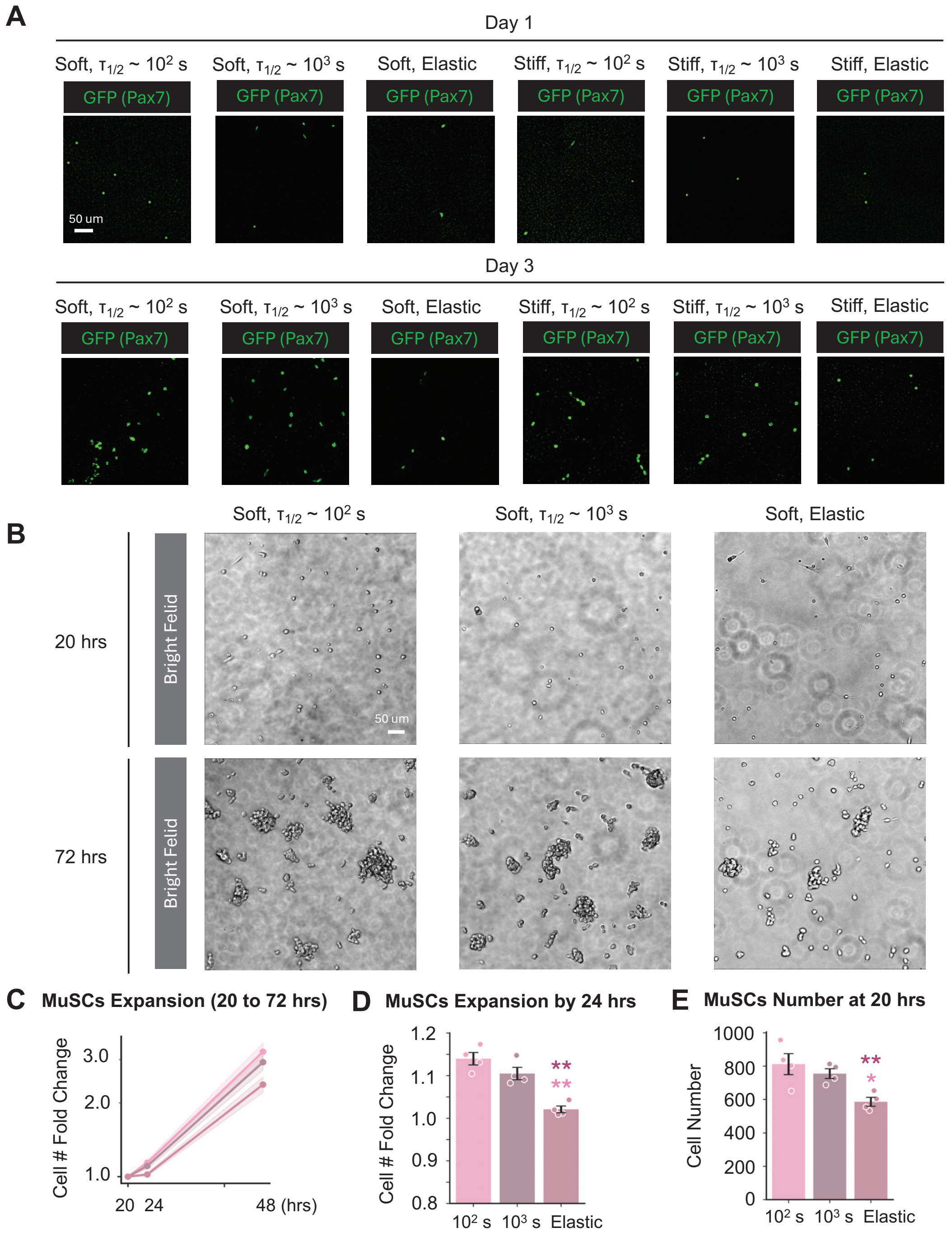

**Figure S2. Live-cell imaging and quantification of MuSC expansion on hydrogels with varying stiffness and stress-relaxation dynamics.** Representative live-cell fluorescence images of Pax7-EGFP reporter MuSCs cultured on soft or stiff hydrogels with varying stress relaxation properties at day 1 and day 3. GFP fluorescence indicates Pax7 expression in MuSCs. Scale bar, 50 µm. (**B**) Representative bright-field images of wild-type MuSCs cultured on soft hydrogels with varying stress relaxation properties at 20 and 72 hrs after cell seeding. Scale bar, 50 µm. (**C**) Quantification of fold change in MuSC number from 20 to 72 h, showing greater expansion on stress-relaxing hydrogels compared with elastic controls. (**D**) Quantification of MuSC expansion over the first 24 h, showing increased cell number on soft hydrogels with physiological stress relaxation ($\tau_{1/2}$ ~$10^2$ s) compared with slower-relaxing ($\tau_{1/2}$ ~$10^3$ s) and elastic hydrogels. (**E**) Quantification of MuSC number at 20 h after seeding, showing improved cell survival on stress-relaxing hydrogels relative to elastic controls. Data are presented as mean ± s.d. from $n$ = 4 independent experiment. Statistical significance was determined using pairwise Welch's $t$-tests with multiple-comparison correction. *P < 0.05 (*), P < 0.01 (**), and P < 0.001 (***).*

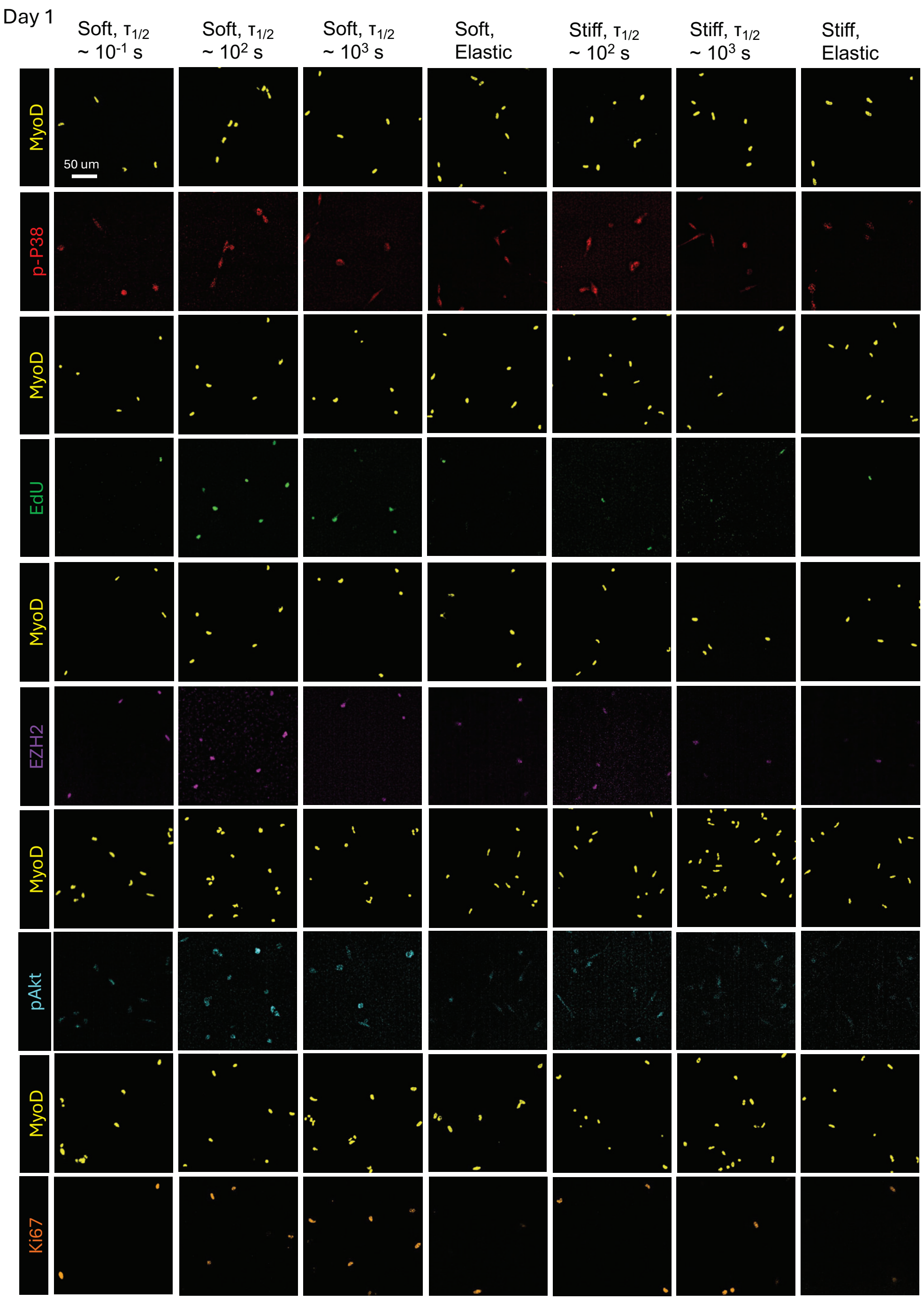

**Figure S3. Effects of matrix stiffness and stress relaxation on MuSC activation and proliferation at day 1.** Representative immunofluorescence images of MuSCs cultured for 1 day on soft or stiff hydrogels with varying stress relaxation properties, stained for MyoD together with phosphorylated p38 (p-p38), EdU, EZH2, phosphorylated Akt (pAkt), or Ki67. Scale bar, 50 µm.

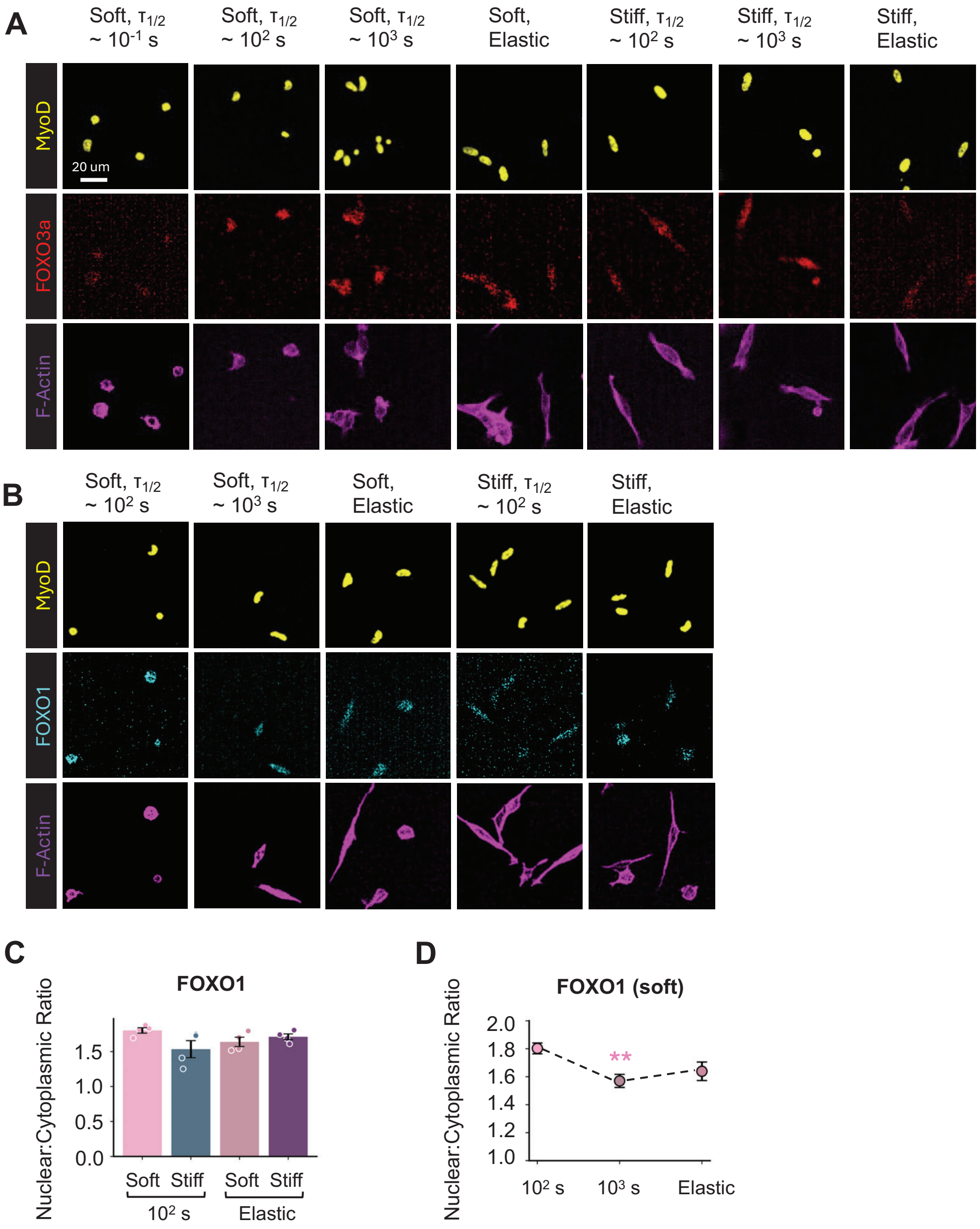

**Figure S4. Effects of matrix stiffness and stress relaxation on FOXO localization in MuSCs at day 1.** (**A**) Representative immunofluorescence images of MuSCs cultured for 1 day on soft or stiff hydrogels with varying stress relaxation properties, stained for MyoD, FOXO3a, and F-actin. Scale bar, 50 µm. (**B**) Representative immunofluorescence images of MuSCs cultured on soft or stiff hydrogels with varying stress relaxation properties, stained for MyoD, FOXO1, and F-actin. (**C**) Quantification of the nuclear-to-cytoplasmic ratio of FOXO1 in MuSCs cultured on soft and stiff hydrogels with $\tau_{1/2} \sim 10^2$ s or predominantly elastic behavior. (**D**) Quantification of the nuclear-to-cytoplasmic ratio of FOXO1 in MuSCs cultured on soft hydrogels with $\tau_{1/2} \sim 10^2$, $10^3$, and $\gg 10^4$ s. Each data point represents the mean nuclear-to-cytoplasmic translocation ratio calculated from $N \geq 300$ cells per hydrogel condition. Data are presented as mean ± s.d. from $n \geq 3$ independent biological replicates. Statistical significance was determined using pairwise Welch's *t*-tests with multiple-comparison correction. *$P < 0.05$ (*), $P < 0.01$ (**), and $P < 0.001$ (***)*.

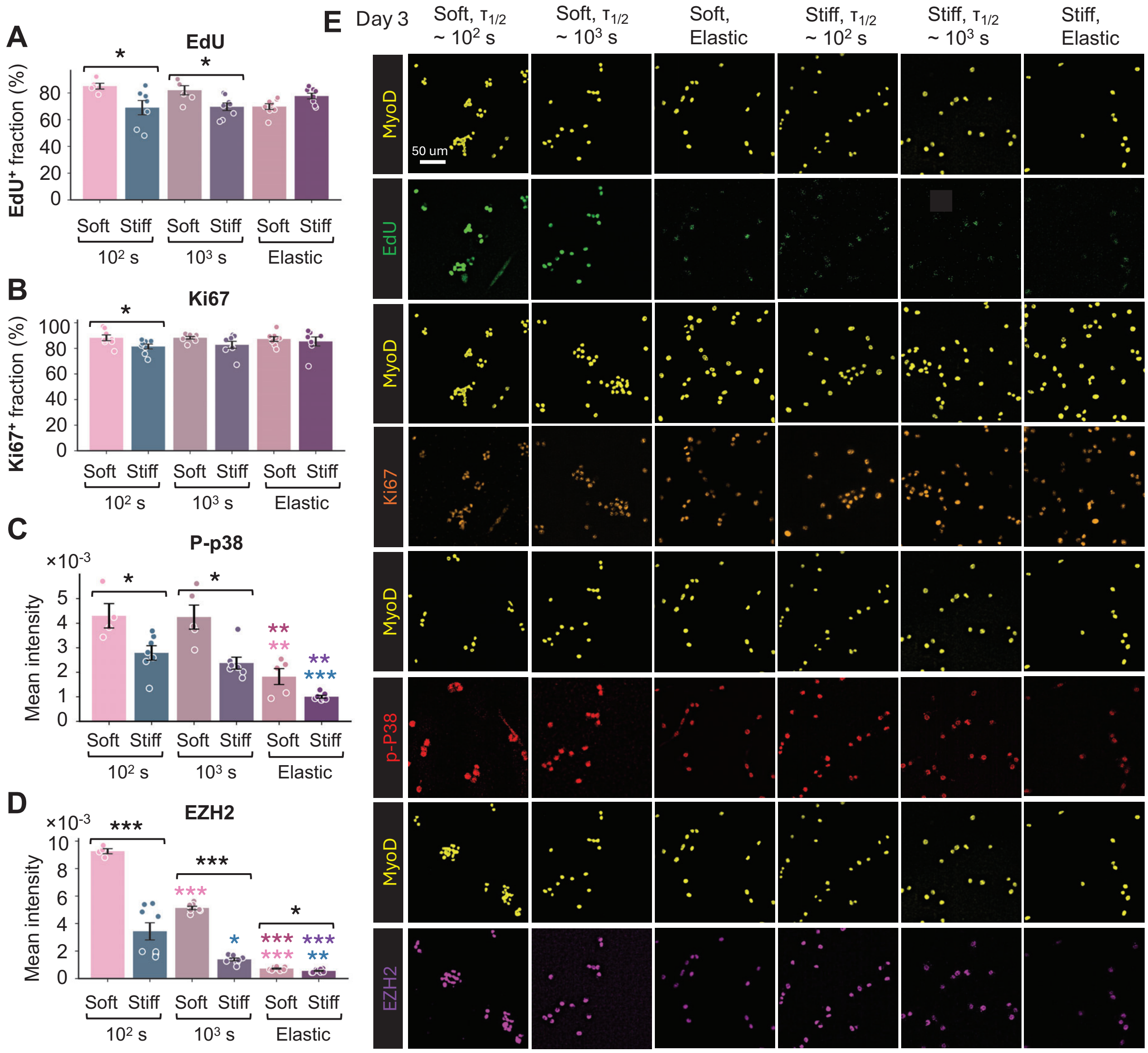

**Figure S5. Effects of matrix stiffness and stress relaxation on MuSC activation and proliferation at day 3.** Quantification of (**A**) EdU$^+$ fraction, (**B**) Ki67$^+$ fraction, (**C**) p-p38 mean intensity, (**D**) EZH2 mean intensity in MuSCs cultured for 3 days on soft or stiff hydrogels with varying stress relaxation properties. **(E)** Representative immunofluorescence images of MuSCs cultured under the indicated hydrogel conditions, stained for MyoD together with EdU, Ki67, p-p38, or EZH2. Scale bar, 50 µm. Each data point represents the fraction of positive cells (**A**, **B**) and mean fluorescence intensity (**C**, **D**) calculated from $N \geq 300$ cells per hydrogel condition. Each data point represents the fraction of positive cells (**A**, **B**), mean fluorescence intensity (**C**), or mean nuclear-to-cytoplasmic translocation ratio (**D**) calculated from $N \geq 300$ cells per hydrogel condition. Data are presented as mean ± s.d. from $n \geq 3$ independent biological replicates. Statistical significance was determined using pairwise Welch's *t*-tests with multiple-comparison correction. *P < 0.05 (*), P < 0.01 (**), and P < 0.001 (***).*

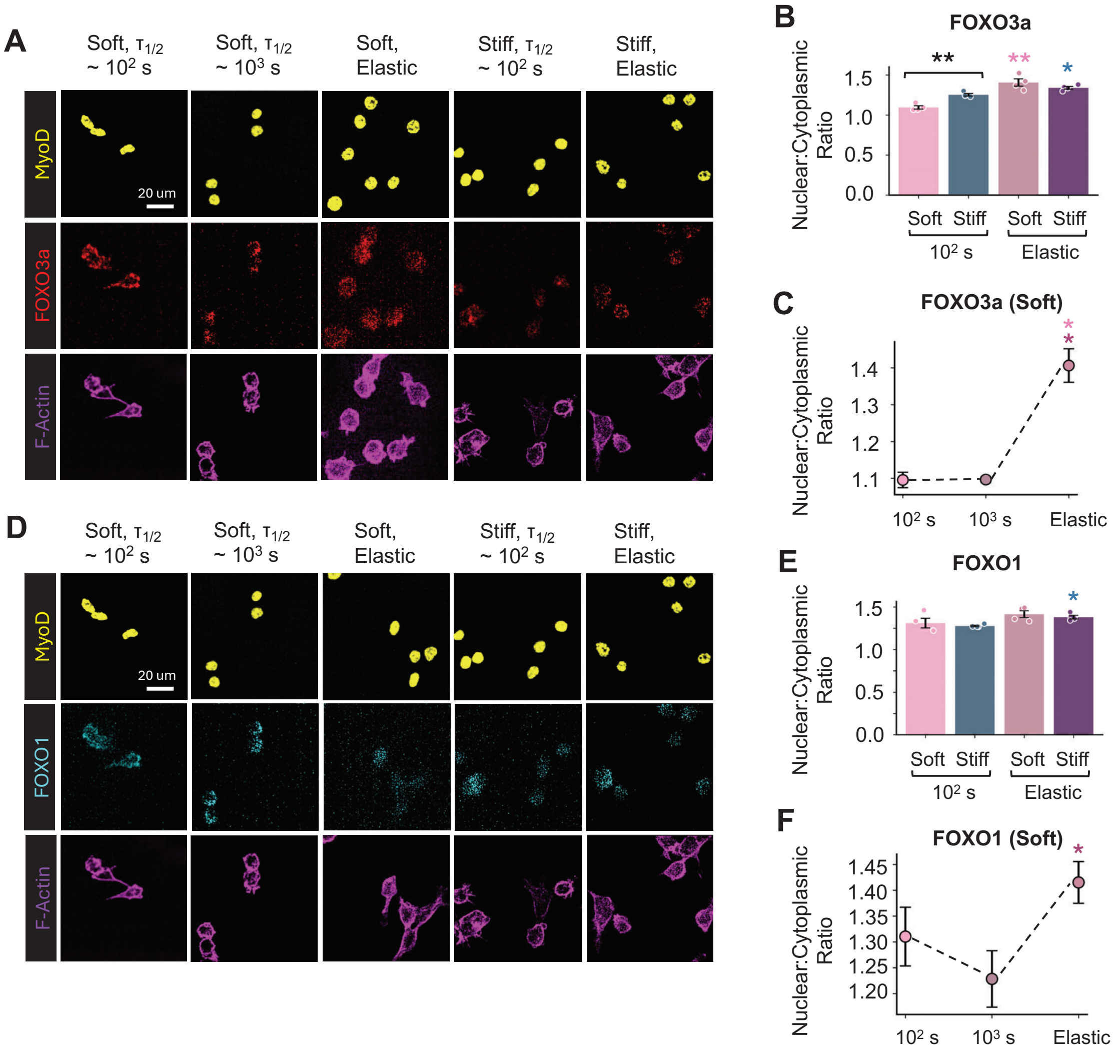

**Figure S6. Effects of matrix stiffness and stress relaxation on FOXO3a and FOXO1 localization in MuSCs at day 3.** (**A**) Representative immunofluorescence images of MuSCs cultured for 3 days on soft or stiff hydrogels with varying stress relaxation properties, stained for MyoD, FOXO3a, and F-actin. Scale bar, 50 µm. (**B**) Quantification of the nuclear-to-cytoplasmic ratio of FOXO3a in MuSCs cultured on soft and stiff hydrogels with $\tau_{1/2} \sim 10^2$ s or predominantly elastic behavior. (**C**) Quantification of the nuclear-to-cytoplasmic ratio of FOXO3a in MuSCs cultured on soft hydrogels with $\tau_{1/2} \sim 10^2$ s, $\tau_{1/2} \sim 10^3$ s, or predominantly elastic behavior. (**D**) Representative immunofluorescence images of MuSCs cultured for 3 days on soft or stiff hydrogels with varying stress relaxation properties, stained for MyoD, FOXO1, and F-actin. Scale bar, 50 µm. (**E**) Quantification of the nuclear-to-cytoplasmic ratio of FOXO1 in MuSCs cultured on soft and stiff hydrogels with $\tau_{1/2} \sim 10^2$ s or predominantly elastic behavior. (**F**) Quantification of the nuclear-to-cytoplasmic ratio of FOXO1 in MuSCs cultured on soft hydrogels with $\tau_{1/2} \sim 10^2$, $10^3$, and $\gg 10^4$ s. Each data point represents mean nuclear-to-cytoplasmic translocation ratio (**B**, **C**, **E**, **F**) calculated from $N \geq 300$ cells per hydrogel condition. Data are presented as mean ± s.d. from $n \geq 3$ independent biological replicates. Statistical significance was determined using pairwise Welch's *t*-tests with multiple-comparison correction. $P < 0.05$ (*), $P < 0.01$ (**), and $P < 0.001$ (***).

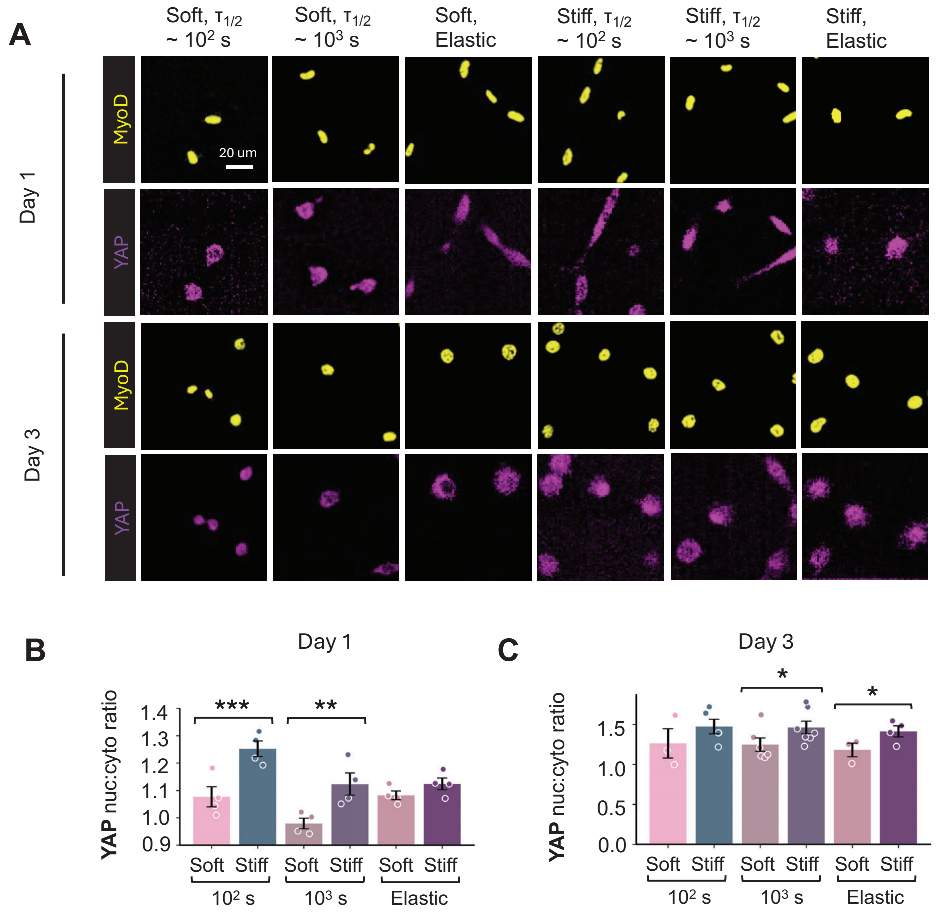

**Figure S7. Effects of matrix stiffness and stress relaxation on YAP localization in MuSCs at day 1 and day 3.** (**A**) Representative immunofluorescence images of MuSCs cultured for 1 or 3 days on soft or stiff hydrogels with varying stress relaxation properties, stained for MyoD and YAP. Scale bar, 50 µm. Quantification of the YAP nuclear-to-cytoplasmic ratio in MuSCs cultured on soft and stiff hydrogels with $\tau_{1/2} \sim 10^2$, $10^3$, and $\gg 10^4$ s at (**B**) day 1 and (**C**) day 3 of culture. Each data point represents mean nuclear-to-cytoplasmic translocation ratio (**B**, **C**) calculated from $N \geq 300$ cells per hydrogel condition. Data are presented as mean ± s.d. from $n \geq 3$ independent biological replicates. Statistical significance was determined using linear mixed-effects models (LMMs), with independent muscle batches included as a random effect. *$P < 0.05$ (*), $P < 0.01$ (**), and $P < 0.001$ (***).*

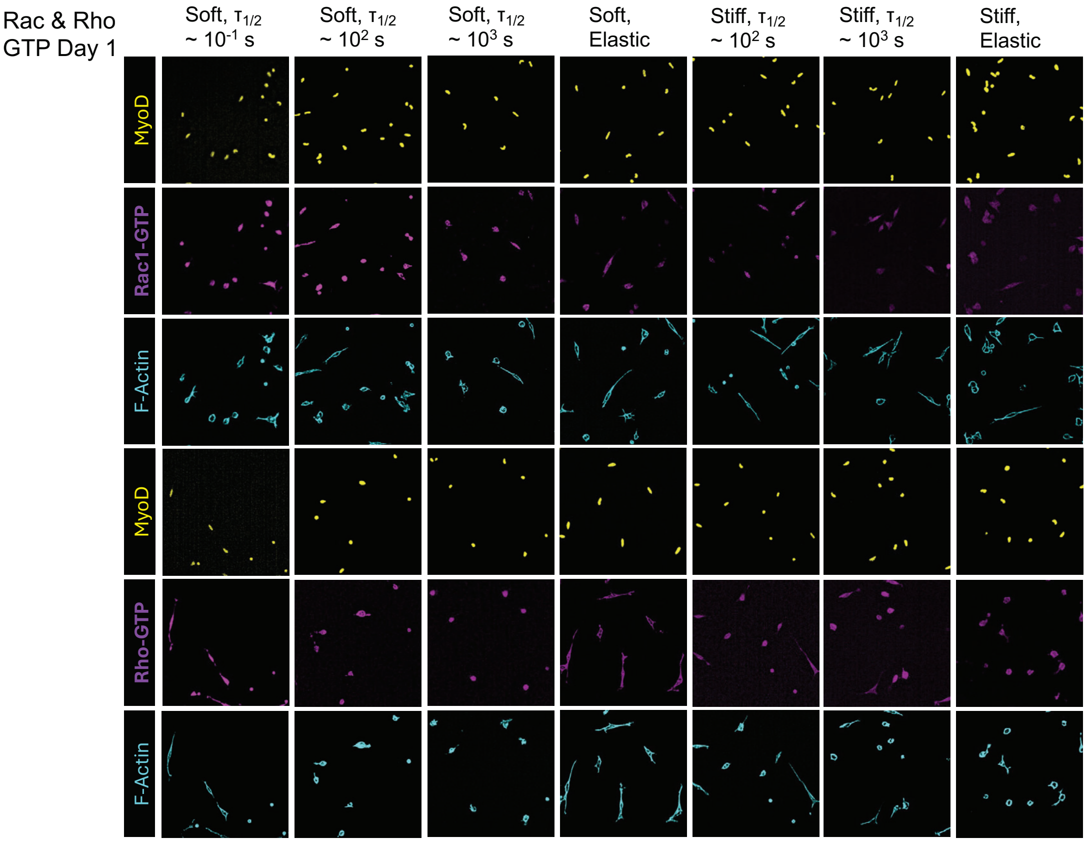

**Figure S8. Effects of matrix stiffness and stress relaxation on Rac/Cdc42 and Rho activity in MuSCs at day 1.** Representative immunofluorescence images of MuSCs cultured for 1 day on soft or stiff hydrogels with varying stress relaxation properties, stained for MyoD, active Rac1/Cdc42-GTP or Rho-GTP, and F-actin.

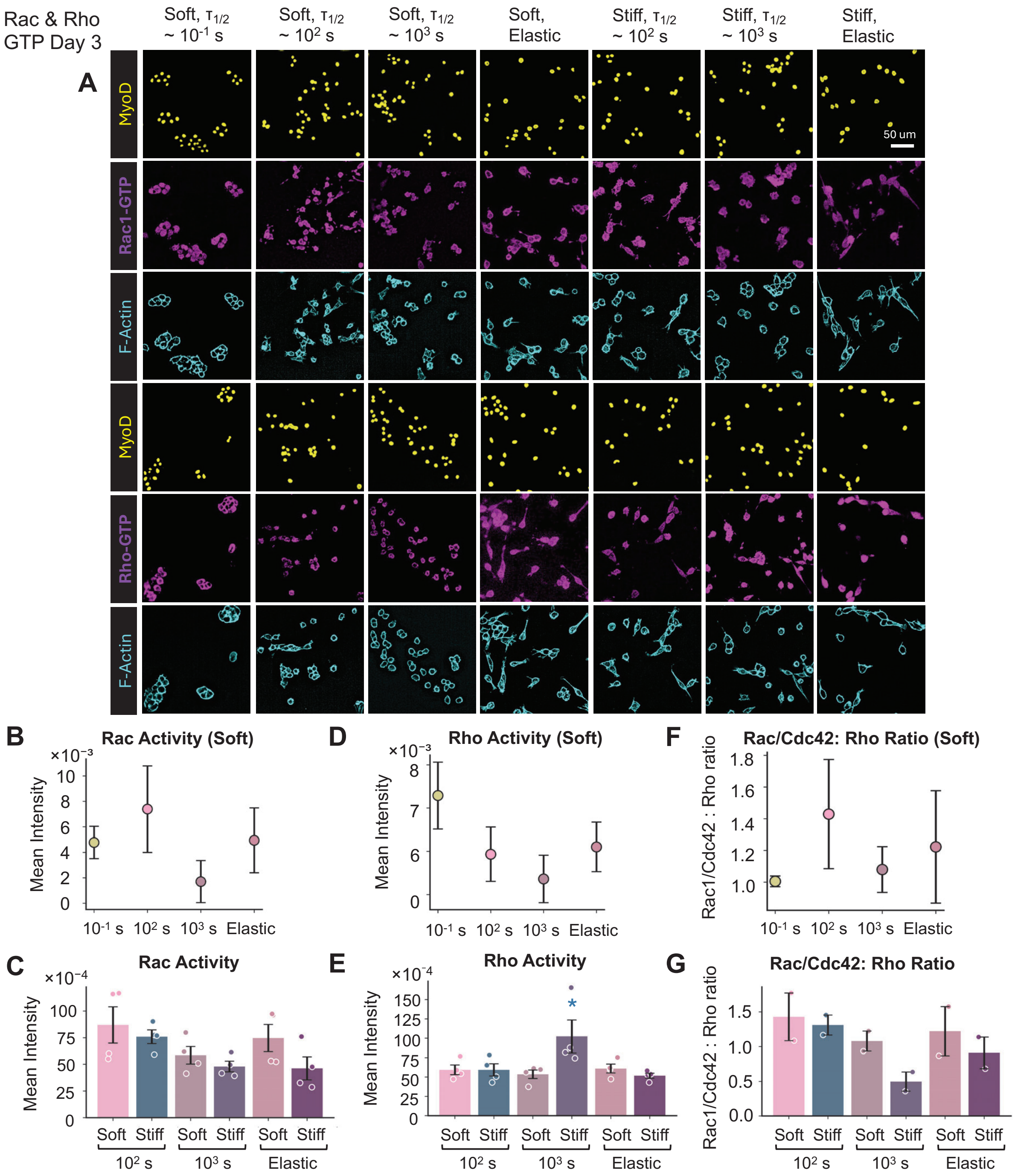

**Figure S9. Effects of matrix stiffness and stress relaxation on Rac1/Cdc42 and Rho GTPase activity in MuSCs at day 3.** (**A**) Representative immunofluorescence images of MuSCs cultured for 3 days on soft or stiff hydrogels with varying stress relaxation properties, stained for MyoD, active Rac1/Cdc42-GTP or Rho-GTP, and F-actin. Scale bar, 50 µm. (**B**) Quantification of Rac1/Cdc42 activity in MuSCs cultured on soft hydrogels with varying stress relaxation properties. (**C**) Quantification of Rac1/Cdc42 activity in MuSCs cultured on soft and stiff hydrogels with $\tau_{1/2}$ ~ $10^2$, $10^3$, and $\gg 10^4$ s. (**D**) Quantification of Rho activity in MuSCs cultured on soft hydrogels with varying stress relaxation properties. (**E**) Quantification of Rho activity in MuSCs cultured on soft and stiff hydrogels with $\tau_{1/2} \sim 10^2$, $10^3$, and $\gg 10^4$ s. (**F**) Rac1/Cdc42-to-Rho activity ratio in MuSCs cultured on soft hydrogels with varying stress relaxation properties. (**G**) Rac1/Cdc42-to-Rho activity ratio in MuSCs cultured on soft and stiff hydrogels with $\tau_{1/2} \sim 10^2$, $10^3$, and $\gg 10^4$ s. Each data point represents mean fluorescence intensity (**B**-**E**) calculated from $N \geq 300$ cells per hydrogel condition. Data are presented as mean ± s.d. from $n \geq 3$ independent biological replicates. Statistical significance was determined using pairwise Welch's *t*-tests with multiple-comparison correction. *$P < 0.05$ (*), $P < 0.01$ (**), and $P < 0.001$ (***).*

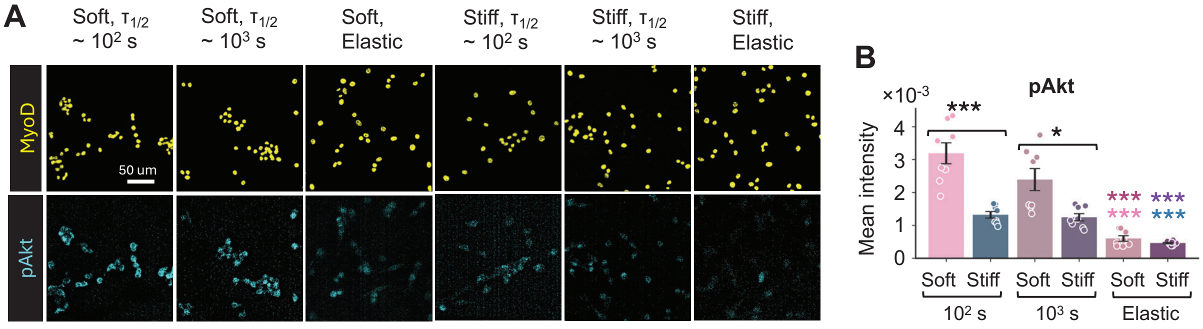

**Figure S10. Effects of matrix stiffness and stress relaxation on MuSC pAkt expression at day 3.** (**A**) Representative immunofluorescence images of MuSCs cultured under the indicated hydrogel conditions, stained for MyoD together with pAkt. Scale bar, 50 µm. (**B**) Quantification of pAkt mean intensity in MuSCs cultured for 3 days on soft or stiff hydrogels with varying stress relaxation properties. Each data point represents mean fluorescence intensity calculated from N ≥ 300 cells per hydrogel condition. Data are presented as mean ± s.d. from $n \geq 3$ independent biological replicates. Statistical significance was determined using pairwise Welch's *t*-tests with multiple-comparison correction. *$P < 0.05$ (*), $P < 0.01$ (**), and $P < 0.001$ (***).*

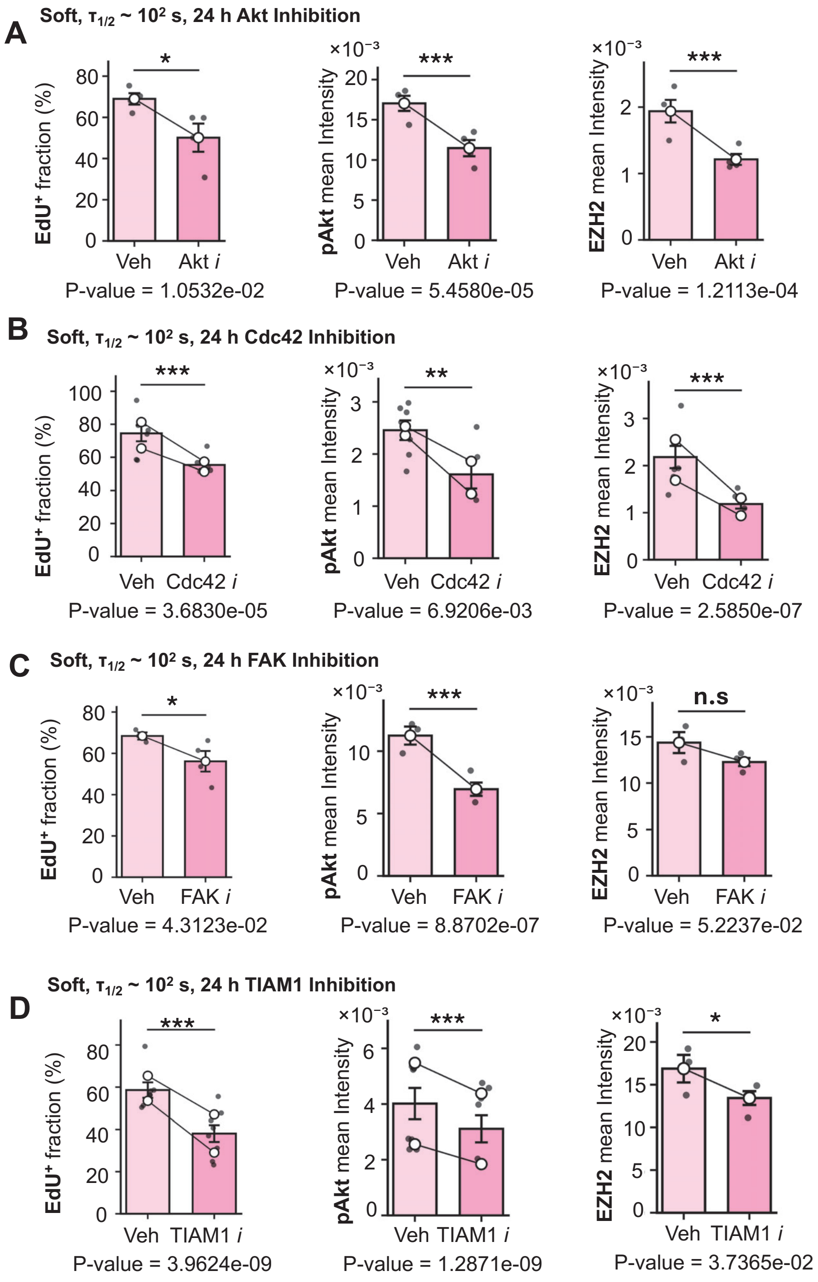

**Figure S11. Quantitative analysis of MuSC responses to inhibition of FAK-Rac/Cdc42-Akt signaling.** Quantification of $EdU^+$ fraction, pAkt mean intensity, and EZH2 mean intensity in MuSCs cultured on soft, viscoelastic hydrogels ($\tau_{1/2} \sim 10^2$ s) following 24 h treatment with (**A**) Akt inhibitor, (**B**) Cdc42 inhibitor, (**C**) FAK inhibitor, or (**D**) TIAM1 inhibitor, compared with the corresponding vehicle controls. Data are presented as mean ± s.d.; open circles represent the mean values of independent experiments, with lines connecting paired experimental replicates. Each data point represents the fraction of positive cells or mean intensity calculated from $N \geq 80$ cells per hydrogel condition. Statistical significance was determined using linear mixed-effects models (LMMs), with independent muscle batches included as a random effect. Exact P values are indicated in the figure; *P < 0.05 (*), P < 0.01 (**), and P < 0.001 (***)*, *n.s., not significant*.

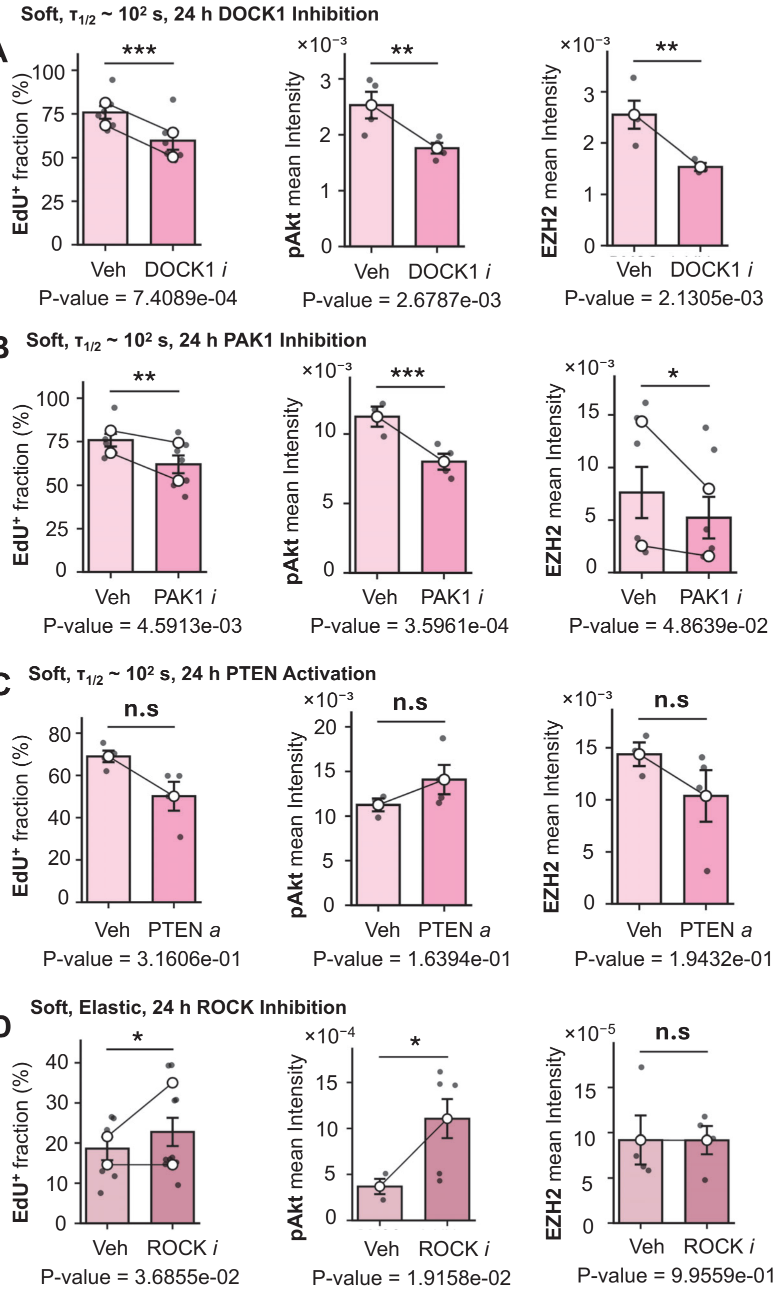

**Figure S12. Quantitative analysis of MuSC responses to modulation of Rac/Rho-PAK-Akt signaling.** Quantification of EdU$^+$ fraction, pAkt mean intensity, and EZH2 mean intensity in MuSCs cultured on soft, viscoelastic hydrogels ($\tau_{1/2} \sim 10^2$ s) following 24 h treatment with (**A**) DOCK1 inhibitor, (**B**) PAK1 inhibitor, or (**C**) PTEN activator, compared with the DMSO controls. (**D**) Quantification of EdU$^+$ fraction, pAkt mean intensity, and EZH2 mean intensity in MuSCs cultured on soft, predominantly elastic hydrogels following 24 h treatment with ROCK inhibitor compared with the vehicle control. Data are presented as mean ± s.e.m.; open circles represent the mean values of independent experiments, with lines connecting paired experimental replicates. Each data point represents the fraction of positive cells or mean intensity calculated from N ≥ 80 cells per hydrogel condition. Statistical significance was determined using linear mixed-effects models (LMMs), with independent muscle batches included as a random effect. Exact P values are indicated in the figure; *P < 0.05 (*), P < 0.01 (**), and P < 0.001 (***)*, *n.s., not significant*.

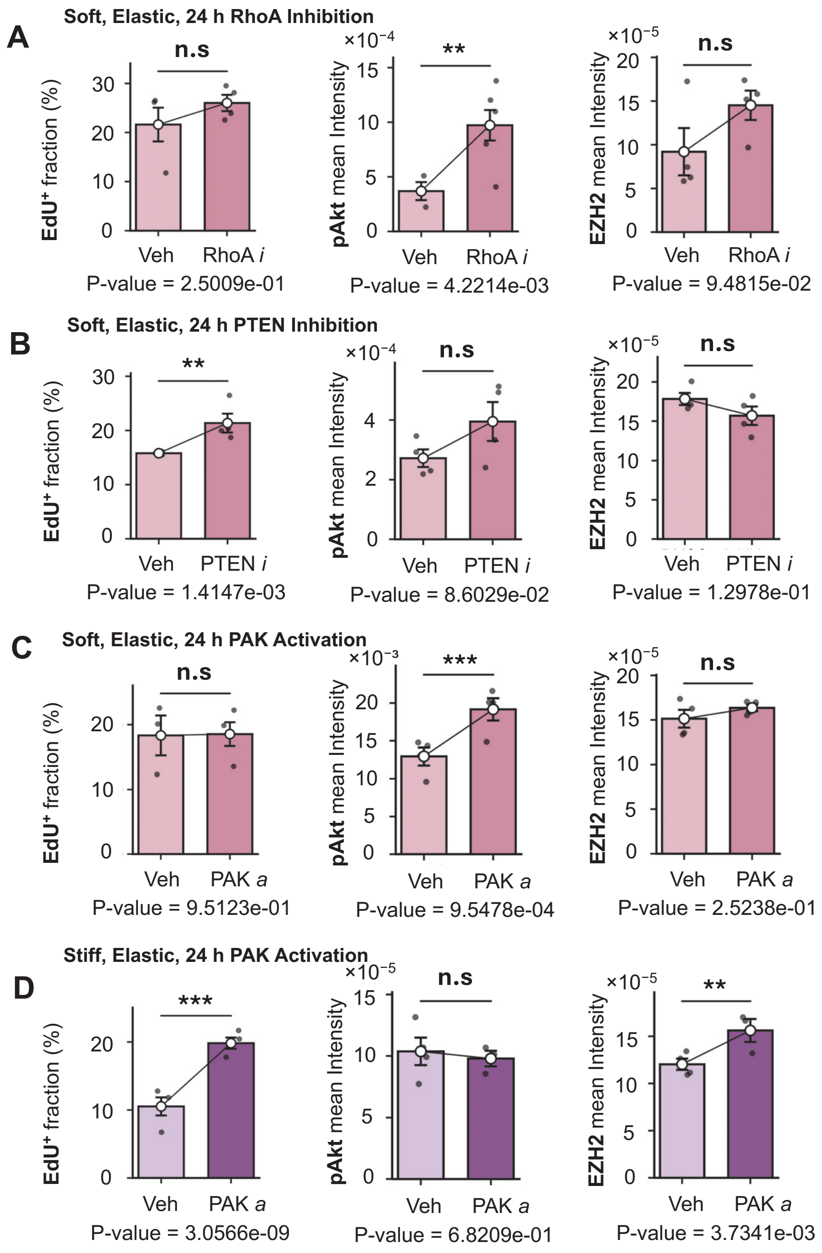

**Figure S13. Quantitative analysis of MuSC responses to modulation of RhoA-PTEN-PAK-Akt signaling.** Quantification of EdU⁺ fraction, pAkt mean intensity, and EZH2 mean intensity in MuSCs cultured on soft, predominantly elastic hydrogels following 24 h treatment with (**A**) RhoA inhibitor, (**B**) PTEN inhibitor, or (**C**) PAK activator, compared with the corresponding vehicle controls. (**D**) Quantification of EdU⁺ fraction, pAkt mean intensity, and EZH2 mean intensity in MuSCs cultured on stiff, predominantly elastic hydrogels following 24 h treatment with PAK activator compared with the vehicle control. Data are presented as mean ± s.e.m.; open circles represent the mean values of independent experiments, with lines connecting paired experimental replicates. Each data point represents the fraction of positive cells or mean intensity calculated from N ≥ 80 cells per hydrogel condition. Statistical significance was determined using linear mixed-effects models (LMMs), with independent muscle batches included as a random effect.

Exact P values are indicated in the figure; *P < 0.05 (*), P < 0.01 (**), and P < 0.001 (***)*, *n.s., not significant*.

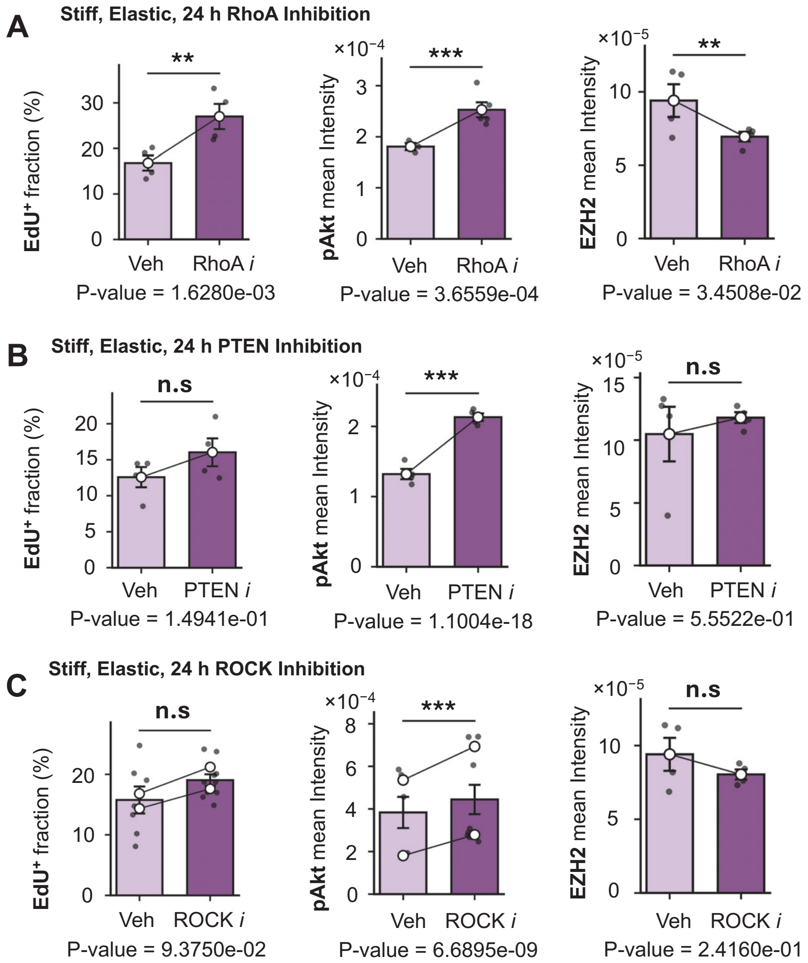

**Figure S14. Quantitative analysis of MuSC responses to modulation of RhoA-PTEN-ROCK-Akt signaling.** Quantification of EdU$^+$ fraction, pAkt mean intensity, and EZH2 mean intensity in MuSCs cultured on stiff, predominantly elastic hydrogels following 24 h treatment with (**A**) RhoA inhibitor, (**B**) PTEN inhibitor, or (**C**) ROCK inhibitor, compared with the corresponding vehicle controls. Data are presented as mean ± s.e.m.; open circles represent the mean values of independent experiments, with lines connecting paired experimental replicates. Each data point represents the fraction of positive cells or mean intensity calculated from $N \geq 80$ cells per hydrogel condition. Statistical significance was determined using linear mixed-effects models (LMMs), with independent muscle batches included as a random effect. Exact P values are indicated in the figure; *P < 0.05 (*), P < 0.01 (**), and P < 0.001 (***)*, *n.s., not significant*.

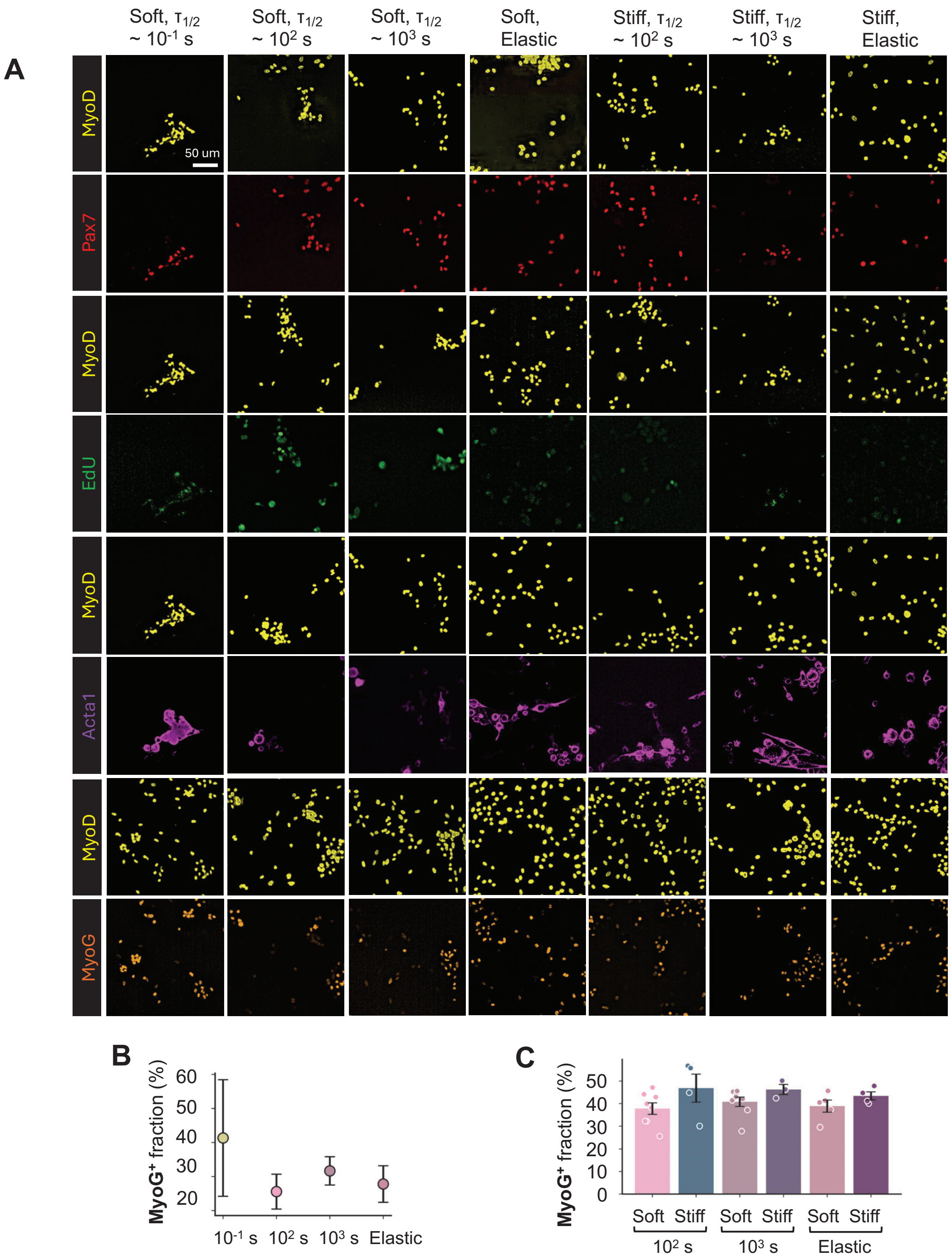

**Figure S15. Effects of matrix stiffness and stress relaxation on MuSC phenotype at day 7.** (**A**) Representative immunofluorescence images of MuSCs cultured for 7 days on soft or stiff hydrogels with varying stress relaxation properties, stained for MyoD together with Pax7, EdU, Acta1, or MyoG. Scale bar, 50 µm. (**B**) Quantification of the MyoG$^+$ fraction of MuSCs cultured on soft hydrogels with varying stress relaxation half-times ($\tau_{1/2} \sim 10^{-1}$, $10^2$, or $10^3$ s) or elastic hydrogels. (**C**) Quantification of the MyoG$^+$ fraction of MuSCs cultured on soft and stiff hydrogels with $\tau_{1/2} \sim 10^2$ s, $\tau_{1/2} \sim 10^3$ s, or elastic behavior. Each data point represents the fraction of positive cells calculated from $N \geq 300$ cells per hydrogel condition. Data are presented as mean ± s.d.; individual points represent independent experiments.

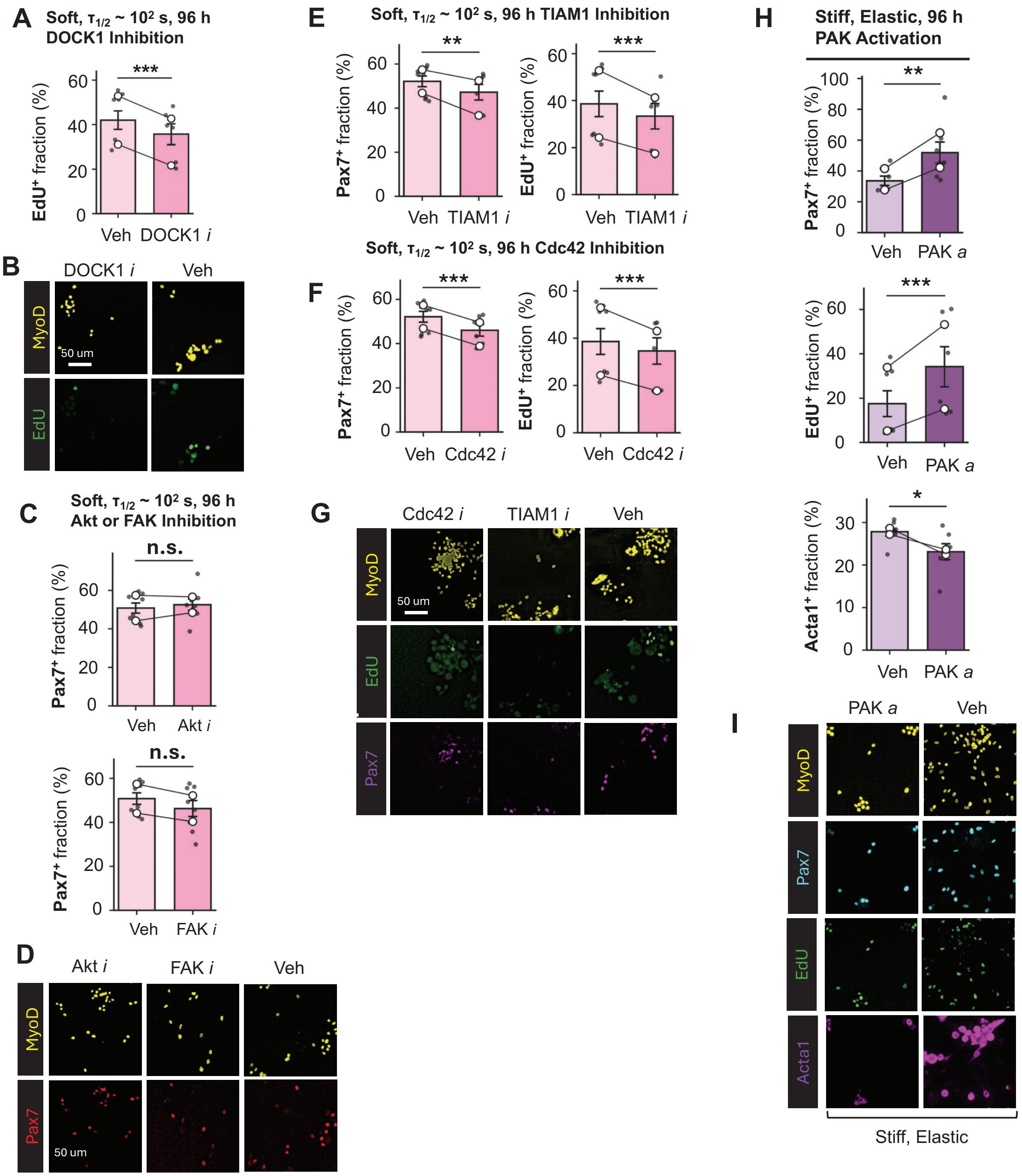

**Figure S16. Quantitative analysis of Day 7 immunofluorescence following 96 h pharmacological inhibition.** Cells cultured on soft, viscoelastic hydrogels (G' ~ 4 kPa, $\tau_{1/2}$~ $10^2$ s) or stiff (G' ~ 13 kPa), elastic hydrogels were treated with vehicle or the indicated small molecular drugs for 3 days, followed by washout and culture for an additional 4 days before immunofluorescence analysis on Day 7. (**A**, **B**) DOCK1 inhibition reduced proliferation (less EdU incorporation). (**C**, **D**) Transient Akt or FAK inhibition did not significantly alter the Pax7+ fraction on soft, viscoelastic hydrogels. (**E, F**) TIAM1 or Cdc42 inhibition reduced $Pax7^+$ and $EdU^+$ cell fractions on soft, viscoelastic hydrogels. (**G**) Representative immunofluorescence images following TIAM1 or Cdc42 inhibition on soft, viscoelastic hydrogels. (**H**) PAK1 activation on stiff elastic hydrogels increased Pax7$^+$ and EdU$^+$ fractions and decreased the Acta1$^+$ fraction. (**I**) Representative immunofluorescence images following PAK1 activation on stiff elastic hydrogels. Scale bars, 50 µm. Data are presented as mean ± s.d.; open circles represent the mean values of independent experiments, with lines connecting paired experimental replicates. Each data point represents the fraction of positive cells calculated from $N \geq 80$ cells per hydrogel condition. Statistical significance was determined using linear mixed-effects models (LMMs), with independent muscle batches included as a random effect. Exact P values are indicated in the figure; *P < 0.05 (*), P < 0.01 (**), and P < 0.001 (***)*, *n.s., not significant*.

NMR and FT-IR Spectra

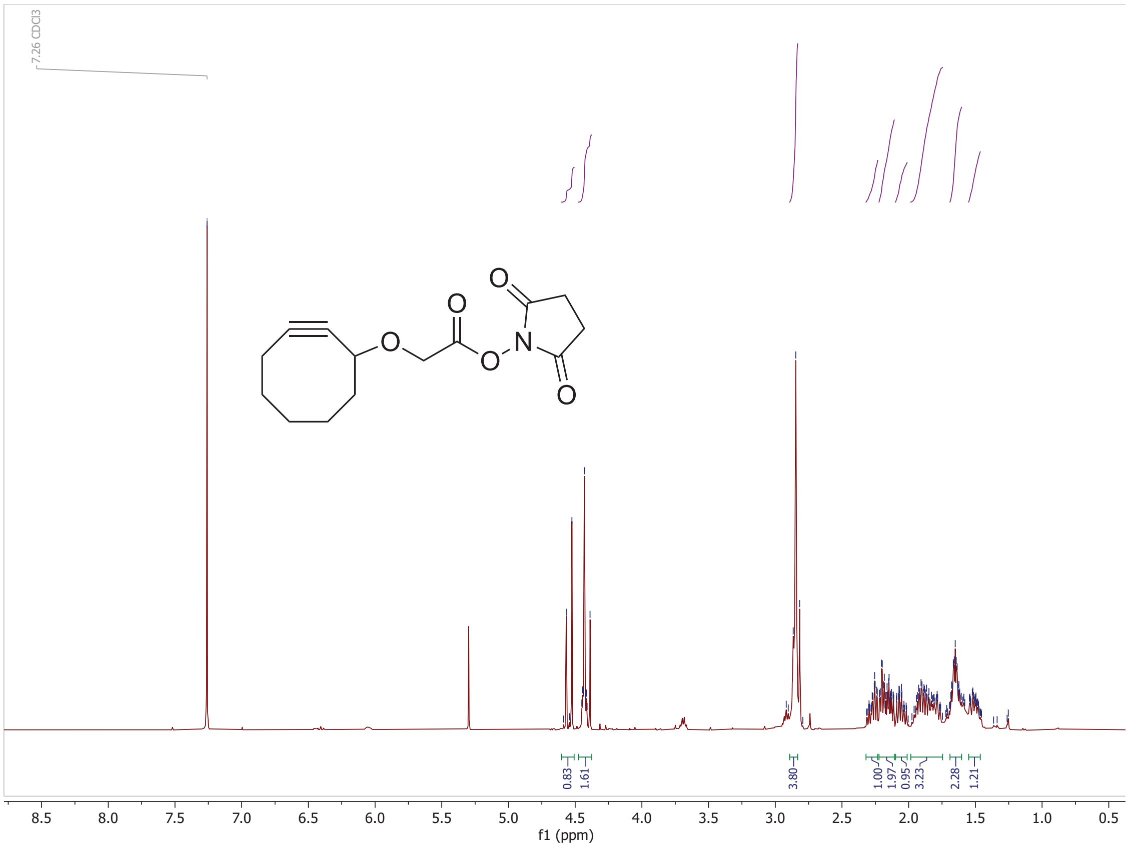

**Figure S17**. $^{1}$H NMR spectrum of cyclooctyne NHS ester (400 MHz, $CDCl_3$). δ 4.55 (d, J = 17.0 Hz, 1H), 4.47 – 4.38 (m, 2H), 2.86 (d, J = 7.9 Hz, 4H), 2.32 – 2.23 (m, 1H), 2.22 – 2.11 (m, 2H), 2.10 – 2.01 (m, 1H), 1.98 – 1.75 (m, 3H), 1.65 (m, J = 15.0, 10.1, 4.8, 2.7 Hz, 2H), 1.55 – 1.46 (m, 1H).

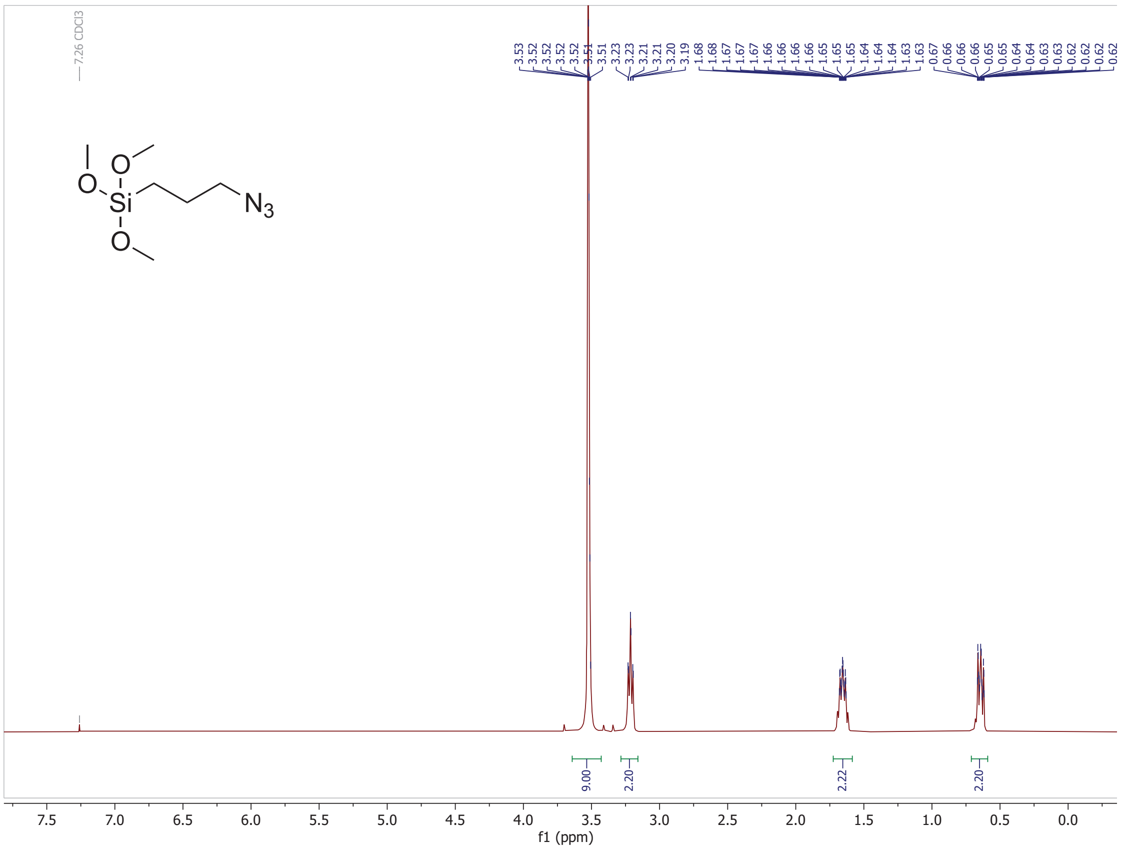

**Figure S18**. $^{1}$H NMR spectrum of (3-azidopropyl)trimethoxysilane. $^{1}$H NMR (400 MHz, $CDCl_3$) δ 3.52 (dt, J = 2.4, 1.3 Hz, 9H), 3.21 (td, J = 6.9, 1.5 Hz, 2H), 1.66 (m, J = 9.4, 5.8, 2.4, 1.2 Hz, 2H), 0.71 – 0.59 (m, 2H).

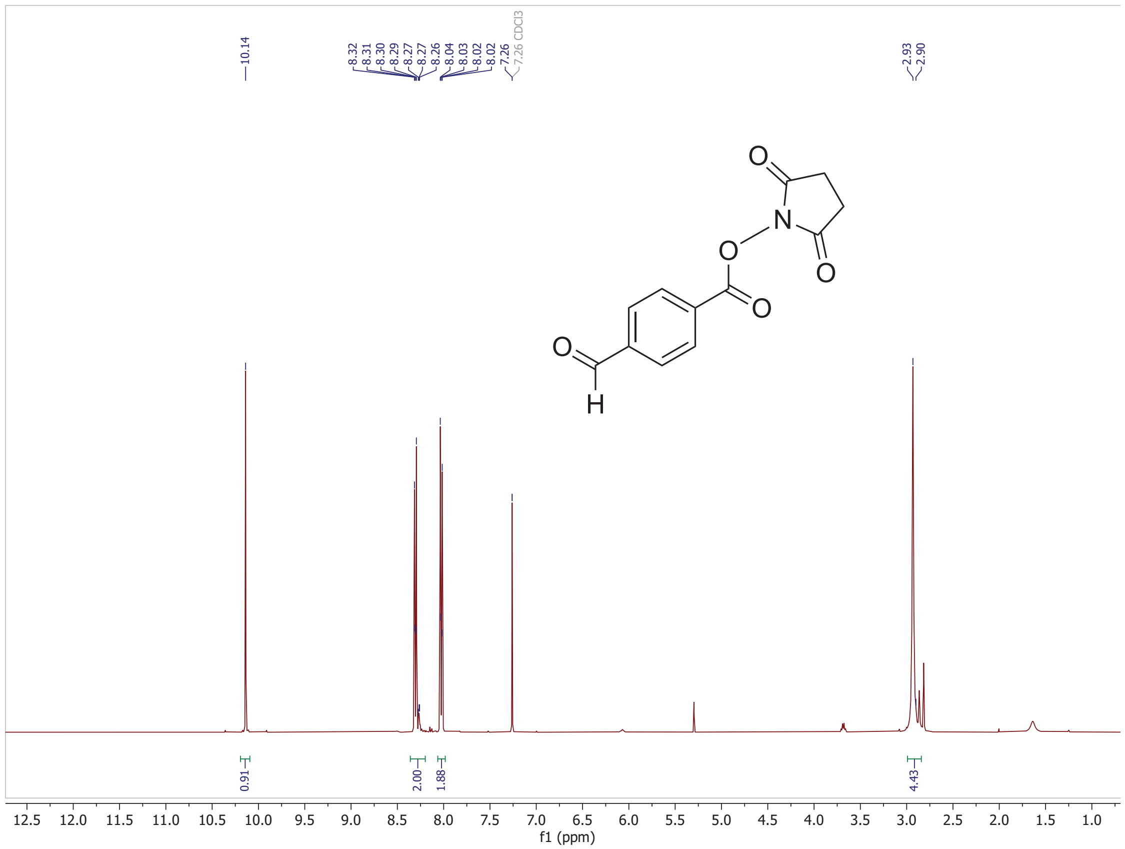

**Figure S19**. $^{1}$H NMR spectrum of *N*-succinimidyl 4-formylbenzoate. $^{1}$H NMR (400 MHz, $CDCl_3$) δ 10.14 (s, 1H), 8.36 – 8.20 (m, 2H), 8.06 – 7.98 (m, 2H), 2.93 (s, 4H).

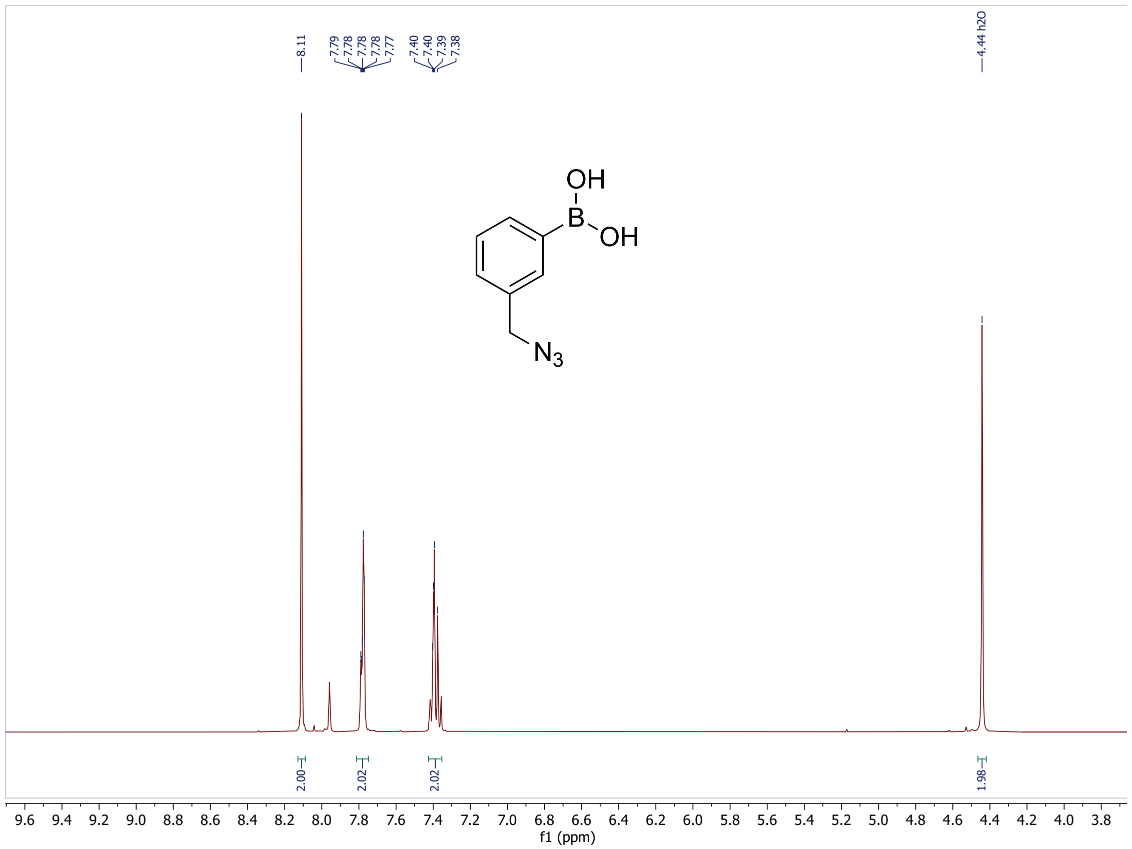

**Figure S20**. $^{1}H$ NMR spectrum of (3-(azidomethyl)phenyl)boronic acid. $^{1}H$ NMR (400 MHz, DMSO) δ 8.11 (s, 2H), 7.78 (dd, J = 4.6, 2.7 Hz, 2H), 7.42 – 7.35 (m, 2H), 4.44 (s, 2H).

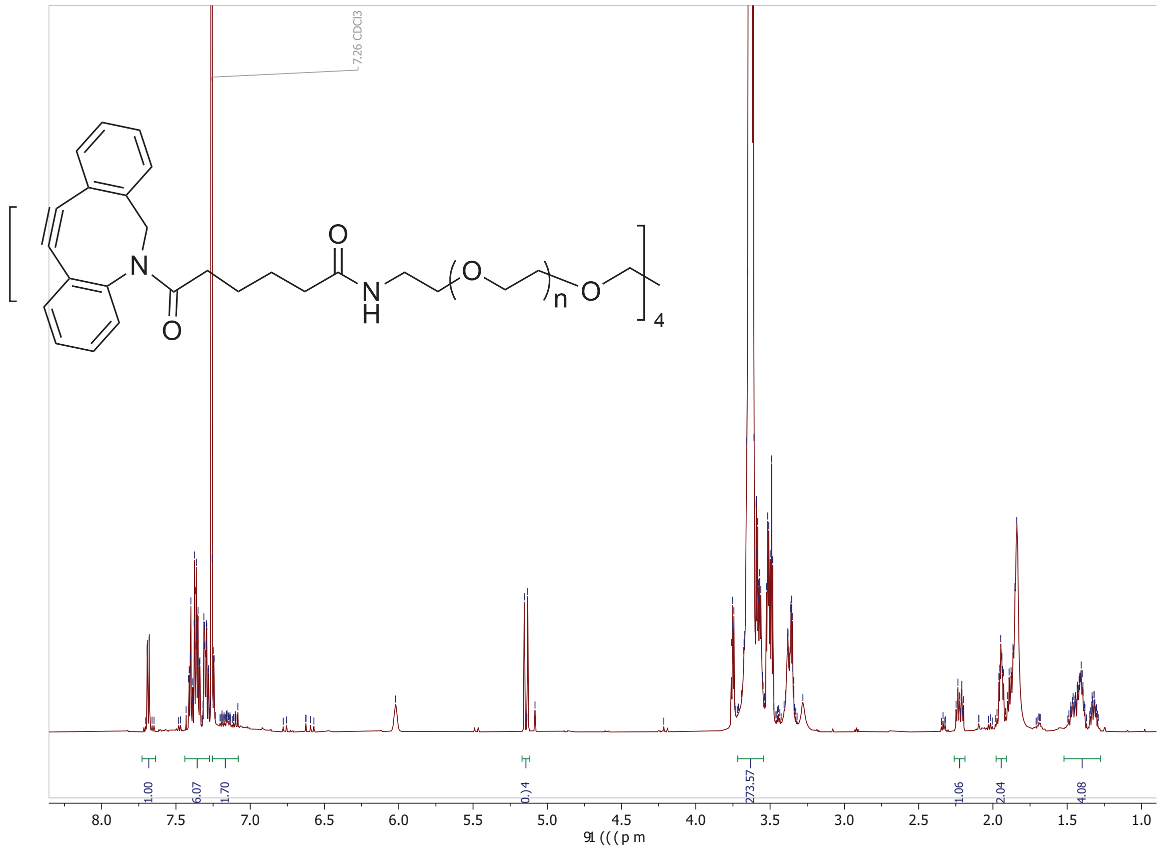

**Figure S21**. $^1$H NMR spectrum of DBCO-functionalized PEG. $^1$H NMR (600 MHz, $CDCl_3$) δ 7.69 (dd, J = 7.6, 1.2 Hz, 1H), 7.44 – 7.27 (m, 6H), 7.25 – 7.08 (m, 2H), 5.14 (d, J = 13.8 Hz, 1H), 3.64 (s, 282H), 2.26 – 2.19 (m, 1H), 1.98 – 1.91 (m, 2H), 1.52 – 1.28 (m, 4H). The end group conversion was determined to be 82% from $^1$H NMR analysis.

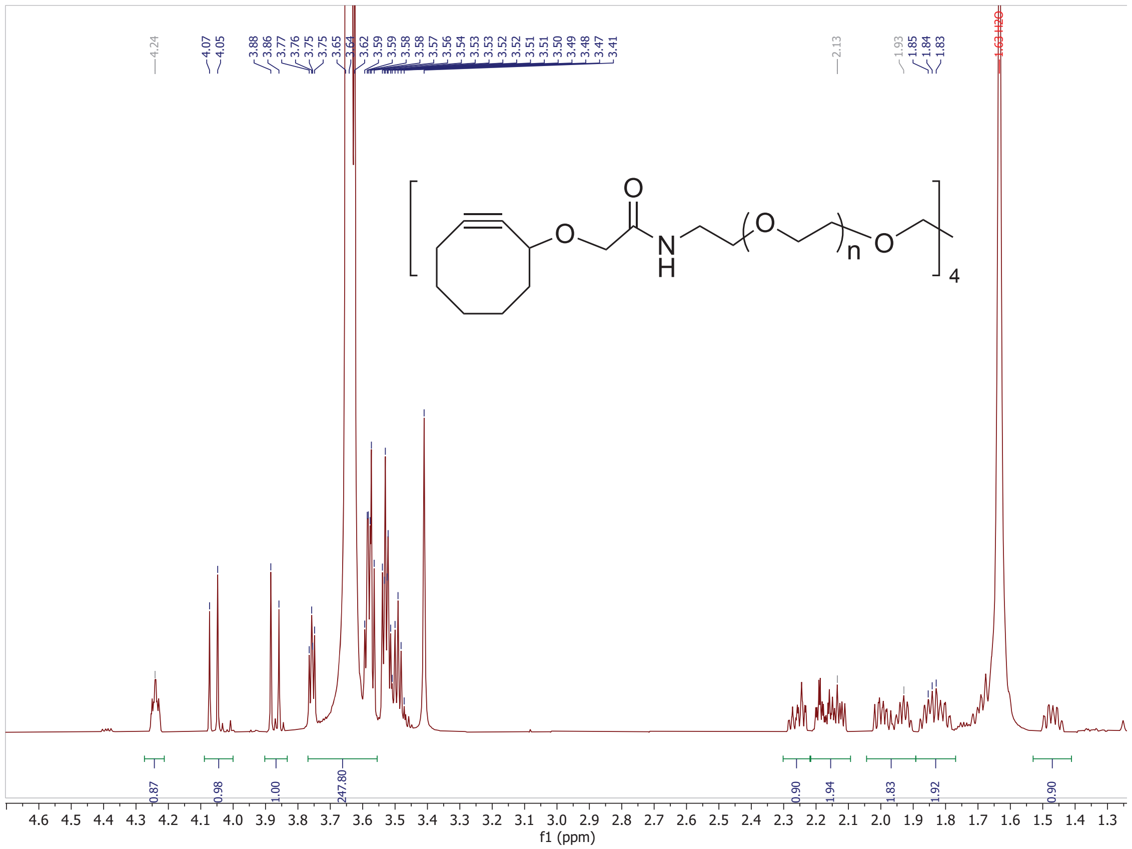

**Figure S22**. $^1$H NMR spectrum of cyclooctyne-functionalized PEG. $^1$H NMR (600 MHz, $CDCl_3$) δ 4.24 (m, J = 7.0, 4.8, 2.4 Hz, 1H), 4.06 (d, J = 15.1 Hz, 1H), 3.87 (d, J = 15.1 Hz, 1H), 3.64 (s, 241H), 2.26 (m, J = 16.9, 7.9, 6.1, 1.9 Hz, 1H), 2.22 – 2.09 (m, 2H), 2.04 – 1.89 (m, 2H), 1.89 – 1.77 (m, 2H), 1.47 (dt, J = 16.6, 9.0 Hz, 1H).The end group conversion was determined to be 95% from $^1$H NMR analysis.

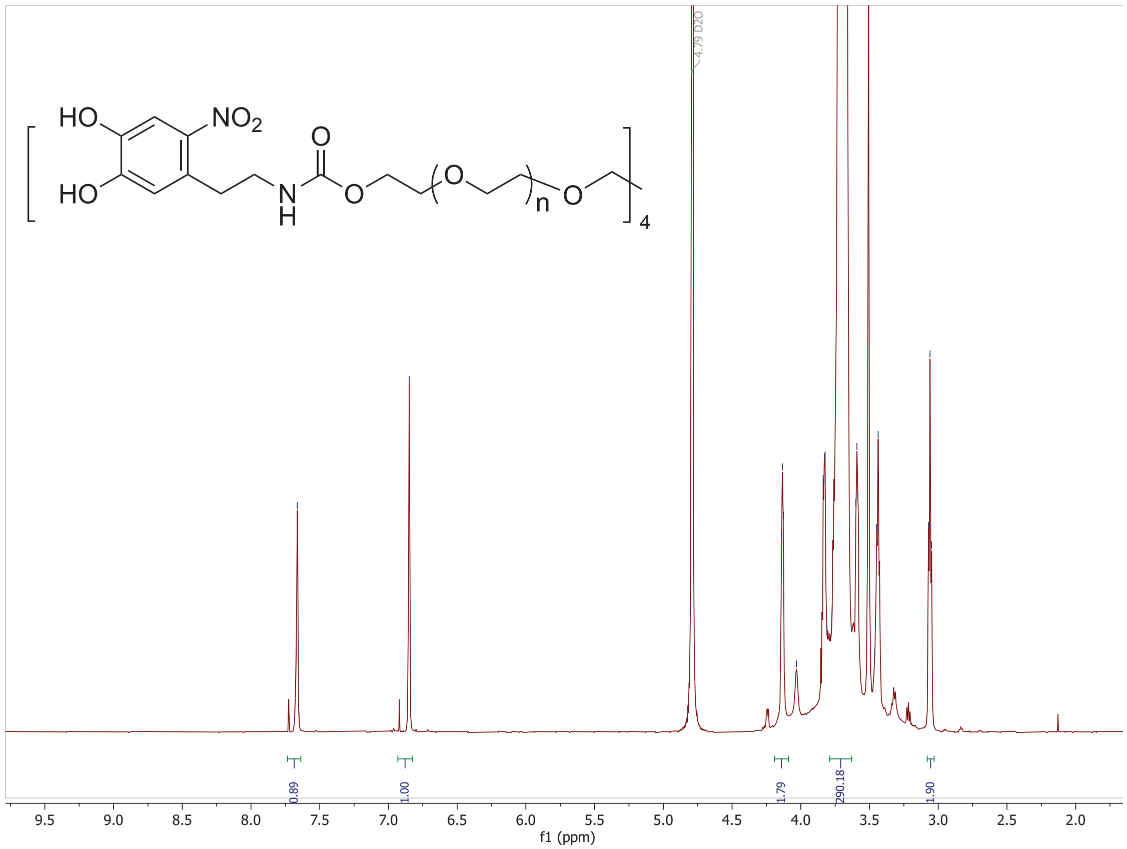

**Figure S23**. $^1$H NMR spectrum of nitrodopamine-functionalized PEG. $^1$H NMR (600 MHz, $D_2O$) δ 7.66 (s, 1H), 6.85 (s, 1H), 4.13 (t, J = 4.5 Hz, 2H), 3.85 – 3.81 (m, 2H), 3.71 (s, 290H), 3.60 – 3.43 (m, 6H), 3.06 (t, J = 6.6 Hz, 2H). The end group conversion was determined to be 78% from $^1$H NMR analysis.

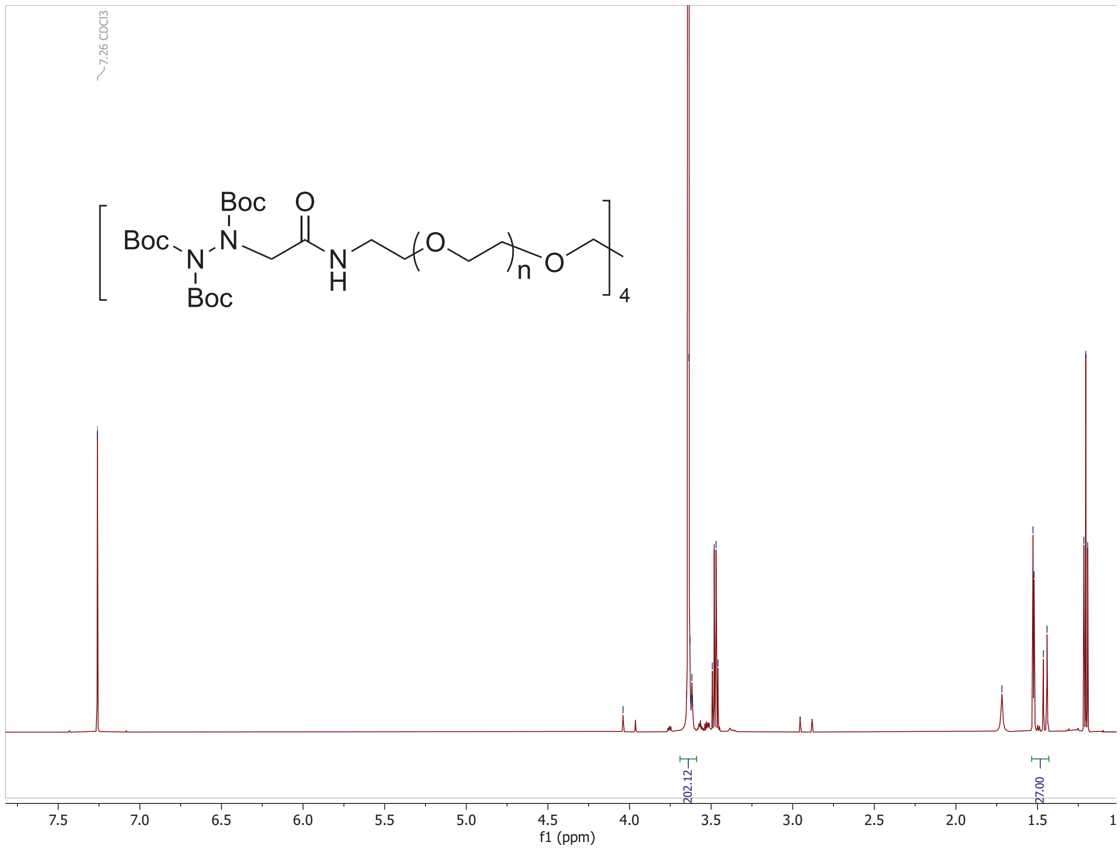

**Figure S24**. $^1$H NMR spectrum of Boc-protected hydrazine-functionalized PEG. $^1$H NMR (600 MHz, $CDCl_3$) δ 3.64 (s, 210H), 1.54 – 1.43 (m, 27H). The end group conversion was determined to be 100% from $^1$H NMR analysis.

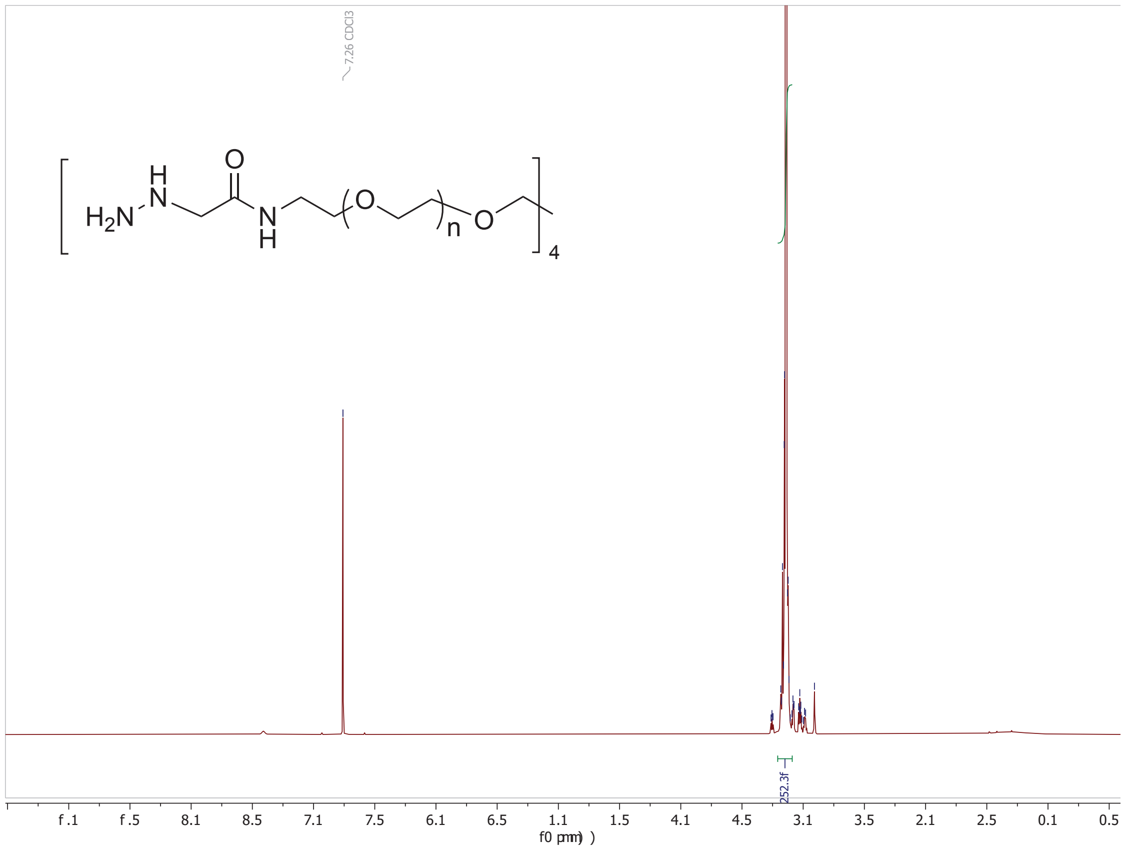

**Figure S25**. $^1H$ NMR spectrum of hydrazine-functionalized PEG (600 MHz, $CDCl_3$).

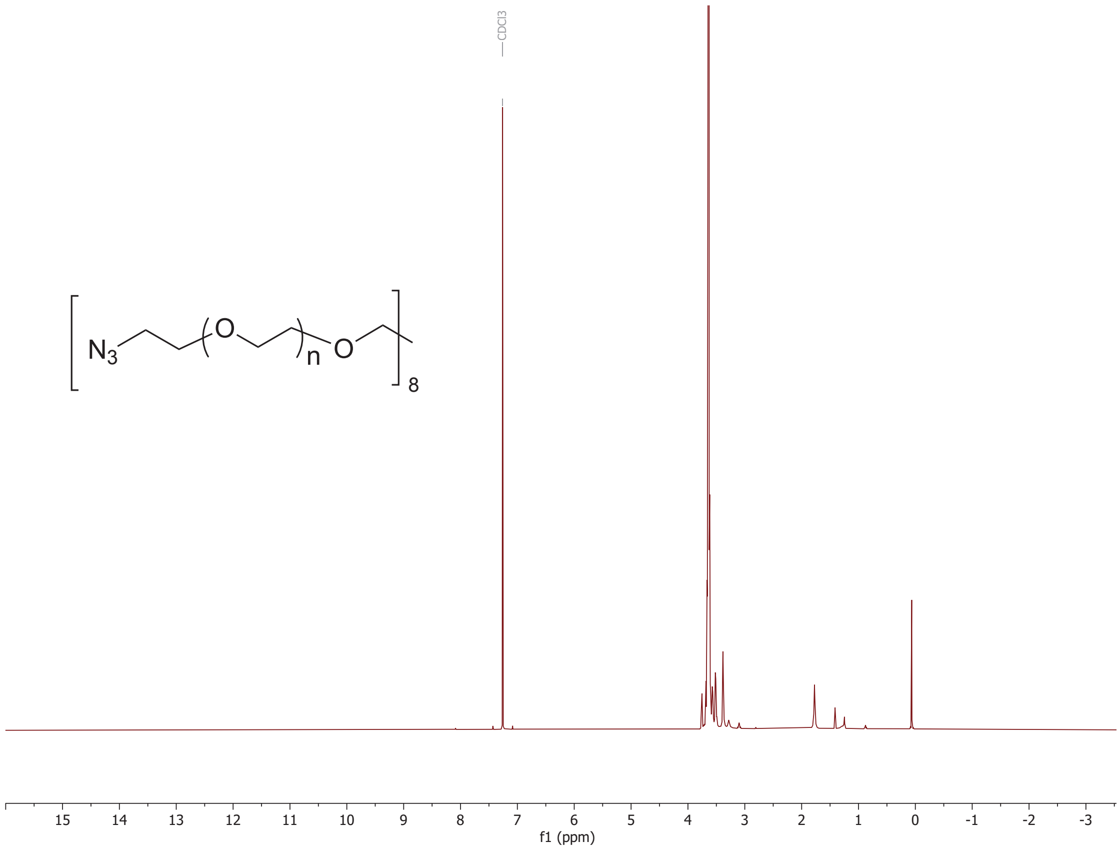

**Figure S26**. $^{1}$H NMR spectrum of azide-functionalized PEG (600 MHz, $CDCl_3$).

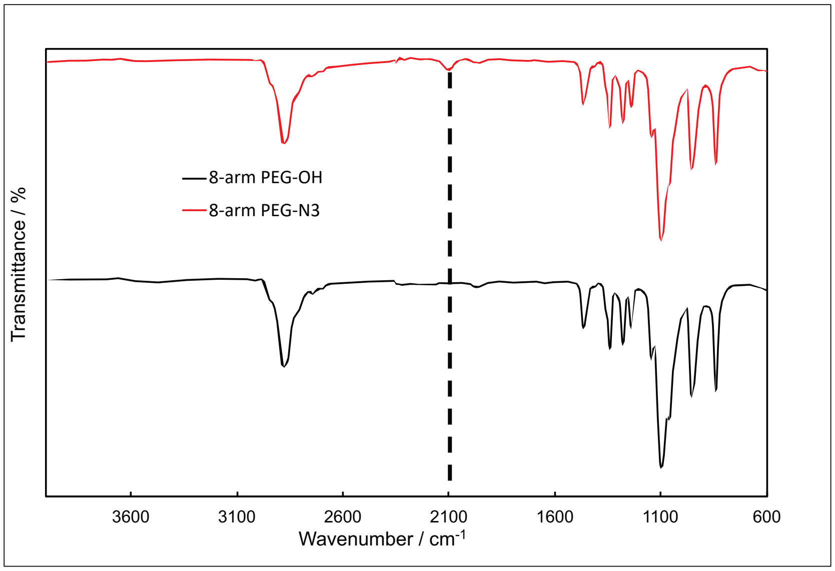

**Figure S27**. Fourier-transform infrared (FT-IR) spectrum of azide-functionalized 8-arm PEG, showing the characteristic azide ($–N_3$) absorption peak at 2098 $cm^{-1}$, confirming successful azide functionalization.

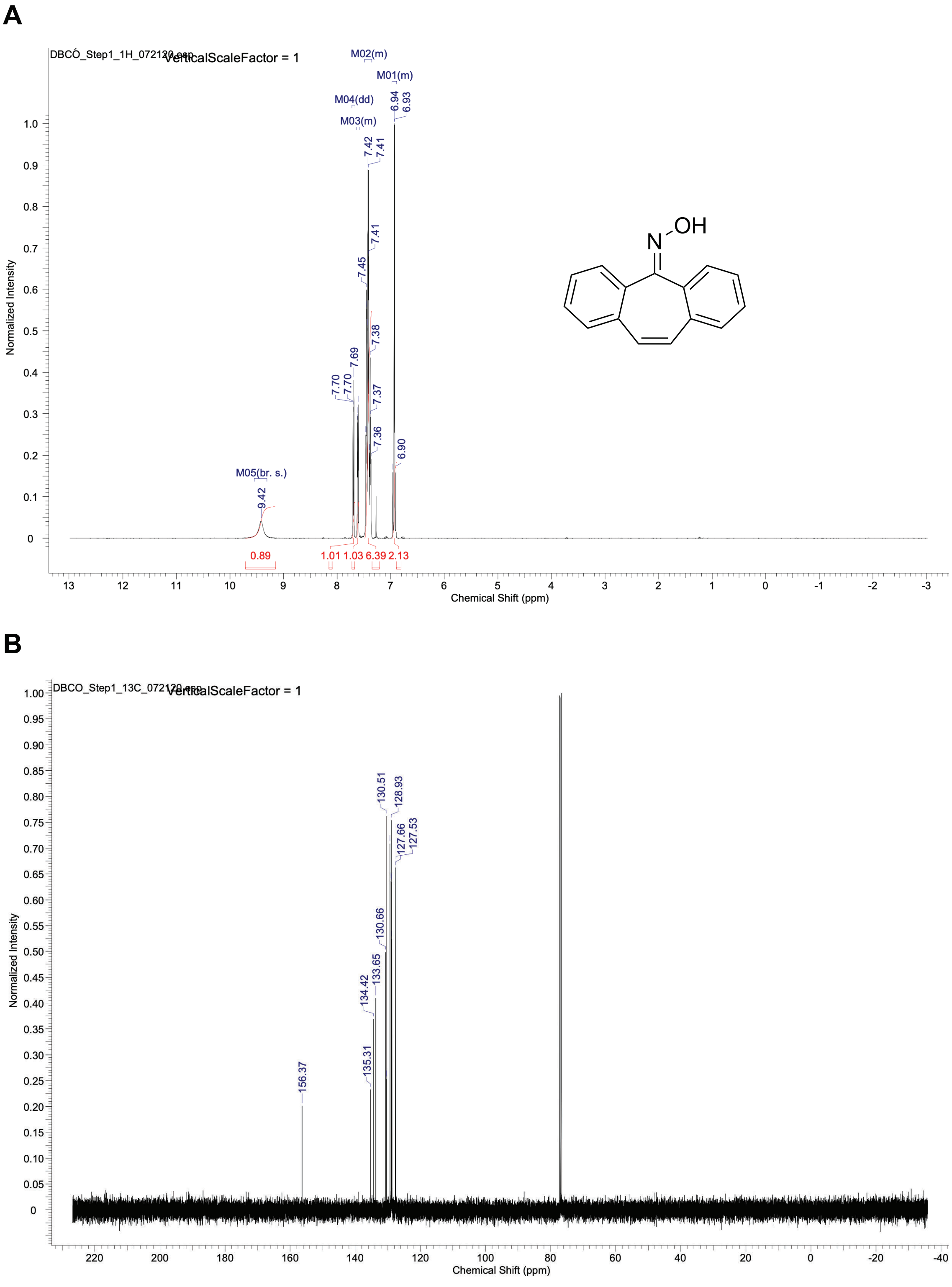

**Figure S28.** $^{1}H$ NMR (**A**) and $^{13}C$ NMR (**B**) spectra of the dibenzosuberenone oxime in $CDCl_3$.

**A**

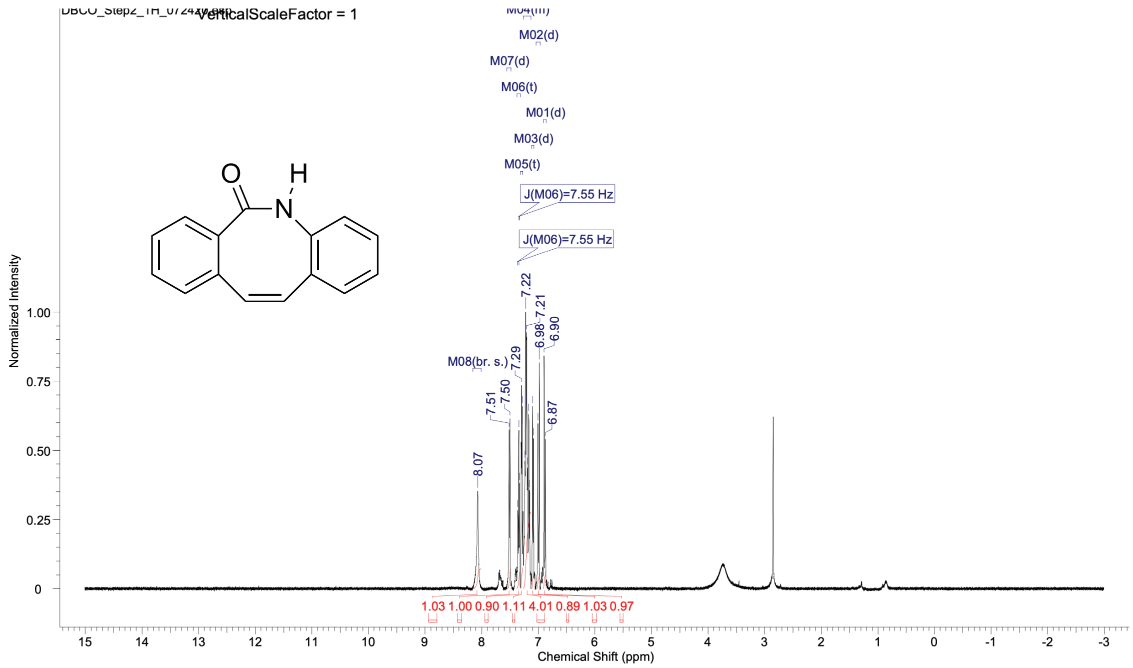

**B**

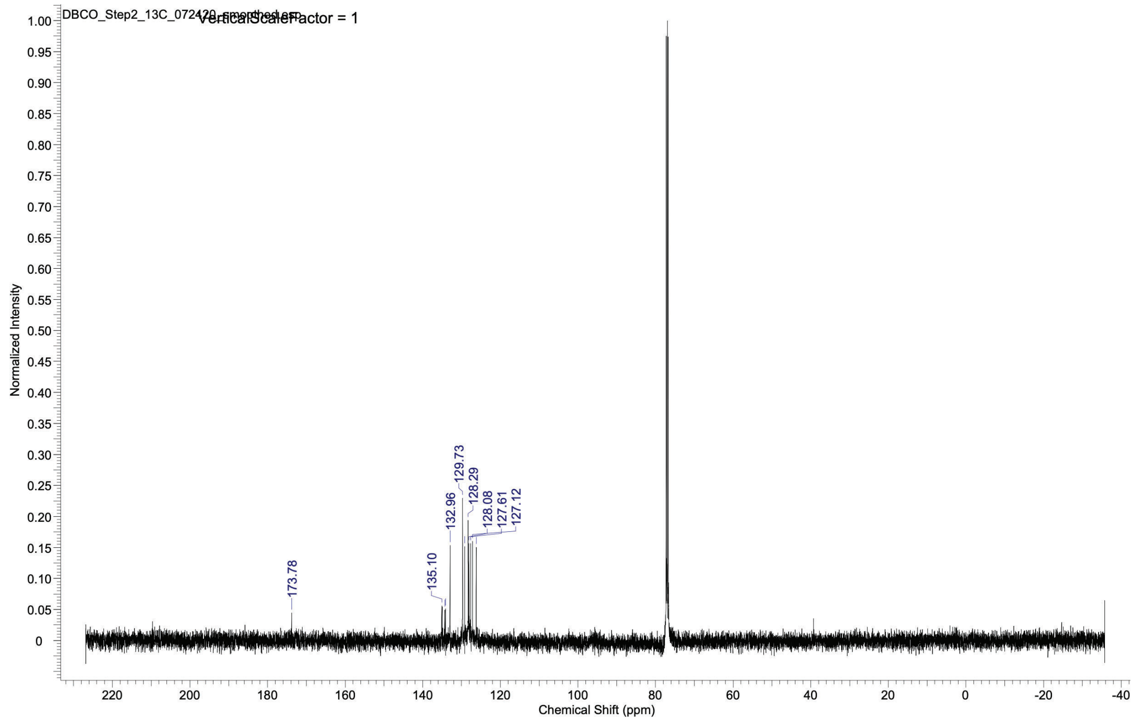

**Figure S29.** $^{1}H$ NMR (**A**) and $^{13}C$ NMR (**B**) spectra of the dibenzo[c,e]azepine *N*-oxide in $CDCl_3$.

A

B

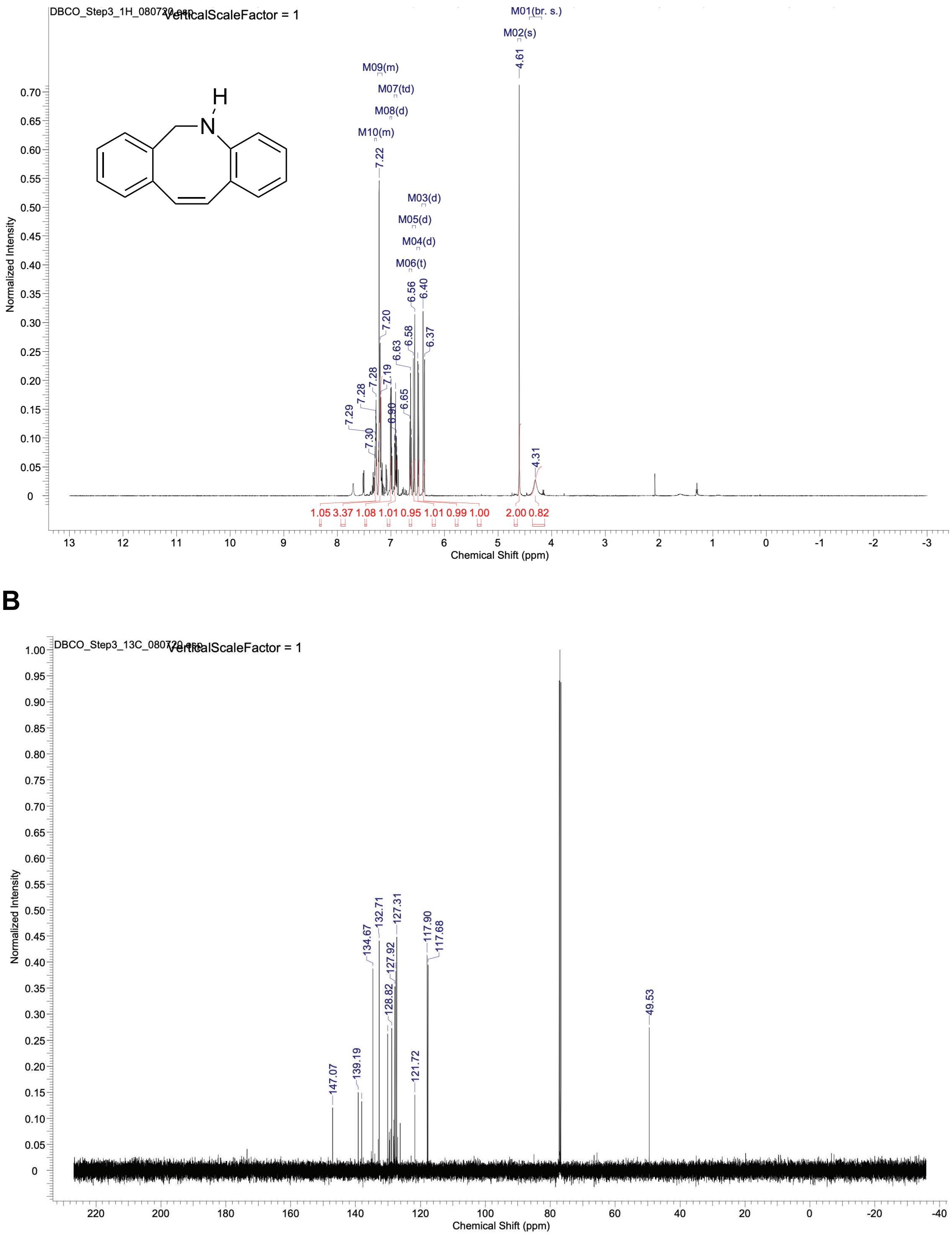

**Figure S30.** $^{1}H$ NMR (**A**) and $^{13}C$ NMR (**B**) spectra of the dibenzocyclooctene in $CDCl_3$.

**A**

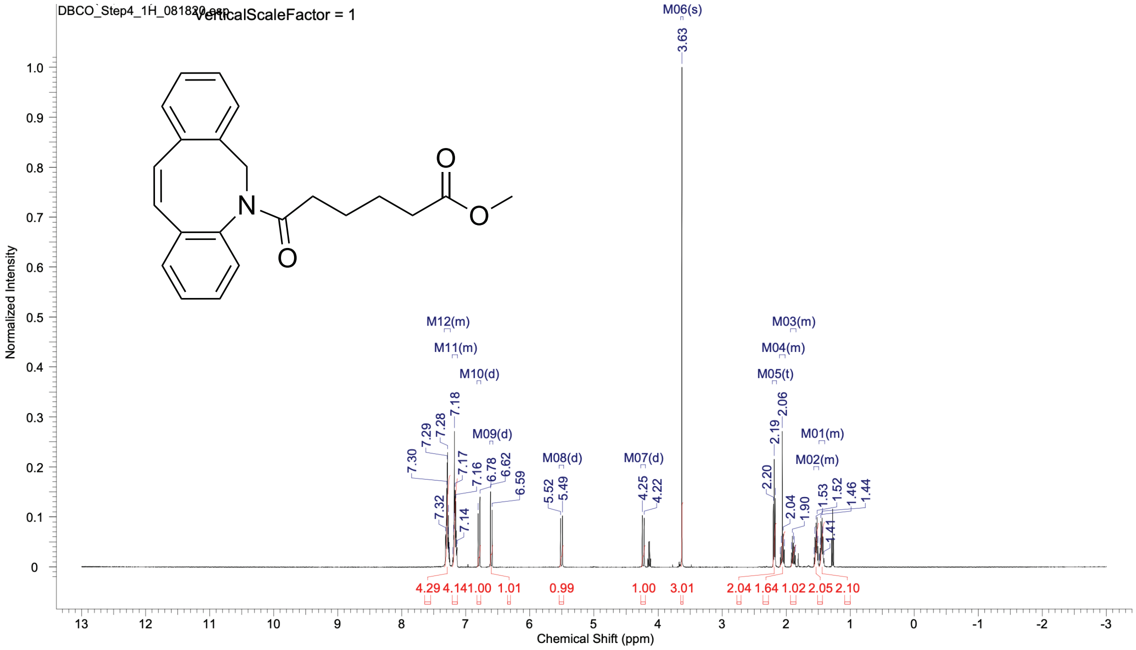

**B**

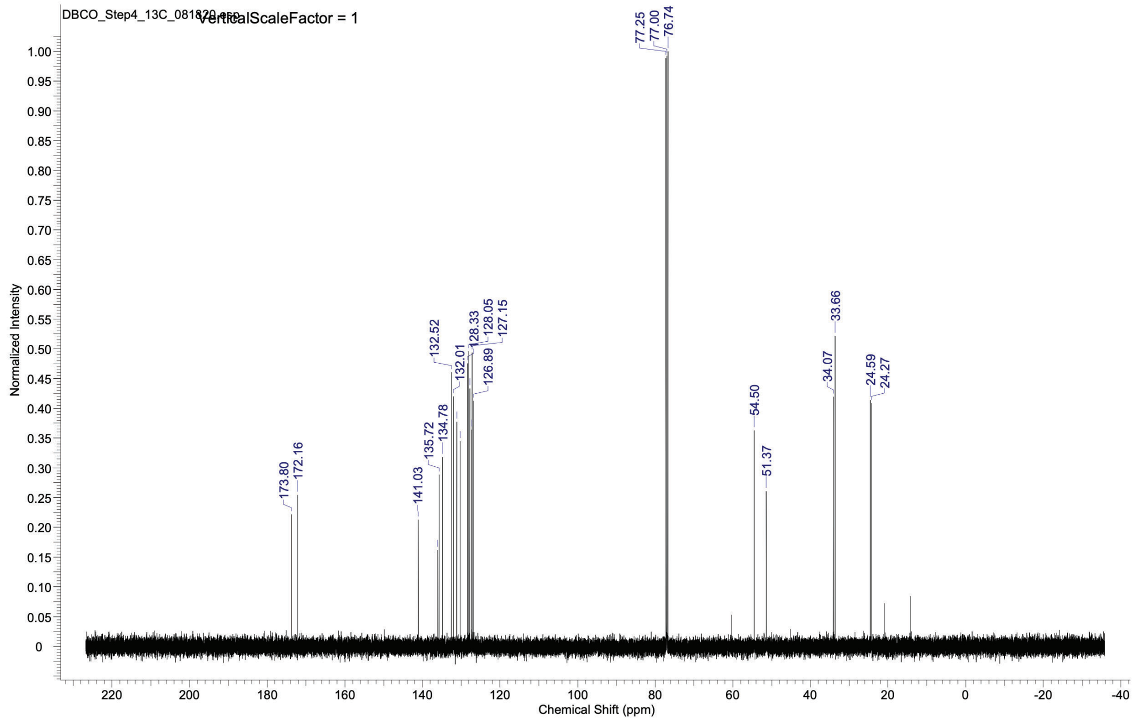

**Fig. S31.** $^{1}$H NMR (**A**) and $^{13}$C NMR (**B**) spectra of the dibenzocyclooctene-$C_6$-COOMe in $CDCl_3$.

**A**

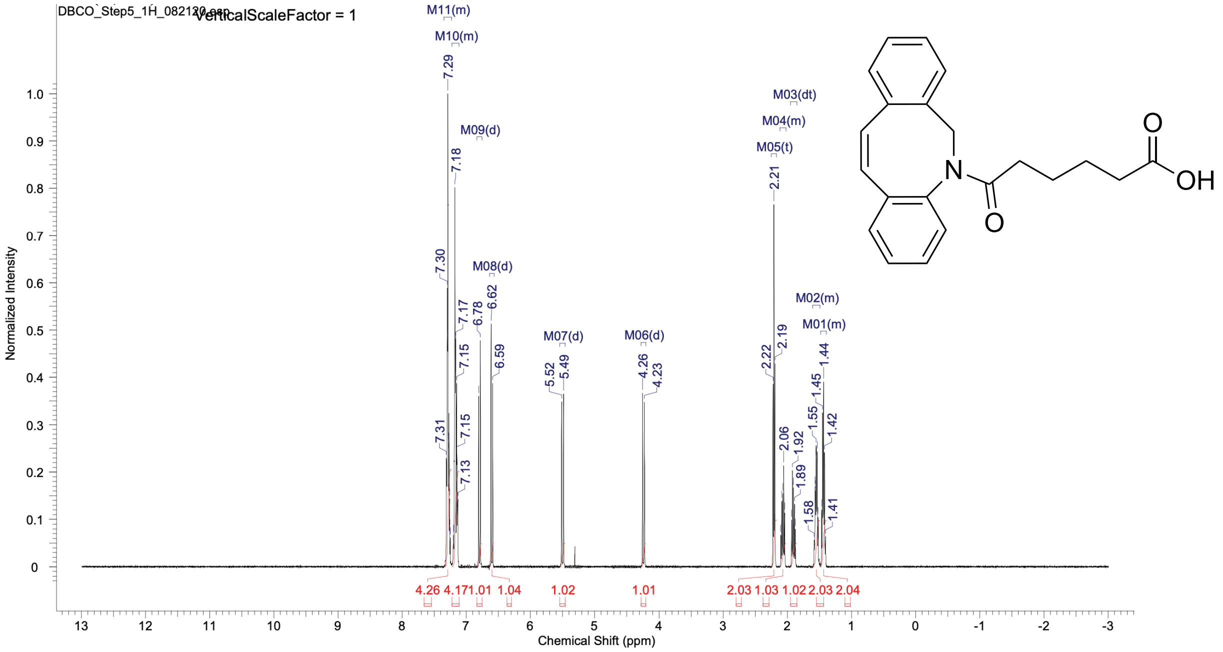

**B**

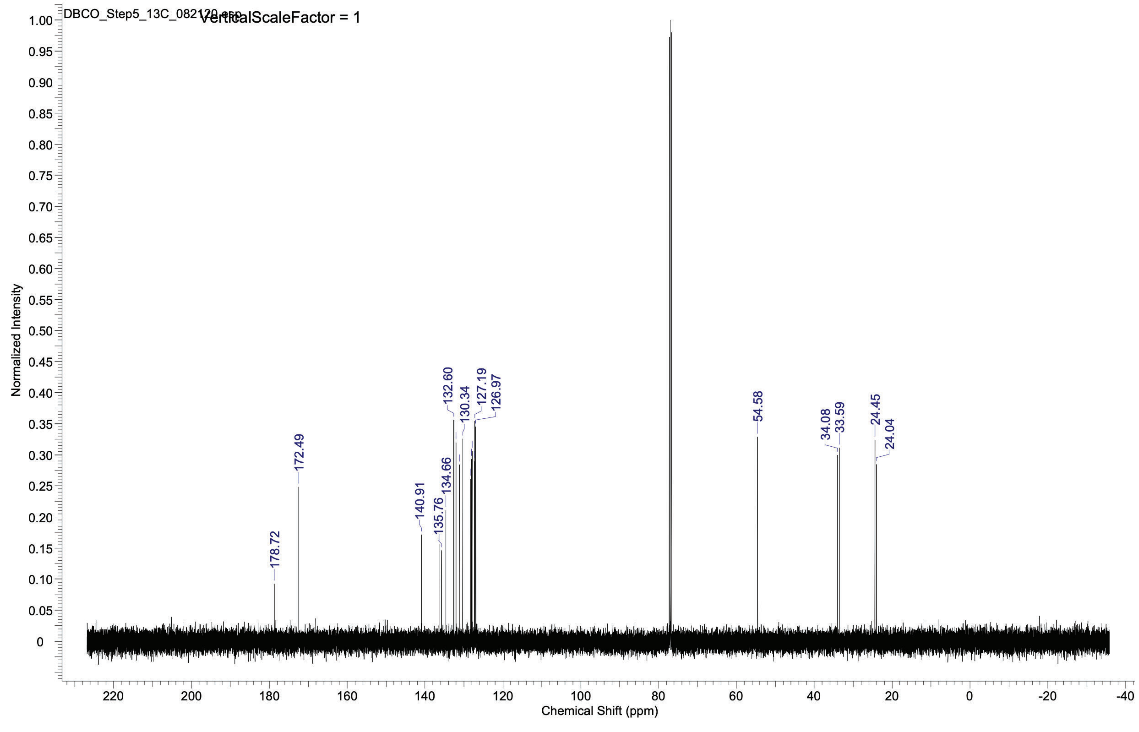

**Fig. S32.** $^{1}H$ NMR (**A**) and $^{13}C$ NMR (**B**) spectra of the dibenzocyclooctene -$C_6$-COOH in $CDCl_3$.

A

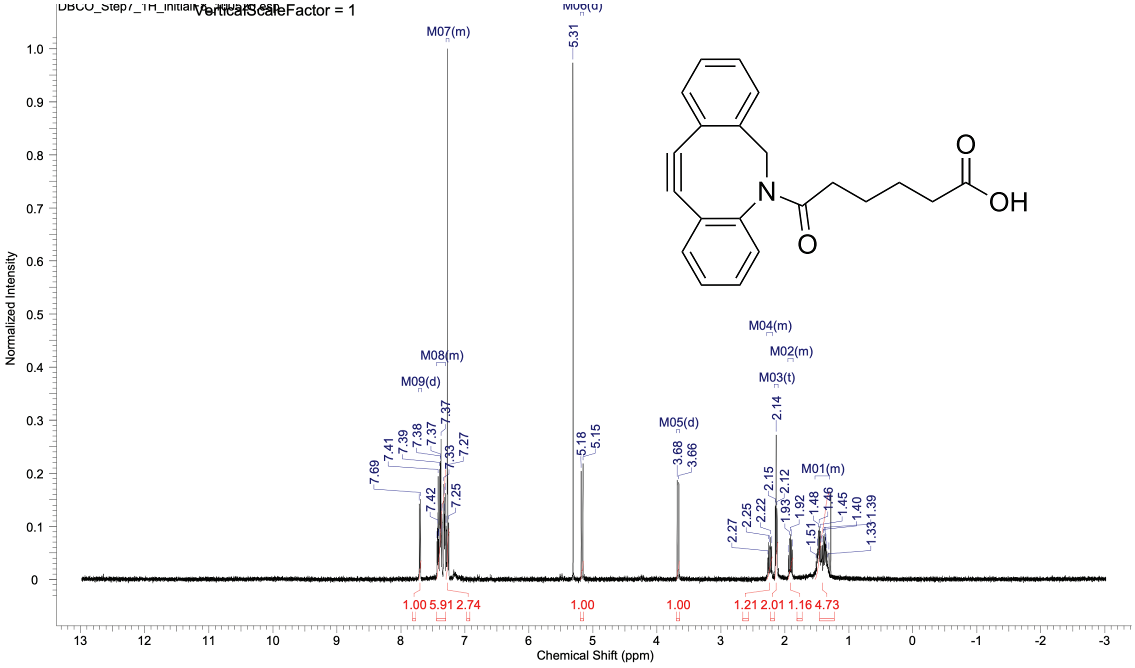

B

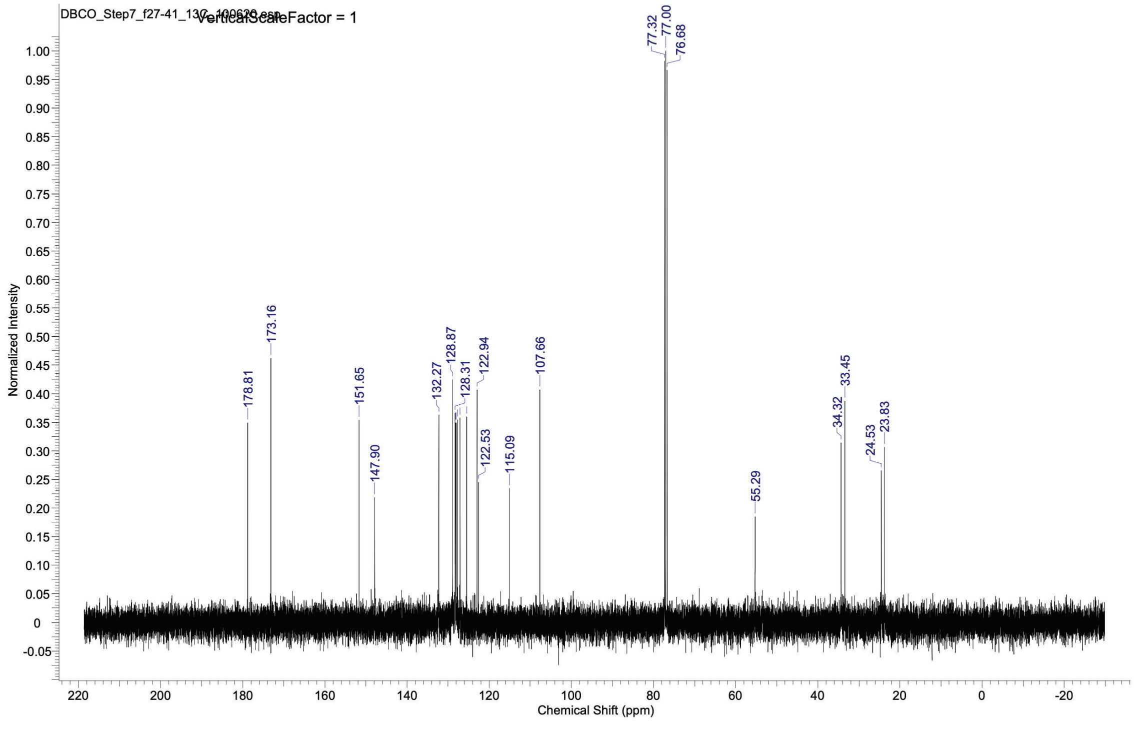

**Fig. S33.** $^{1}$H NMR (**A**) and $^{13}$C NMR (**B**) spectra of the DBCO-$C_6$-COOH in $CDCl_3$.

**A**

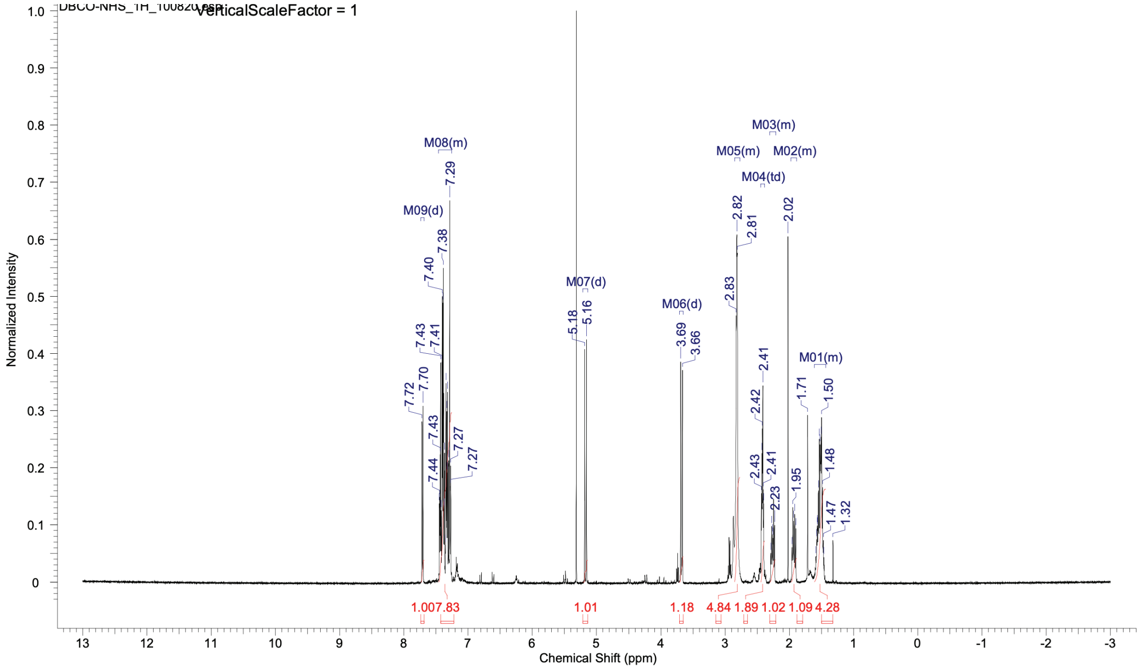

**B**

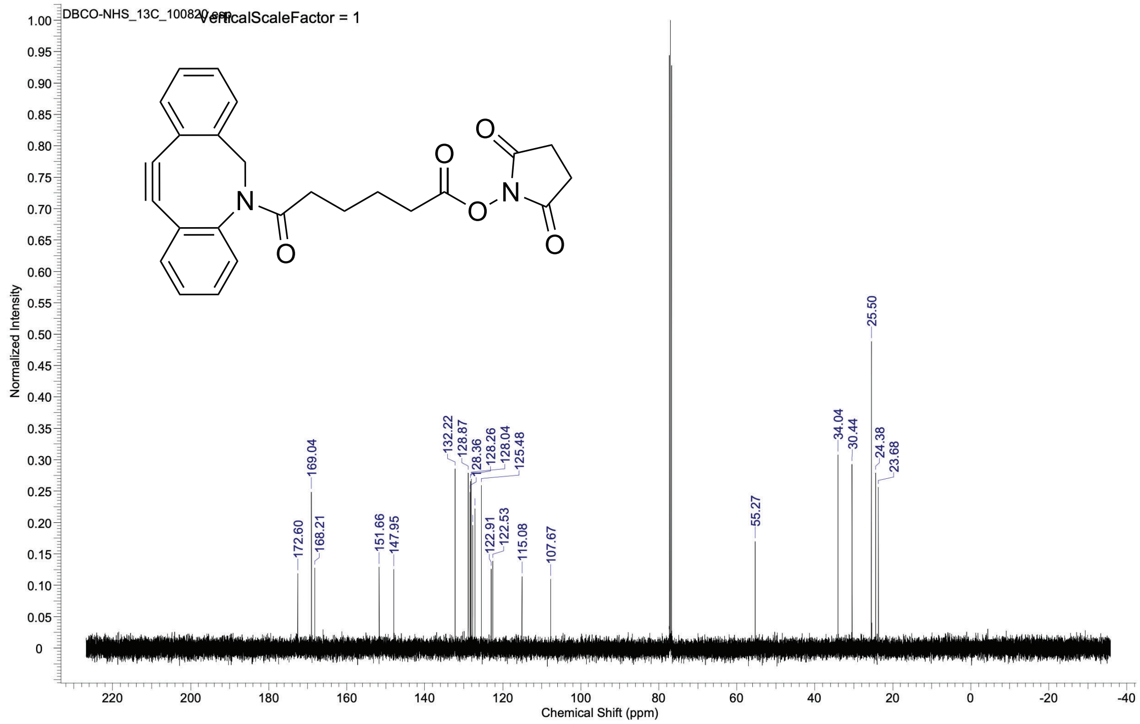

**Fig. S34.** $^{1}H$ NMR (**A**) and $^{13}C$ NMR (**B**) spectra of DBCO-$C_6$-NHS ester in $CDCl_3$.

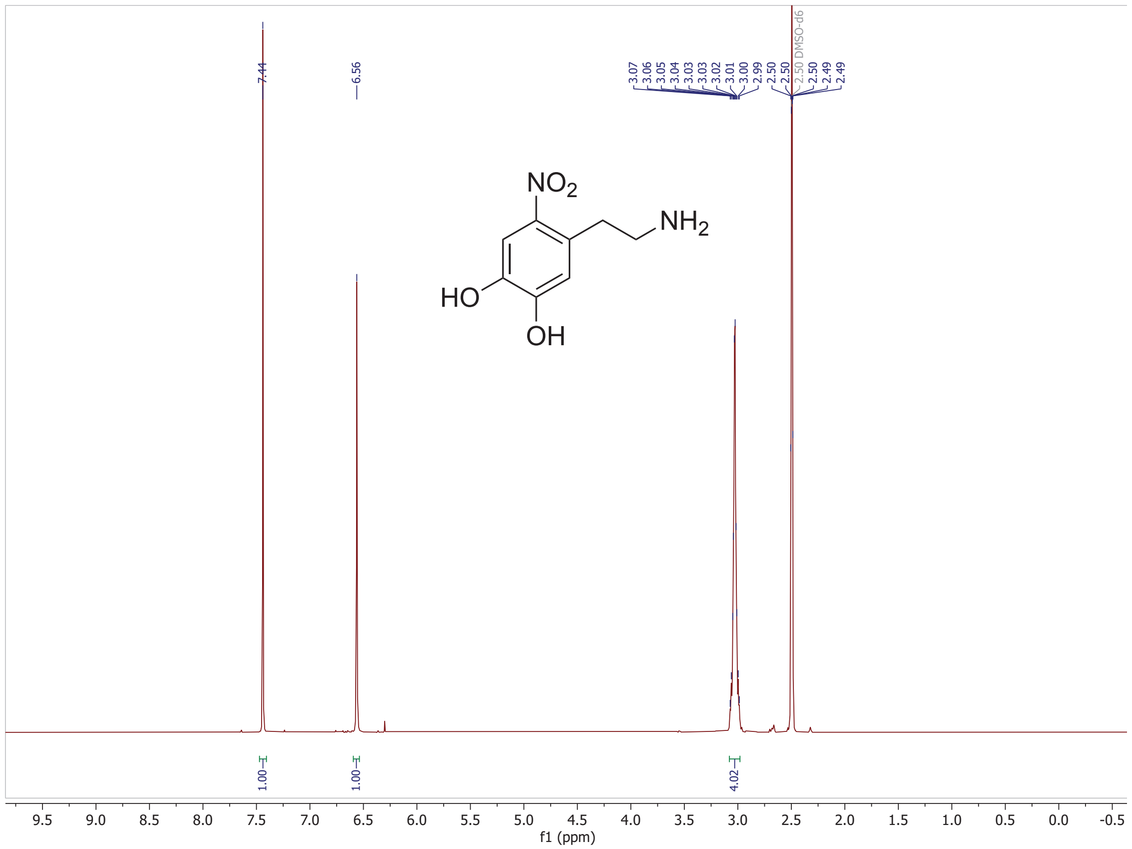

**Figure S35.** $^1$H NMR spectrum of nitrodopamine (400 MHz, DMSO).

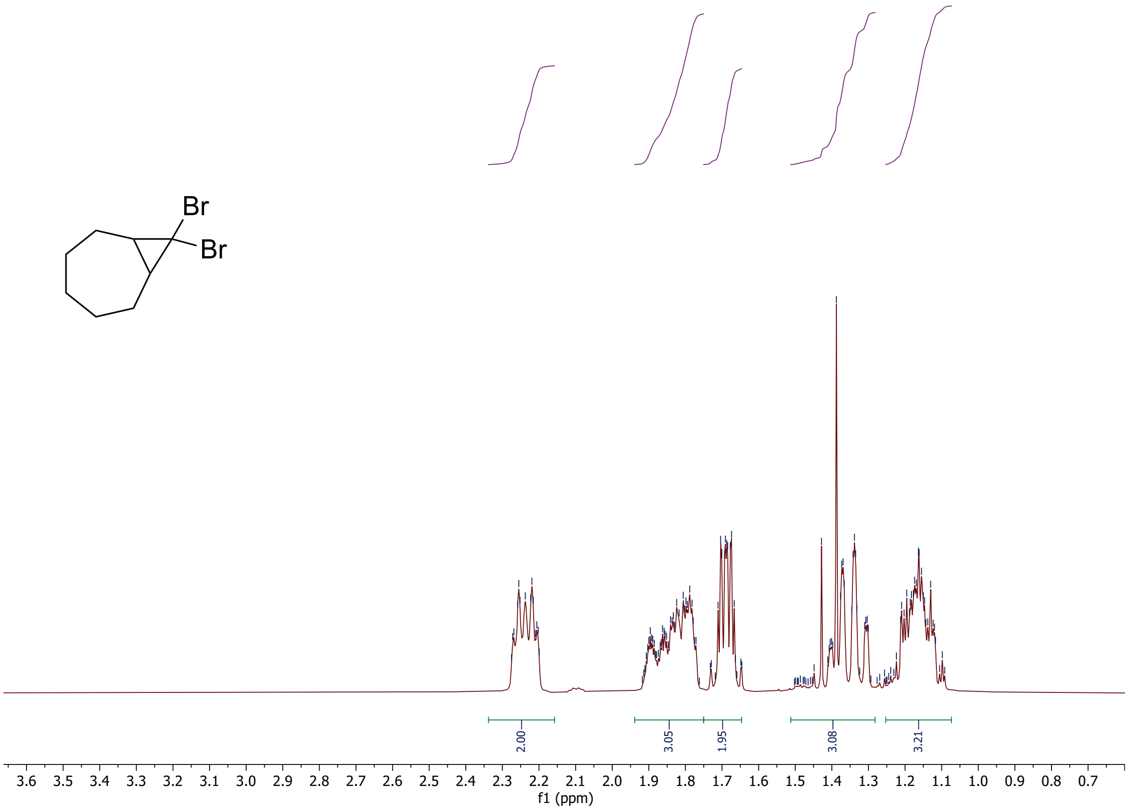

**Figure S36.** $^{1}$H NMR spectrum of 8,8-dibromobicyclo [5.1.0] octane (400 MHz, $CDCl_3$).

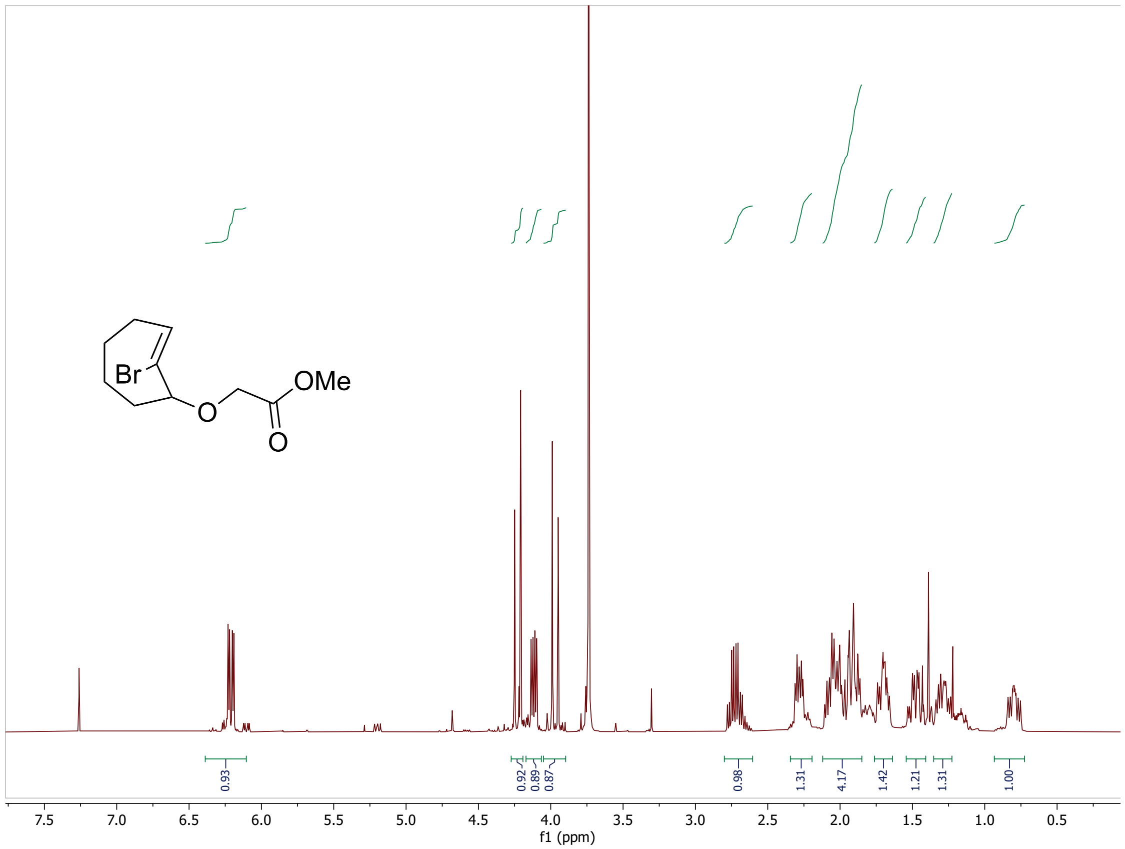

**Figure S37.** [1]H NMR spectrum of [(Z)-2-bromo-2-cycloocten-1-yloxy] acetic acid, methyl ester (400 MHz, $CDCl_3$).

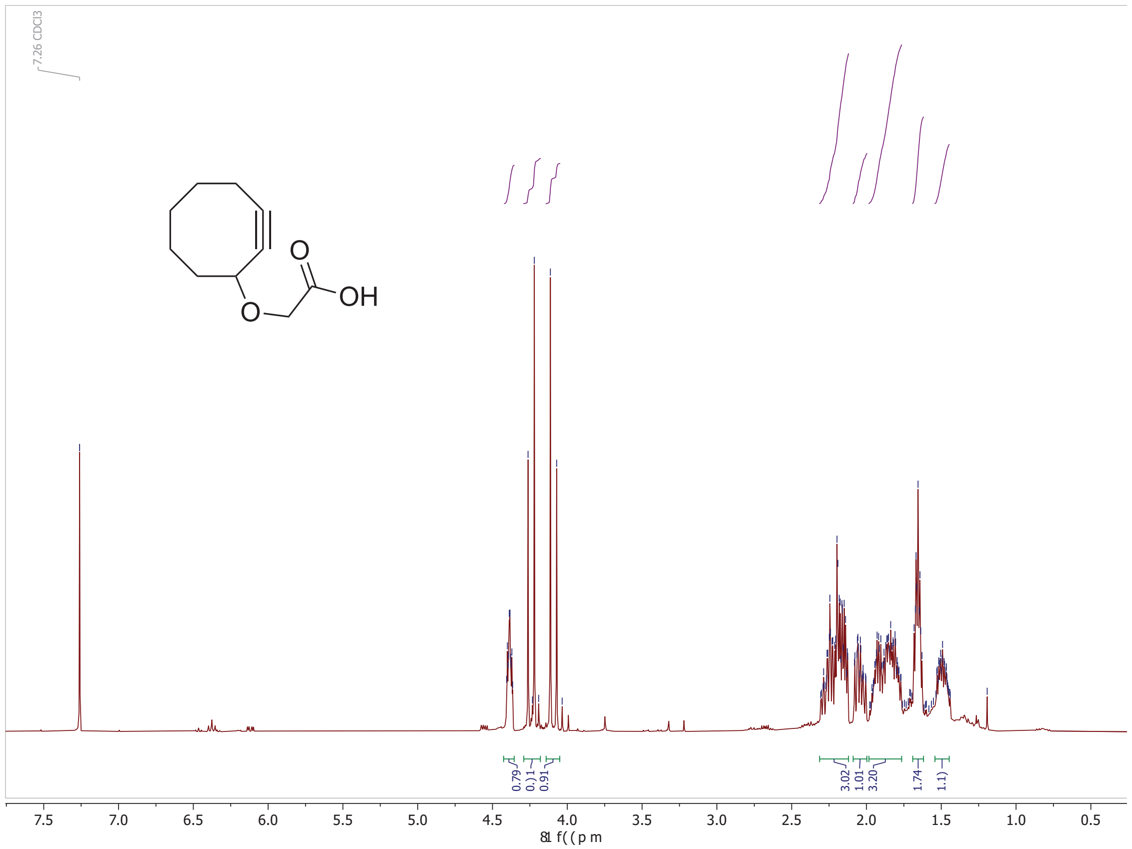

**Figure S38.** $^1$H NMR spectrum of (2-Cyclooctyn-1-yloxy) acetic Acid (400 MHz, $CDCl_3$).

Flow cytometry gating strategy for MuSC isolation

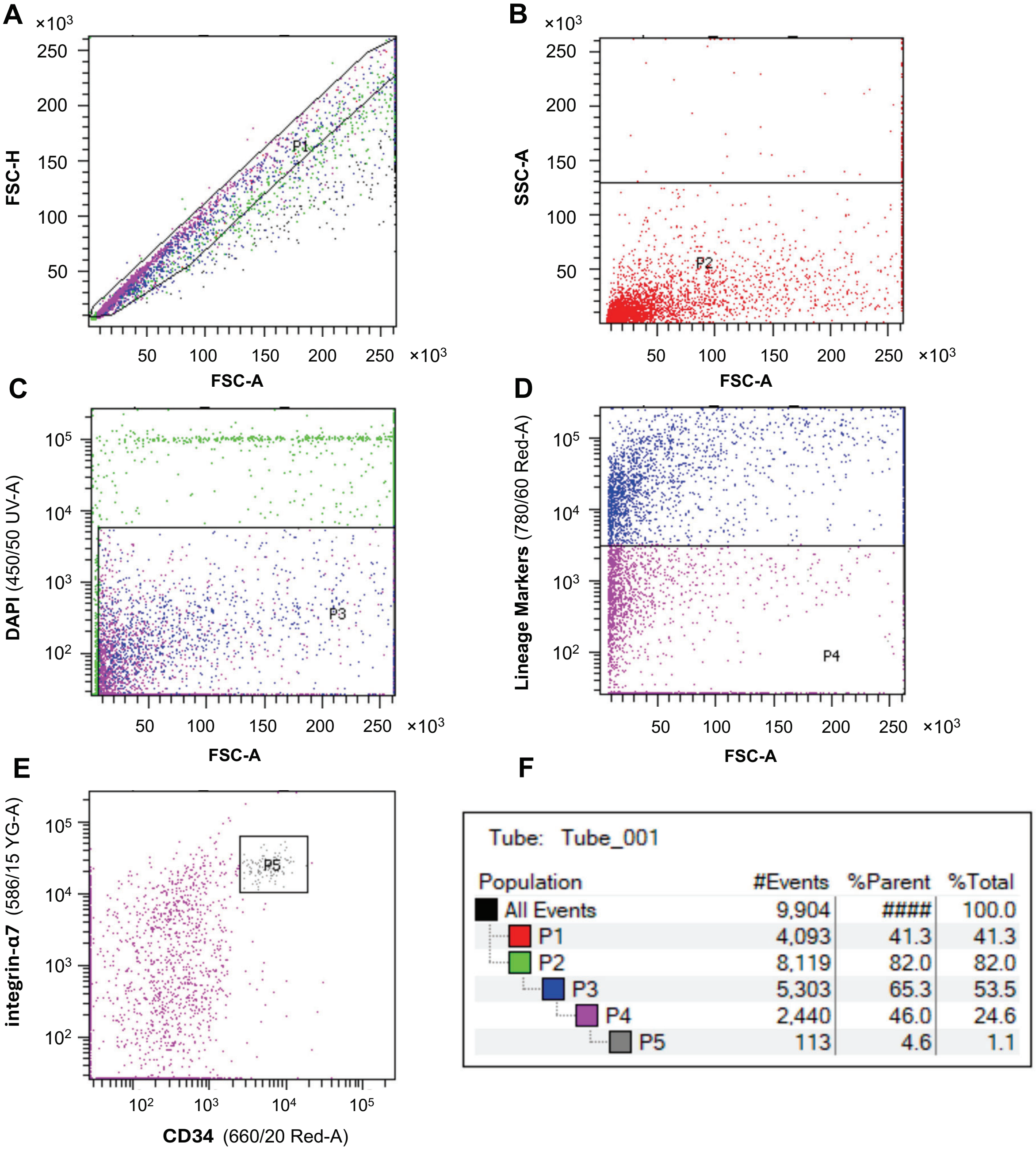

**Figure S39.** FACS gating strategy for MuSC isolation. (**A**) Doublets were excluded by gating on forward scatter area (FSC-A) versus forward scatter height (FSC-H) to select singlet cells (P1). (**B**) Debris was excluded based on forward and side scatter properties (FSC-A versus SSC-A) to obtain the intact cell population (P2). (**C**) Dead cells were excluded using DAPI staining, and DAPI-negative viable cells were selected (P3). (**D**) Hematopoietic, endothelial, and stromal cells were excluded by negative selection using a lineage marker antibody cocktail (CD45, CD11b, CD31, and Sca1) labeled with APC-Cy7 or APC-Fire750 (780/60 Red-A), yielding the lineage-negative population (P4). (**E**) MuSCs were identified and sorted as integrin-α7$^{+}$CD34$^{+}$ cells (P5). (**F**)

Representative gating statistics showing the hierarchy, number of events, and the percentage of cells retained at each gating step.

**Supporting Tables**

**Table S1.** Soft, stress relaxing hydrogel formulations.

| Type | Category | Soft, $\tau_{1/2} \sim 10^2$ s | Soft, $\tau_{1/2} \sim 10^3$ s |
|---|---|---|---|
| PEG-hyd | Concentration | 4.5% | 4% |
| | Architecture | 4-arm, 10kDa | 4-arm, 10kDa |
| PEG-ald | Concentration | 6% | 2.42% |
| | Architecture | 4-arm, 20kDa | 8-arm, 20kDa |
| PEG-DBCO | Concentration | 4.2% | 3.2% |
| | Architecture | 4-arm, 10kDa | 4-arm, 10kDa |
| PEG-$N_3$ | Concentration | 6.3% | 4.8% |
| | Architecture | 8-arm, 20kDa | 8-arm, 20kDa |

**Table S2.** Stiff, stress relaxing hydrogel formulations.

| Type | Category | Stiff, $\tau_{1/2} \sim 10^2$ s | Stiff, $\tau_{1/2} \sim 10^3$ s |
|---|---|---|---|
| PEG-hyd | Concentration | 6.3% | 5% |
| | Architecture | 4-arm, 10kDa | 8-arm, 20kDa |
| PEG-ald | Concentration | 8.4% | 3.3% |
| | Architecture | 4-arm, 20kDa | 8-arm, 20kDa |
| PEG-DBCO | Concentration | 3.6% | 3% |
| | Architecture | 4-arm, 10kDa | 4-arm, 10kDa |
| PEG-CO | Concentration | 2.5% | 2.7% |
| | Architecture | 4-arm, 10kDa | 4-arm, 10kDa |
| PEG-$N_3$ | Concentration | 6.3% | 6.5% |
| | Architecture | 8-arm, 20kDa | 8-arm, 20kDa |

**Table S3.** Soft, fast relaxing hydrogel formulation.

| Type | Category | Soft, $\tau_{1/2} \sim 10^{-1}$ s |
|---|---|---|
| PEG-ND | Concentration | 6% |
| | Architecture | 4-arm, 10kDa |
| PEG-CO | Concentration | 15% |
| | Architecture | 4-arm, 10kDa |
| PEG-$N_3$ | Concentration | 9% |
| | Architecture | 8-arm, 20kDa |
| (3-(Azidomethyl)phenyl)boronic acid | Concentration | 0.0011M |

**Table S4.** Elastic hydrogel formulations.

| Type | Category | Soft, Elastic | Stiff, Elastic |
|---|---|---|---|
| PEG-CO | Concentration | 5% | 8.5% |
| | Architecture | 4-arm, 10kDa | 4-arm, 10kDa |
| PEG-$N_3$ | Concentration | 8.5% | 8.5% |
| | Architecture | 8-arm, 20kDa | 8-arm, 20kDa |

**Table S5.** Antibodies for FACS enrichment of MuSCs.

| Antibody | Conjugate | Vendor | Catalog |
|---|---|---|---|
| Rat anti-mouse CD34 | AF647 | BD Biosciences | 560230 |
| Rat anti-mouse CD31 | APC-Cy7 | BioLegend | 102534 |
| Rat anti-mouse CD11b | APC-Cy7 | BioLegend | 101226 |
| Rat anti-mouse Sca-1 | APC-Fire750 | BioLegend | 122526 |
| Rat anti-mouse CD45 | APC-Cy7 | BioLegend | 103116 |
| Rat anti-mouse Alpha 7 Integrin | PE | AbLab | 53-0010 |

**Table S6.** Small molecule drugs for mechanotransduction perturbation experiments.

| Target | Type | Name | Vendor | Concentration | Time |
|---|---|---|---|---|---|
| TIAM1 | Inhibitor | NSC23766 | Caymen Chem | 150 uM | 24, 72 h |
| FAK | Inhibitor | Defactinib | Caymen Chem | 10 uM | 24, 72 h |
| Akt | Inhibitor | Akt inhibitor VIII | Caymen Chem | 37.5 uM | 24, 72 h |
| DOCK1 | Inhibitor | TBOPP | MedChemExpress | 20 uM | 24, 72 h |
| PAK1 | Inhibitor | NVS-PAK1-1 | Caymen Chem | 30 uM | 24 h |
| PTEN | Activator | Tyroserleutide hydrochloride | MedChemExpress | 100 uM | 24 h |
| RhoA | Inhibitor | Rhosin | Caymen Chem | 60 uM | 24 h |
| ROCK | Inhibitor | Y-27632 | Caymen Chem | 50 uM | 24 h |
| PTEN | Inhibitor | bpV(HOpic) | Caymen Chem | 10 uM | 24 h |
| Cdc42 | Inhibitor | ML-141 | Caymen Chem | 20 uM | 24, 72 h |
| PAK1 | Activator | Fingolimod | Caymen Chem | 3 uM | 24, 72 h |

**Table S7.** Primary antibodies and reagents for immunofluorescence.

| Antibody | Conjugate | Vendor | Catalog | Dilution |
|---|---|---|---|---|
| Goat anti-mouse IgG1 | AF647 | Jackson ImmunoResearch | 115-605-205 | 1:400 |
| Goat anti-mouse IgG2b | Cy3 | Jackson ImmunoResearch | 115-165-207 | 1:400 |
| Goat anti-mouse IgG2b | AF405 | Jackson ImmunoResearch | 115-475-207 | 1:400 |
| Goat anti-Rabbit IgG | AF750 | Invitrogen | A-21039 | 1:400 |
| Goat anti-Rabbit IgG | AF405 | Invitrogen | A48254 | 1:250 |
| Goat anti-mouse IgM | DyLight 755 | Invitrogen | SA5-10155 | 1:400 |
| Phalloidin-TRITC | - | Millipore Sigma | P1951 | 1:1000 |
| DAPI | - | Biotium | 40043 | 1:10000 |

**Table S8.** Secondary antibodies and reagents for immunofluorescence.

| Antibody | Conjugate | Vendor | Catalog | Dilution |
|---|---|---|---|---|
| Goat anti-mouse IgG1 | AF647 | Jackson ImmunoResearch | 115-605-205 | 1:400 |
| Goat anti-mouse IgG2b | Cy3 | Jackson ImmunoResearch | 115-165-207 | 1:400 |
| Goat anti-mouse IgG2b | AF405 | Jackson ImmunoResearch | 115-475-207 | 1:400 |
| Goat anti-Rabbit IgG | AF750 | Invitrogen | A-21039 | 1:400 |
| Goat anti-Rabbit IgG | AF405 | Invitrogen | A48254 | 1:250 |
| Goat anti-mouse IgM | DyLight 755 | Invitrogen | SA5-10155 | 1:400 |
| Phalloidin-TRITC | - | Millipore Sigma | P1951 | 1:1000 |
| DAPI | - | Biotium | 40043 | 1:10000 |

**Table S9.** Two-way ANOVA for Day1, EdU fraction.

| Source | sum_sq | df | F | PR(>F) |
|---|---|---|---|---|
| Stiffness | 22742.49259 | 1 | 557.5761 | 6.56E-24 |
| Stress Relaxation | 9029.917745 | 2 | 110.6929 | 2.43E-16 |
| Stiffness x Stress Relaxation | 8411.494724 | 2 | 103.112 | 7.42E-16 |
| Residual | 1509.161172 | 37 | | |

**Table S10.** Two-way ANOVA for Day1, pAkt mean intensity.

| Source | sum_sq | df | F | PR(>F) |
|---|---|---|---|---|
| Stiffness | 1.37E-05 | 1 | 65.16573 | 4.43E-10 |
| Stress Relaxation | 2.23E-05 | 2 | 53.10236 | 3.16E-12 |
| Stiffness x Stress Relaxation | 6.93E-06 | 2 | 16.52688 | 5.07E-06 |
| Residual | 8.8E-06 | 42 | | |

**Table S11.** Two-way ANOVA for Day1, Rac1/Cdc42 mean intensity.

| Source | sum_sq | df | F | PR(>F) |
|---|---|---|---|---|
| Stiffness | 7.54E-06 | 1 | 36.38461 | 1.28E-06 |
| Stress Relaxation | 4.14E-06 | 2 | 9.993898 | 0.000472 |
| Stiffness x Stress Relaxation | 2.62E-06 | 2 | 6.317327 | 0.005133 |
| Residual | 6.21E-06 | 30 | | |

**Table S12.** Two-way ANOVA for Day3, pAkt mean intensity.

| Source | sum_sq | df | F | PR(>F) |
|---|---|---|---|---|
| Stiffness | 1.33E-05 | 1 | 41.04854 | 1.03E-07 |
| Stress Relaxation | 2.58E-05 | 2 | 39.72502 | 2.07E-10 |
| Stiffness x Stress Relaxation | 6.05E-06 | 2 | 9.325281 | 0.000445 |
| Residual | 1.36E-05 | 42 | | |

**Table S13.** Two-way ANOVA for Day3, EZH2 mean intensity.

| Source | sum_sq | df | F | PR(>F) |
|---|---|---|---|---|
| Stiffness | 8.76E-05 | 1 | 137.6128 | 4.98E-14 |
| Stress Relaxation | 0.000185 | 2 | 145.2382 | 3.03E-18 |
| Stiffness x Stress Relaxation | 5.55E-05 | 2 | 43.54652 | 1.89E-10 |
| Residual | 2.36E-05 | 37 | | |

**Table S14.** Two-way ANOVA for Day7, EdU fraction.

| Source | sum_sq | df | F | PR(>F) |
|---|---|---|---|---|
| Stiffness | 2010.685 | 1 | 67.71429 | 4.7E-10 |
| Stress Relaxation | 1869.967 | 2 | 31.48764 | 7.25E-09 |
| Stiffness x Stress Relaxation | 427.7522 | 2 | 7.202753 | 0.002177 |
| Residual | 1158.053 | 39 | | |

**Table S15.** Two-way ANOVA for Day7, Pax7 fraction.

| Source | sum_sq | df | F | PR(>F) |
|---|---|---|---|---|
| Stiffness | 113.5068 | 1 | 1.415323 | 0.239194 |
| Stress Relaxation | 1459.544 | 2 | 9.099563 | 0.000379 |
| Stiffness x Stress Relaxation | 44.31705 | 2 | 0.276296 | 0.759617 |
| Residual | 4491.119 | 56 | | |

**Table S16.** Two-way ANOVA for Day7, Acta1 fraction.

| Source | sum_sq | df | F | PR(>F) |
|---|---|---|---|---|
| Stiffness | 1065.881 | 1 | 23.95388 | 6.83E-06 |
| Stress Relaxation | 620.8819 | 2 | 6.976639 | 0.0018 |
| Stiffness x Stress Relaxation | 130.7372 | 2 | 1.469049 | 0.237683 |
| Residual | 2892.318 | 65 | | |

### References

1. T. E. Brown, J. S. Silver, B. T. Worrell, I. A. Marozas, F. M. Yavitt, K. A. Günay, C. N. Bowman, K. S. Anseth, Secondary Photocrosslinking of Click Hydrogels To Probe Myoblast Mechanotransduction in Three Dimensions. *Journal American Chemical Society* **140**, 11585–11588 (2018).

2. S. A. McNelles, J. L. Pantaleo, A. Adronov, Highly Efficient Multigram Synthesis of Dibenzoazacyclooctyne (DBCO) without Chromatography. *Org. Process Res. Dev.* **23**, 2740–2745 (2019).

3. M. Burk, S. Rothstein, P. Dubé, Enabling the Multigram Synthesis of (2-Cyclooctyn-1-yloxy)acetic Acid. *Org. Process Res. Dev.* **22**, 108–110 (2018).

4. S. Tang, H. Ma, H. Tu, H. Wang, P. Lin, K. S. Anseth, Adaptable Fast Relaxing Boronate-Based Hydrogels for Probing Cell–Matrix Interactions. *Adv Sci (Weinh)* **5**, 1800638 (2018).

**Movies**

**Movie S1. Live-cell imaging of MuSC expansion on soft, $\tau_{1/2}$ ~ $10^2$ s hydrogel.** Live-cell imaging of freshly isolated wild-type MuSCs from 1-month-old mice cultured on soft hydrogel with a stress-relaxation half-time ($\tau_{1/2}$) of ~$10^2$ s. Imaging was initiated approximately 20 h after cell seeding and continued for ~72 h post-seeding, with images acquired at 33-min intervals. Scale bar: 50 um.

**Movie S2. Live-cell imaging of MuSC expansion on soft, $\tau_{1/2}$ ~ $10^3$ s hydrogel.** Live-cell imaging of freshly isolated wild-type MuSCs from 1-month-old mice cultured on soft hydrogel with a stress-relaxation half-time ($\tau_{1/2}$) of ~$10^3$ s. Imaging was initiated approximately 20 h after cell seeding and continued for ~72 h post-seeding, with images acquired at 33-min intervals. Scale bar: 50 um.

**Movie S3. Live-cell imaging of MuSC expansion on soft, elastic hydrogel.** Live-cell imaging of freshly isolated wild-type MuSCs from 1-month-old mice cultured on soft, elastic hydrogels. Imaging was initiated approximately 20 h after cell seeding and continued for ~72 h post-seeding, with images acquired at 33-min intervals. Scale bar: 50 um.
